# Epigenetic regulator genes direct lineage switching in *MLL-AF4* leukaemia

**DOI:** 10.1101/2021.07.16.452676

**Authors:** Ricky Tirtakusuma, Katarzyna Szoltysek, Paul Milne, Vasily V Grinev, Anetta Ptasinska, Claus Meyer, Sirintra Nakjang, Jayne Y Hehir-Kwa, Daniel Williamson, Pierre Cauchy, Salam A Assi, Maria R Imperato, Fotini Vogiatzi, Shan Lin, Mark Wunderlich, Janine Stutterheim, Alexander Komkov, Elena Zerkalenkova, Paul Evans, Hesta McNeill, Alex Elder, Natalia Martinez-Soria, Sarah E Fordham, Yuzhe Shi, Lisa J Russell, Deepali Pal, Alex Smith, Zoya Kingsbury, Jennifer Becq, Cornelia Eckert, Oskar A Haas, Peter Carey, Simon Bailey, Roderick Skinner, Natalia Miakova, Matthew Collin, Venetia Bigley, Muzlifah Haniffa, Rolf Marschalek, Christine J Harrison, Catherine A Cargo, Denis Schewe, Yulia Olshanskaya, Michael J Thirman, Peter N Cockerill, James C Mulloy, Helen J Blair, Josef Vormoor, James M Allan, Constanze Bonifer, Olaf Heidenreich, Simon Bomken

## Abstract

The fusion gene *MLL-AF4* defines a high-risk subtype of pro-B acute lymphoblastic leukaemia. However, relapse can be associated with a switch from acute lymphoblastic to acute myeloid leukaemia. Here we show that these myeloid relapses share oncogene fusion breakpoints with their matched lymphoid presentations and can originate in either early, multipotent progenitors or committed B-cell precursors. Lineage switching is linked to substantial changes in chromatin accessibility and rewiring of transcriptional programmes indicating that the execution and maintenance of lymphoid lineage differentiation is impaired. We show that this subversion is recurrently associated with the dysregulation of repressive chromatin modifiers, notably the nucleosome remodelling and deacetylation complex, NuRD. In addition to mutations, we show differential expression or alternative splicing of NuRD members and other genes is able to reprogram the B lymphoid into a myeloid gene regulatory network. Lineage switching in *MLL-AF4* leukaemia is therefore driven and maintained by defunct epigenetic regulation.

**Statement of Significance:** We demonstrate diverse cellular origins of lineage switched relapse within *MLL*-*AF4* pro-B acute leukaemia. Irrespective of the developmental origin of relapse, dysregulation of NuRD and/or other epigenetic machinery underpins fundamental lineage reprogramming with profound implications for the increasing use of epitope directed therapies in this high-risk leukaemia.

## Introduction

Translocation of Mixed Lineage Leukaemia (*MLL*) with one of over 130 alternative partner genes is a recurrent cytogenetic finding in both acute myeloid and lymphoblastic leukaemias and is generally associated with poor prognosis (1, 2). Amongst the most common translocations is t(4;11)(q21;q23), forming the *MLL-AF4* (also known as *KMT2A-AFF1*) fusion gene. Uniquely amongst *MLL* rearrangements (*MLL*r), *MLL-AF4* is almost exclusively associated with pro-B cell acute lymphoblastic leukaemia and is prototypical of infant acute lymphoblastic leukaemia (ALL) where it carries a very poor prognosis (2). However, despite this general lymphoid presentation, *MLL-AF4* leukaemias have an intriguing characteristic - that of lineage switched relapses. Lineage switch acute leukaemias (LSALs) lose their lymphoid specific features and gain myeloid phenotype upon relapse (3-5). Alternatively, *MLL-AF4* leukaemias may harbour distinct lymphoid and myeloid populations at the same time, thus classifying as mixed phenotype acute leukaemias (MPALs) of the bilineage subtype.

In order to understand the molecular basis of lineage promiscuity and switching, we examined a unique cohort of *MLL-AF4*-positive LSAL presentation/relapse pairs and MPALs. We demonstrate that disruption of the epigenetic machinery, including the nucleosome remodelling and deacetylation complex (NuRD), is associated with the loss of lymphoid restriction. Lineage switch is then enacted through redistribution of transcription factor binding and chromatin reorganisation. Whilst identified here within this rare clinical context, our findings bare relevance for our understanding of the transforming capacity of *MLL-AF4*, and how this oncoprotein imposes lineage determination on haematopoietic precursor cells. Furthermore, given the high-risk nature of this disease, we provide a novel insight into factors which may prove critical to the effective implementation of lineage specific, epitope-directed therapies such as chimeric antigen receptor T-cell (CAR-T) cell or bi-specific T-cell engaging antibody (BiTE) approaches.

## Results

### Characterisation of MLL-AF4 acute leukaemias with lineage switch

To characterize lineage promiscuity in *MLL-AF4* leukaemia and the underlying molecular mechanisms, we collected a cohort of ten cases of *MLL-AF4* ALL comprising 6 infant, 2 paediatric and 2 adult patients who had relapsed with acute myeloid leukaemia (AML). Amongst these, one infant patient (LS10) had relapsed following B-lineage directed blinatumomab treatment (Table S1). The time to relapse ranged from 3 to 48 months. Seven patients within the cohort subsequently died. Lineage switch was defined as loss of expression of B lymphoid antigens (CD19, CD22, CD79A) with concomitant gain of expression of myeloid antigens (CD33, CD117/KIT, CD64/FCGR1A) and/or an unequivocal change in morphology to AML (Figure 1A and Table S1). In addition, we studied two *MLL-AF4* infant mixed phenotype acute leukaemias (MPALs), comprising distinct lymphoid and myeloid populations (MPAL1, MPAL2; Table S1).

**Figure 1.**
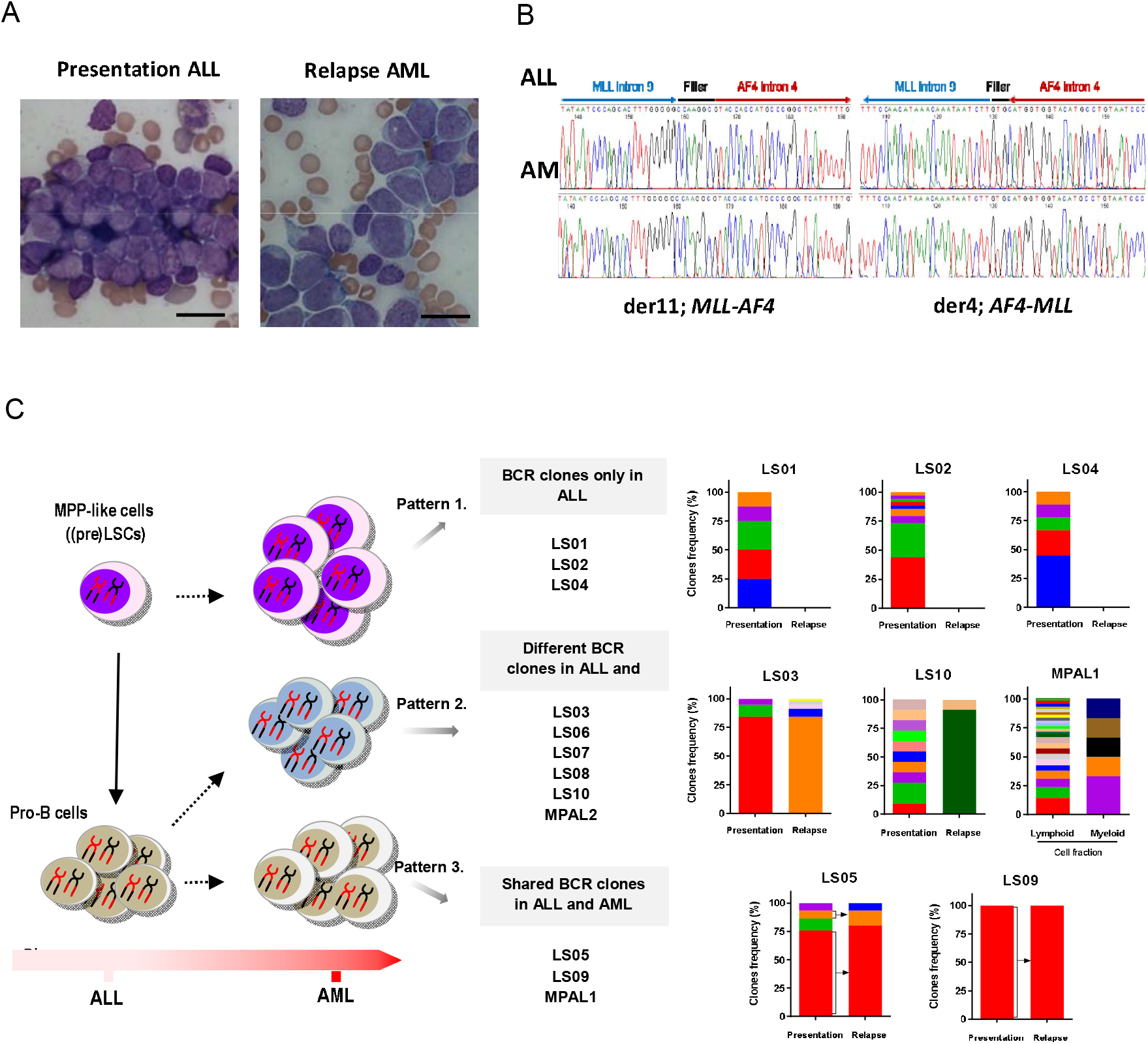
Characterisation of the *MLL-AF4* lineage switch cases. (A) Morphological change from lymphoblastic leukaemia (left panel) to acute monoblastic/monocytic leukaemia (right panel). The scale bar represents 20 μm. (B) Sanger sequencing of *MLL-AF4* and reciprocal *AF4*/*MLL* fusions in LS01 presentation ALL (upper panel) and relapse AML (lower panel) identifies a common breakpoint with identical filler sequence in ALL and AML samples. (C) Evaluation of B-cell receptor repertoires in ALL (presentation) and AML (relapse) lineage switch, and MPAL cases identified three distinct patterns. Pattern 1 - with clonotypes identified only in the ALL (LS01, LS02, LS04). Pattern 2 – distinct clonotypes identified in ALL and AML (LS03, LS06, LS07, LS08, LS10, MPAL2). Pattern 3 – identical clonotypes shared between ALL and AML (LS05, LS09, MPAL1). Graphs represent frequency of clones identified with WES and/or RNAseq in LSAL and MPAL samples, respectively.

All matched samples displayed identical oncogene fusion breakpoints at diagnosis and relapse as shown by DNA (n=14) and/or RNA (n=10) sequencing, confirming a common clonal origin and proving that the relapses are not *de novo* or therapy-associated AMLs (Figures 1B, S1A, S1B, Table S1). Breakpoints of LSALs and MPALs show a similar distribution as *MLL-AF4* ALL cases, clustering in *MLL* introns 9-11 and AF4 introns 3 and 4 (6, 7) (Figure S1C, Table S1) thus excluding that distinct, “non-canonical” chromosomal breakpoints are causative for *MLL-AF4*-positive AML. These data raised the question of the cellular origin of relapse and the nature of events secondary to *MLL-AF4* that affect lineage commitment.

### Cellular origin of lineage switched relapse

We hypothesised that myeloid leukaemias may not have undergone substantial B-cell receptor (BCR) rearrangements. We used this feature to further interrogate the developmental stage at which the relapse arose. To that end we examined BCR rearrangements within RNA-seq and whole exome-seq (WES) derived data, using MiXCR software (8). All ALL cases showed classical oligoclonal rearrangements of BCR loci, supporting the lymphoid lineage decision (Figure S2A, Table S2). However, we observed three distinct patterns for AML relapses (Figure 1C). Pattern 1 comprises AML relapse cells with no BCR rearrangements implying presence of a relapse-initiating cell residing in a primitive precursor population prior to early DJ recombination (Figure 1C, cases LS01, LS02, LS04). As a second pattern, we found unrelated BCR rearrangements, which may indicate either aberrant rearrangement in a myeloid cell or relapse initiating from either a B-lymphoid cell committed to undergo rearrangement, or a transdifferentiated minor ALL clone with an alternative rearrangement (Figure 1C, cases LS03, LS06, LS07, LS08, MPAL2). Interestingly, this pattern is also found in a relapse after blinatumomab treatment (LS10). Pattern 3 comprises shared BCR rearrangements between diagnostic and relapse material, which suggest a direct transdifferentiated myeloid relapse from the ALL (Figure 1C, cases LS05, LS09, MPAL1). These data demonstrate that AML relapses can originate from different stages of lymphoid leukaemogenesis and suggest, at least for a subset, a common precursor preceding the pro-B cell stage.

To functionally assess the plasticity of immunophenotypically defined diagnostic ALL and relapsed AML, we transplanted NSG or MISTRG mice with either ALL or AML cells from patient LS01 (Figure S2B). Because of the expression of several human myeloid growth factors, the MISTRG strain more strongly supports AML engraftment than the NSG strain, which shows a stronger lymphoid bias. Diagnostic ALL transplants rapidly produced representative CD19+CD33-lymphoid leukaemias in both mouse strains. In contrast, transplantation of relapsed AML engrafted only in the more myeloid-permissive MISTRG strain as a CD34+/-CD19-CD33+ AML (Figure S2C). Thus, in contrast to the immunophenotypic plasticity seen in MPAL leukaemias with wildtype *MLL* (9), transplantation of a relapsed, fully switched AML was only capable of generating AML.

### Lineage switch relapse can originate in HSPC compartments

To further investigate a potential origin of relapse in an early progenitor or stem cell, we purified haematopoietic stem/progenitor cell (HSPC) populations from diagnostic ALL and relapsed AML and tested purified populations for the presence of *MLL-AF4* targeted sequencing. The *MLL-AF4* translocation was found in the lymphoid-primed multipotent progenitor-like population (LMPP, CD34+CD38-CD45RA+; lymphoid and myeloid potential, but not megakaryocyte-erythroid potential), in the multipotential progenitor population (MPP, CD34+CD38-CD45RA-CD90-; no lineage restriction) and for MPAL1 even in the haematopoietic stem cell-like population (HSC, CD34+CD38-Lin-CD90+) (Figures 2A-C, S3; Table S3). In line with these findings, serial xenotransplantation of LS01P identified a persistent human CD34+CD38-CD45RA-CD90+ HSC compartment across four generations of mice with maintenance of the *MLL-AF4* fusion gene within purified human CD34+ cells (Figures 2D-F). These findings suggest maintenance of a (pre-)leukaemic clone with high malignant self-renewal potential in HSPC populations and support our findings from BCR analysis that, at least in a subgroup of cases, an early multipotent progenitor or HSC can act as the cell of origin for the AML relapse.

**Figure 2.**
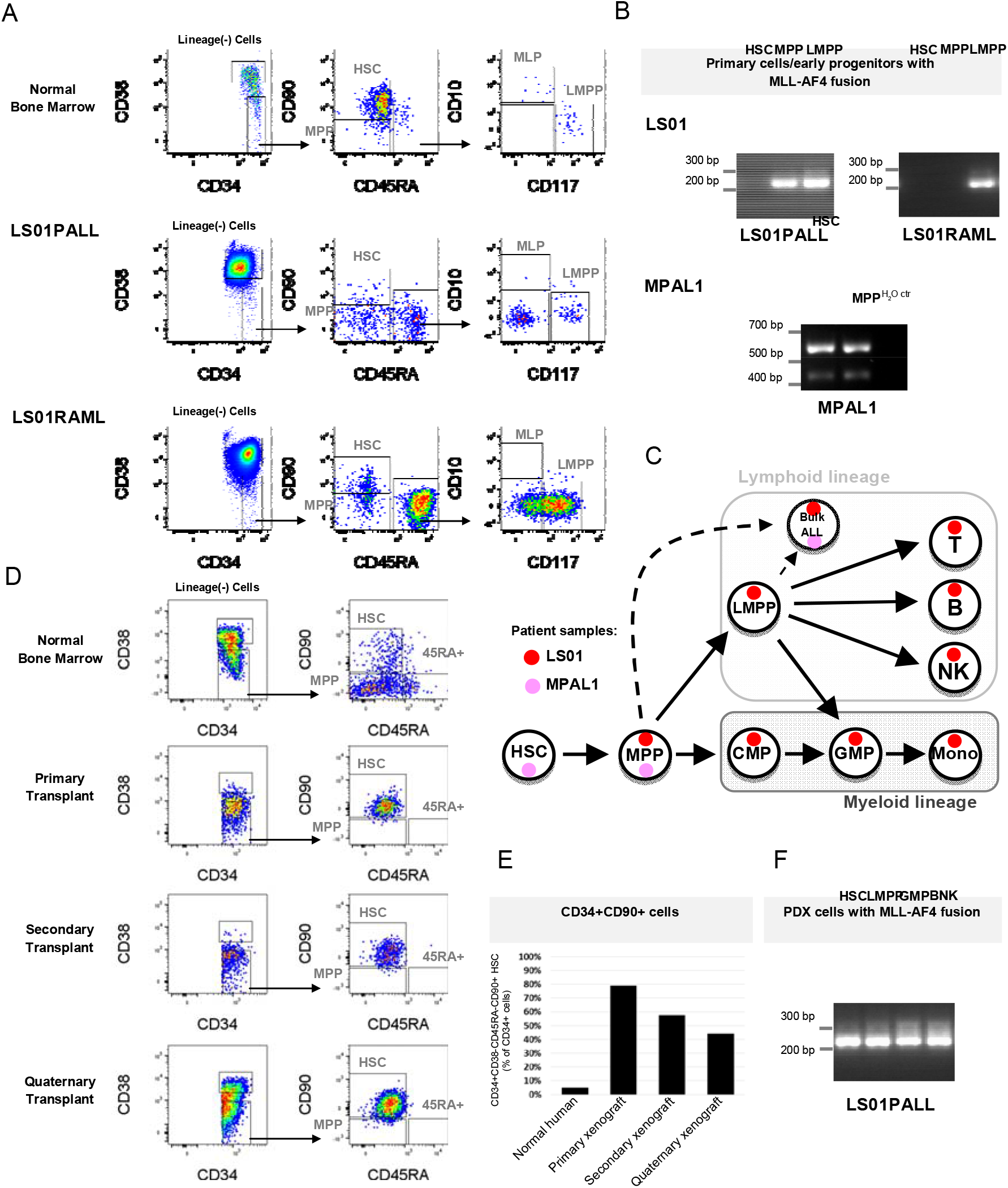
Haematopoietic stem/progenitor populations carry *MLL-AF4*. (A) Flow cytometric sorting strategy for haematopoietic stem/progenitor cell (HSPC) populations. (B) PCR identification of the specific *MLL-AF4* fusion within sorted HSPC populations LS01 and MPAL1 cases. (C) Summary of *MLL-AF4* positivity within different HSPC populations analysed in patients LS01PALL and early progenitors and lymphoid fraction of MPAL1, presented as red and pink circles, respectively. (D) Flow cytometric analysis showing sequential transplantation of LS01 ALL presentation sample across four generations of NSG mouse xenografts. The HSC population (CD34+CD38-CD45RA-CD90+) is maintained across four mice generations. (E) Proportion of bone marrow human CD34+ cells in CD34+CD38-CD45RA-CD90+ HSC gate in all analysed xenografts. (F) PCR identification of the specific *MLL-AF4* fusion within the sorted HSPC xenograft sample.

In concordance with the translocation being present within the early HSPC compartment, we sorted viable differentiated leukocytes and were able to detect the *MLL-AF4* fusion in myeloid and lymphoid lineages including CD34-CD19/3-HLA-DR+CD14/11c+ monocytes, NK, B and mature T cells (Figure 2C, S3A, B; Table S3). These findings imply the existence of a pre-leukaemic progenitor cell, in which *MLL-AF4* does not impose a complete block on haematopoietic differentiation but is compatible with myeloid and lymphoid differentiation. These findings raise the question of which factors and molecular mechanisms affect the ALL and AML lineage choice in *MLL-AF4* leukaemia.

### Lineage switch leukaemia is associated with transcriptional reprogramming

We next investigated the underlying molecular events associated with a lineage switch. To this end we analysed differential gene expression across eight cases, including six LSALs for which RNA was available at both presentation and relapse, and sorted lymphoid and myeloid blast populations of two MPALs. This analysis identified in total 1374 up-(adj. p<0.01, Log Fold change >2) and 1323 down-regulated genes in the AML lineage switches and the myeloid populations of MPAL patients (Table S4). The most substantially down-regulated genes include lymphoid genes such as *PAX5* and *RAG2*, but also the polycomb PRC1 like complex component *AUTS2* and SWI/SNF complex component *BCL7A*, while up-regulated genes comprise several myeloid genes such as cathepsins, cystatins, *PRAM1* and *CSF3R* (Figure 3A). These fingings were consistent, irrespective of which cellular origin of relapse the BCR rearrangement analysis supported (pattern 1, 2 or 3),

**Figure 3.**
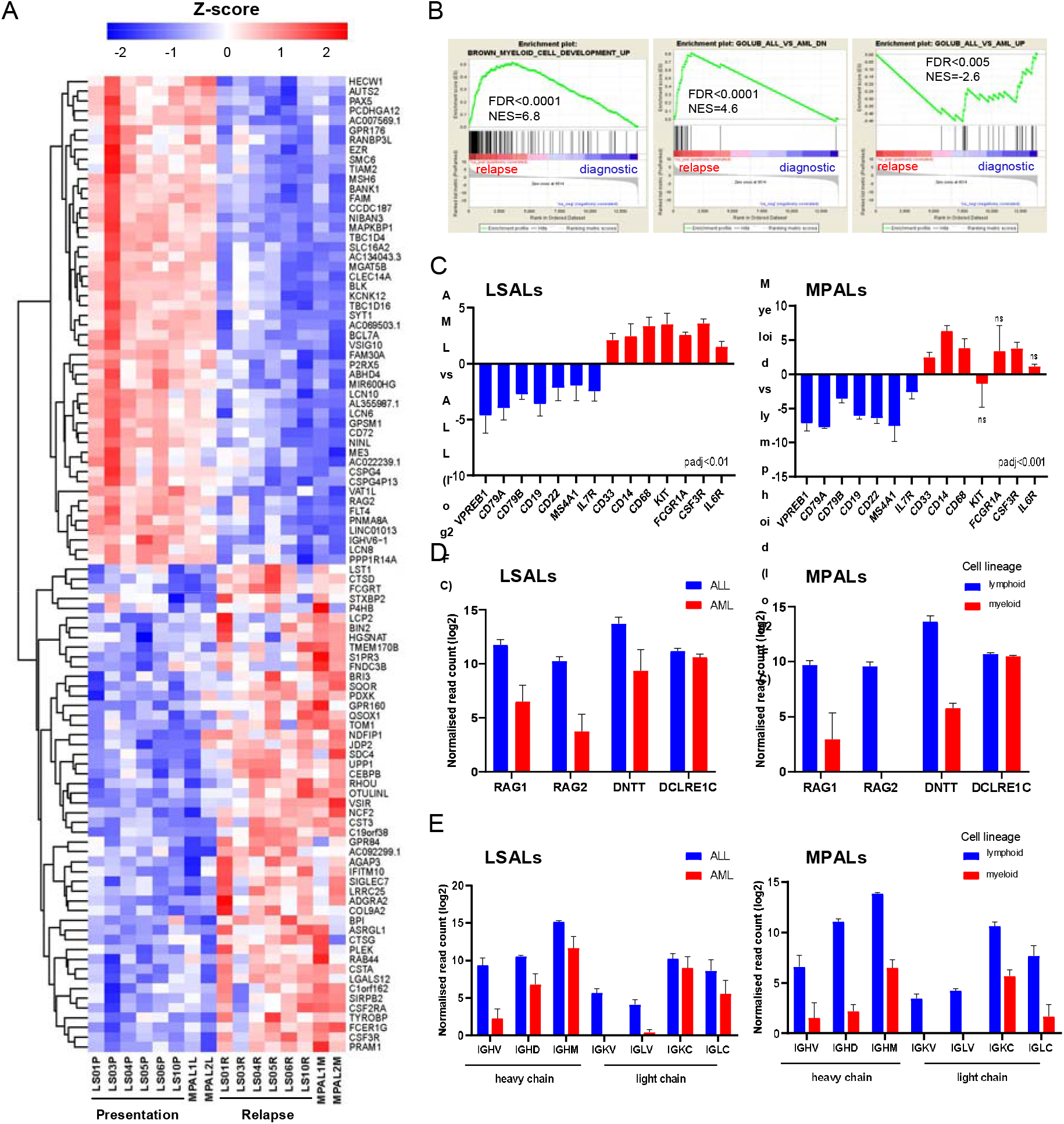
Transcriptional reprogramming in lineage switch and MPAL cases. (A) Heatmap showing the top 100 differentially expressed genes between ALL and AML from six lineage switch (LS01, LS03, LS04, LS05, LS06, LS10) and two MPAL cases, ranked by stat value. (B) Enrichment of myeloid growth and differentiation signature in relapsed samples (left panel) identified by GSEA analyses, also pointing to downregulation of genes highly correlated with acute lymphoblastic leukemia (middle and right panel). Gene set enrichment analyses have been performed based on data derived from six lineage switch samples. FDR – false discovery rate, NES – normalised enrichment score. (C-E) Differential expression of (C) lineage specific, (D) immunoglobulin recombination machinery, and (E) genes encoding immunoglobulin heavy and light chains in lineage switch and MPAL cases. Error bars show standard error of the mean (SEM) for lineage switch cases and ranges for two MPAL cases.

Both gene set enrichment analysis (GSEA) and non-negative matrix factorisation (NMF) showed that presentation and relapse cases with lineage switch have expression signatures similar to previously published *MLL-AF4* ALL and *MLL*r AML cases as well as normal lymphoid and myeloid cell types, respectively (Figures 3B, S4A, B) (10-12). More specifically, lineage switch included increased expression of factors controlling myeloid differentiation (e.g., *CSF3R, KIT)* and changes in haematopoietic surface marker expression (e.g., *CD19, CD22, CD33, CD14, FCGR1A/CD64*), loss of immunoglobulin recombination machinery genes (e.g., *RAG1, RAG2, DNTT*) and reduced expression of genes encoding heavy and light immunoglobulin chains (Figures 3C-E, S4C). Notably, GSEA also indicated impaired DNA repair and cell cycle progression of the AML relapse when compared with diagnostic ALL (Figures S4D). In particular, the reduced self-renewal potential might reflect the transition from ALL with a high incidence of leukaemic stem cells (LSCs) to AML with fewer LSCs.

MLL fusion proteins including MLL-AF4 have previously been shown to directly regulate multiple genes linked to haematopoietic and leukaemic stemness (13-16). For instance, significant changes in gene expression across the *HOXA* cluster, notably with a 50-fold reduction in *HOXA7* expression, represent a major additional disruption of *MLL*r leukaemogenic transcriptional regulation (Figures S5A-C) (13, 14). Furthermore, a very significant portion of target genes including *PROM1, IKZF2* and chromatin modifying factors are highly enriched amongst genes that show lower expression in myeloid lineages (Figures S5D, E, Table S4). In total, 996 out of 5208 bona fide direct target genes of *MLL-AF4* changed expression in the AML relapse (15). These data suggest that the process of lineage switch is associated with a major reorganisation of the *MLL-AF4* transcriptional network and pose the question of which epigenetic regulators are involved in the lineage determination associated with this fusion gene.

### Reorganisation of chromatin accessibility and transcription factor binding upon lineage switch

For case LS01 we had sufficient diagnostic material to perform DNase hypersensitivity site (DHS) analysis and thus link transcriptional changes to altered genome-wide chromatin accessibility. Many differentially expressed genes showed altered chromatin accessibility in proximity to the transcriptional start site (TSS), including genes encoding key hematopoietic surface markers CD33 and CD19, transcription factors and proteins related to differentiation (Figures 4A-C, S6A).

**Figure 4.**
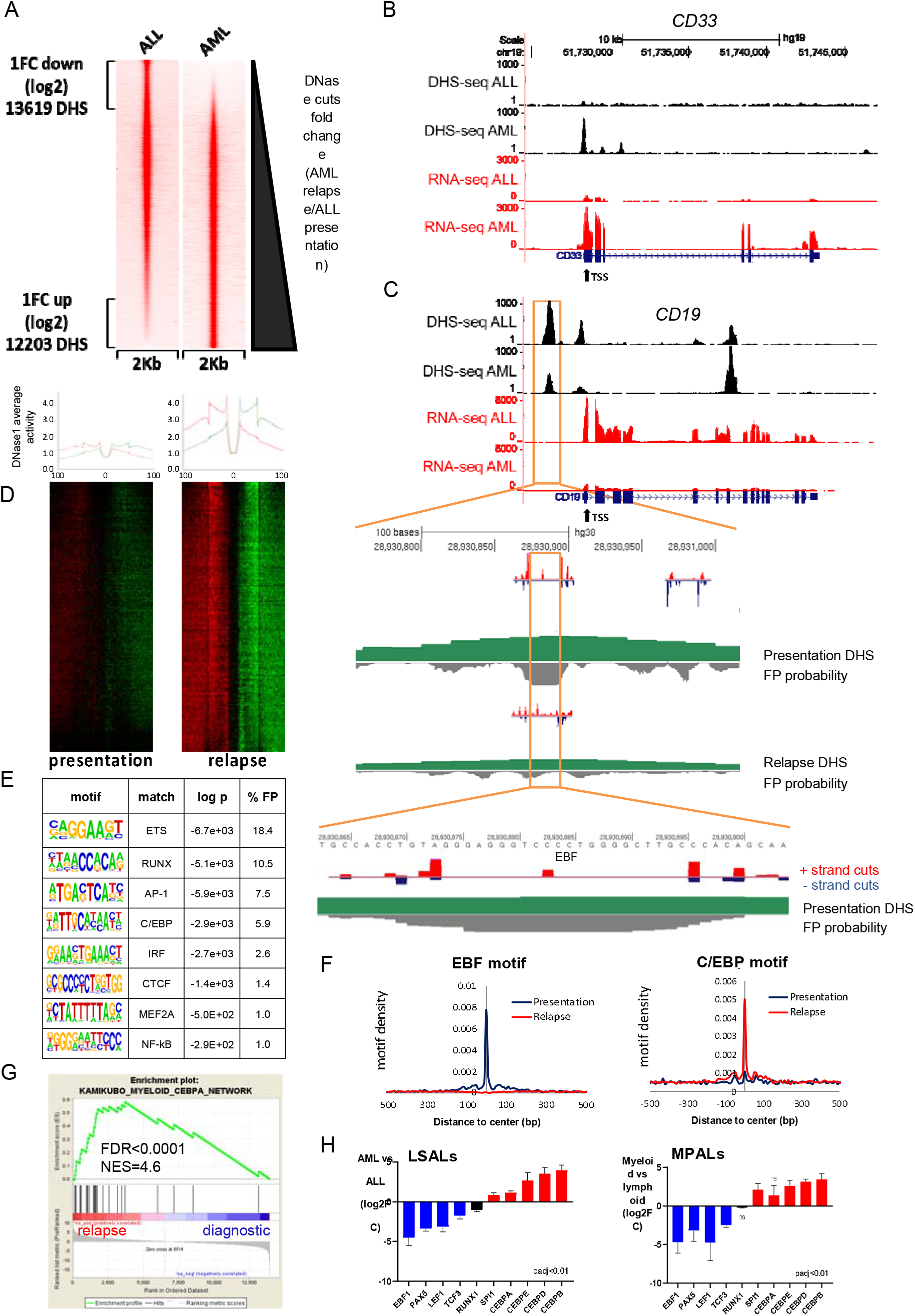
Chromatin re-organisation and differential transcription factor binding underpins lineage switching. (A) DNase hypersensitivity site sequencing identifies 13619 sites with a log2 fold reduction and 12203 sites with a log2 fold increase following lineage switch to AML. (B) University of California, Santa Cruz (UCSC) genome browser screenshot displaying differential expression at lineage specific loci (lower red tracks) accompanied by altered DNase hypersensitivity (upper black tracks) proximal to the transcriptional start site (TSS) of *CD33*. (C) UCSC genome browser screenshot for *CD19* zoomed in on an ALL-associated DHS with EBF occupation as indicated by high resolution DHS-seq and Wellington analysis. FP - footprint. (D) Heat maps showing distal DHS regions specific for AML relapse on a genomic scale. Red and green indicate excess of positive and negative strand cuts, respectively, per nucleotide position. Sites are sorted from top to bottom in order of decreasing Footprint Occupancy Score. (E) De novo motif discovery in distal DHSs unique to AML relapse as compared to ALL relapse as shown in (D). (F) EBF1 and C/EBP binding motifs demonstrate differential motif density in presentation ALL and relapse AML. (G) Enrichment of a myeloid C/EBPA network gene set in signatures associated with relapse AML and diagnostic ALL samples as identified by GSEA (H) Differential expression of TFs cognately binding to differentially accessible motifs shown in (F). TFs whose binding motifs show increased accessibility in ALL are in blue whilst those showing increased accessibility in AML are in red. RUNX1 in black reflects enriched accessibility of different RUNX1 binding sites in ALL and AML. Error bars show SEM or ranges in LSAL and MPAL cases, respectively.

Changes in chromatin accessibility were linked with an altered pattern of transcription factor binding. High resolution DHS-seq (digital footprinting) of presentation ALL and relapse AML, respectively, showed marked genome-wide alterations at sites distal to the TSS, from which several AML and ALL specific *de novo* occupied transcription factor binding motifs were identified (Figures 4C-E, S6B). Lineage switch from ALL to AML was associated with a loss of occupancy of motifs binding lymphoid transcription factors such as EBF or PAX5 and a gain of occupancy of motifs bound by C/EBP, IRF and NF-κB family members (Figure 4E-F). The gain in C/EBP motif binding was associated with the expression of a C/EBPA regulated myeloid transcriptional gene set (Figure 5G). We also observed a redistribution of occupancy of transcription factors controlling both lymphoid and myeloid maturation such as RUNX, AP-1 and ETS members to alternative cognate motifs (Figures 4E and S6B) (17, 18). This finding is exemplified by decreased accessibility of a region located 1 kb upstream of the *CD19* TSS with concomitant loss of EBF binding at this element (Figure 4C). Differential motif enrichments were associated with changes in RNA expression and chromatin accessibility at the genes encoding the corresponding cognate transcription factors *EBF1, PAX5, LEF1* (B lymphoid determinants), *NFKB2* and *CEBPB/D/E* (myeloid determinants), particularly in regions flanking TSSs (Figures 4H, S6B).

**Figure 5.**
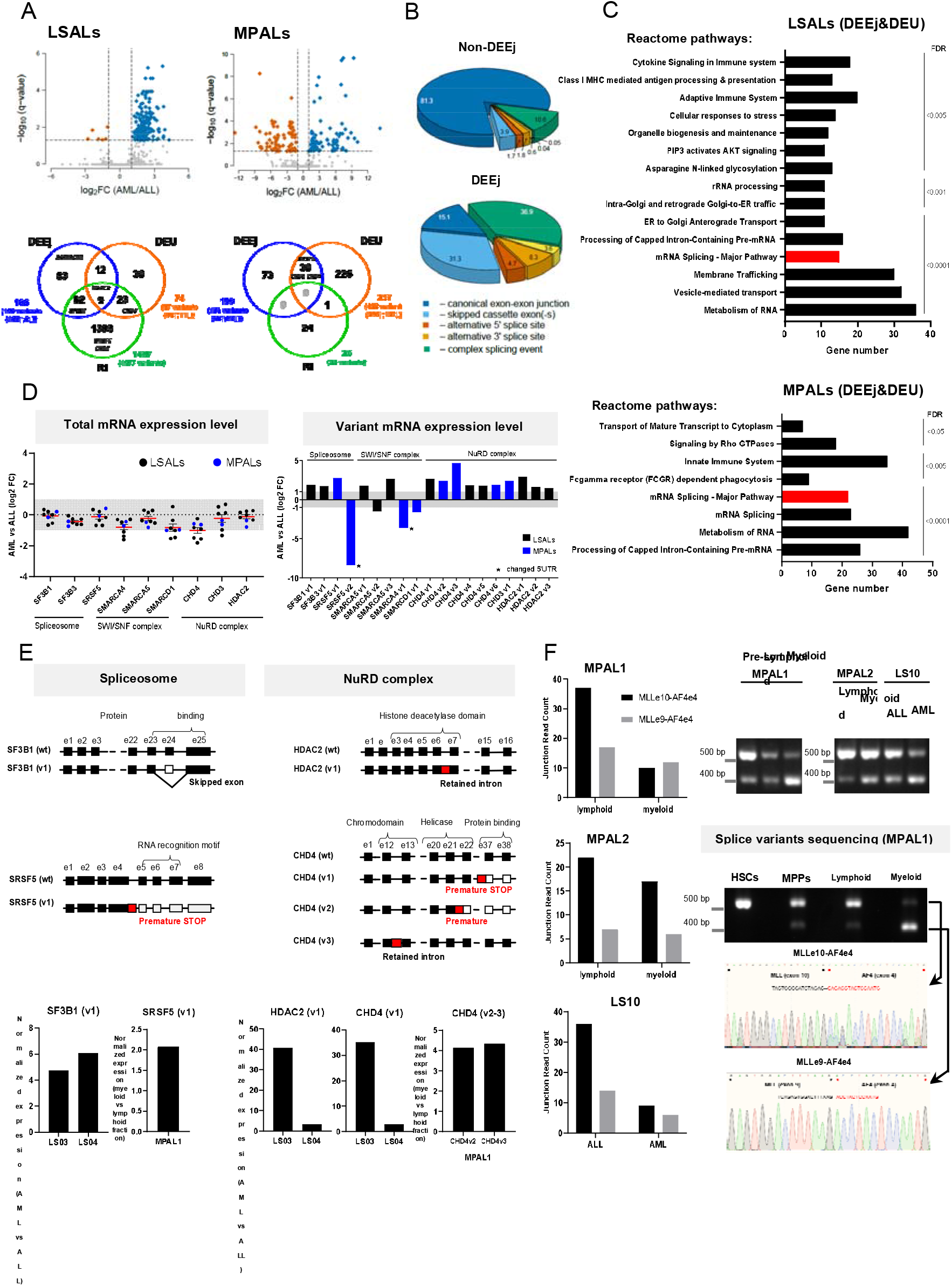
Alternative splicing in lineage switch and MPAL cases. (A) Volcano plots demonstrating differential usage of exon-exon junctions in the transcriptome of AML/myeloid versus ALL/lymphoid cells of lineage switch (LS01, LS03 & LS04) or MPAL patients. The vertical dashed lines represent two-fold differences between the AML and ALL cells and the horizontal dashed line shows the FDR-adjusted q-value threshold of 0.05 (upper panel). Venn diagrams (lower panel) showing distribution of splice variants identified as significantly changed in AML (or myeloid fraction of MPAL patients), including exon-exon junctions (DEEj), differential exome usage (DEU) and retained introns (RI). (B) Pie charts showing the classification of non-differential (non-DEEj) and differential (DEEj) exon-exon junctions. Shown are the percentages of splicing events assigned to a particular mode of splicing. Complex splicing event corresponds to several (two or more) alternative splicing incidents occurring simultaneously in the same sample. (C) Enrichment analysis of affected signalling pathways by the exon-exon junctions (DEEj) and differential exome usage (DEU) in the LSAL AML relapse and myeloid compartment of MPAL patients. Pathway enrichment analysis has been performed with https://biit.cs.ut.ee/gprofiler/gost under the highest significance threshold, with multiple testing correction (g:SCS algorithm). (D) Fold change expression levels of total gene among genes identified to be affected by alternative splicing process (left panel) and differentially spliced variants in lineage switched and myeloid compartments of MPAL patients (right panel). (E) Schematic representation of the affected mRNA structure (and its probable consequence depicted in red) within several selected genes (upper panel). Corresponding normalized expression level (vs reference gene *TBP*) in two tested lineage switch patients (LS03 and LS04) and one MPAL (lower panel). Shown is the ratio of analysed splice variant expression level between AML (or myeloid) and ALL (or lymphoid) populations. (F) Splice variants of *MLL-AF4* identified in MPAL patients and one lineage switch sample (LS10). Left panel represents junction read counts of the fusion oncogene, identified by the RNAseq analysis, with confirmation of the expression of both variants analysed by qRT-PCR (MPAL1, right panel). Both splice variants, further confirmed by Sanger sequencing, showed complete sequences of *MLL*ex9 or *MLL*e10 and either complete or truncated (for 3 nucleotides at the 5’end) *AF*4ex4, respectively.

In conclusion, the transition from lymphoid to myeloid immunophenotype is associated with global lineage specific transcriptional reprogramming and genome-wide alteration in chromatin accessibility and transcription factor binding.

### Lineage switch changes alternative splicing patterns

Lineage fidelity and determination are not only linked to differential gene expression but may also include co- or post-transcriptional mechanisms. It has been previously demonstrated that lineage commitment during haematopoiesis leads to substantial changes in alternative mRNA splicing patterns (19). Furthermore, we recently showed that the *AML*-*ETO* (RUNX1-RUNX1T1) fusion protein controls leukaemic self-renewal by both differential gene transcription and alternative splicing (20). To complement the transcriptional analysis we therefore sought to define the different composition of RNA isoforms in lymphoid and myeloid populations from lineage switch and MPAL cases. Here we focussed on three lineage switch patients and the two MPAL patients whose RNA-seq data provided sufficient read depth for the analysis of exon-exon junctions, exon usage and intron retention (Figures S7A-C). We detected in total 2630 retained introns (RIs) shared amongst the three lineage switches with 653 and 343 RIs exclusively found in the diagnostic ALL or the AML relapses, respectively. This was complemented by 97 exons (DEUs) and 193 exon-exon junctions (DEEjs) differentially used between diagnostic ALL or the AML relapses. In contrast, this analysis identified only 43 RIs present in both MPAL cases with 18 and 6 introns specifically retained in either the lymphoid or myeloid subpopulation, respectively (Figure S7D, Table S5). Intersection of the affected genes identified 21 shared genes out of 166 DEEjs and 74 DEUs. MPALs had 420 DEUs and 155 DEEjs affecting 257 and 103 genes with 30 genes having both DEUs and DEEjs (Figure 5A, Table S6). While more than 80% of the non-differential exon-exon linkages were canonical, this was true for only 15% of the DEEjs. Here, non-canonical exon skipping and complex splicing events contributed more than 30% each, most prominently to differential alternative splicing (Figure 5B).

Pathway analysis revealed an enrichment of alternatively spliced genes in immune pathways including antigen processing, membrane trafficking and FCGR-dependent phagocytosis reflecting the change from a lymphoid to a myeloid state (Figures 5C, S7E). Furthermore, it highlighted RNA processing and maturation including mRNA splicing, processing of capped intron-containing pre-mRNAs and rRNA processing. Indeed, myeloid populations expressed 4-6-fold higher alternatively spliced *SF3B1* and *SRSF5* levels than their matched lymphoid populations (Figure 5D, E). In addition, we noted a significant number of genes encoding epigenetic modulators including *KDM5C, HDAC2* and several *CHD* members being differentially spliced in AML relapse or myeloid subpopulations of MPALs (Figure 5D, E).

Analysing the occurrence of *MLL-AF4* isoforms showed that the fusion site of *MLL-AF4* itself was subject to alternative splicing in lineage switch and MPAL. Three cases shared the breakpoints in introns 10 and 4 of *MLL* and *AF4*, respectively. By examining RNA-seq and competitive RT-PCR data, we identified two co-occurring isoforms with either *MLL* exon9 or exon10 joined to *AF4* exon4 (Figure 5F). Interestingly, lymphoid populations expressed a higher ratio of exon10/exon4 over exon9/exon4 than the myeloid populations in all three cases examined. Furthermore, the HSC and MPP-like populations of MPAL1 showed mainly expression of the exon10/exon4 splice variant of *MLL-AF4*, thus resembling more the lymphoid than the myeloid phenotype (Figure 5F). In conclusion, altered isoform expression of *MLL-AF4* may contribute to lineage choice and the phenotypic switch.

### The mutational landscape of lineage switch

Next, we examined the mutational landscape of lineage switched *MLL-AF4* leukaemias by performing exome sequencing on the entire cohort. In keeping with published data on newly presenting *MLL*r acute leukaemias (12, 21), exome sequencing of presentation ALL samples confirmed a relatively quiet mutational landscape in infant ALL cases, with median of 13 nonsynonymous somatic single nucleotide variants (SNVs) or insertions/deletions (indels) predicted to be deleterious to protein function (Table S7). Many of these were present in less than 30% of reads and considered sub-clonal. The most commonly mutated genes at presentation were *NRAS* (3 cases) and *KRAS* (5 cases) (Figure 6A) as described previously (12). In contrast, relapse AML samples contained a median of 46 deleterious somatic SNVs and indels (Table S7). This increase can be mainly attributed to three samples (LS03RAML, LS07RAML and LS08RAML) that carried deleterious mutations in DNA polymerase genes in the respective major clones linked to hypermutator phenotypes (22, 23) (Figure 6A). However, we observed this phenotype only in three out of ten relapses, arguing against this phenomenon being a general requirement for the lineage switch in relapse. Similarly, many of the predominantly subclonal mutations identified in presentation ALL samples, including half of *RAS* mutations, were subsequently lost at relapse, indicating alternative subclones as the origin of relapse (Figure 6A). Finally, both MPALs harboured several mutations that were exclusively found in either the lymphoid or myeloid subpopulation indicating the presence of subclones with a lymphoid and myeloid bias (Figure 6A).

**Figure 6.**
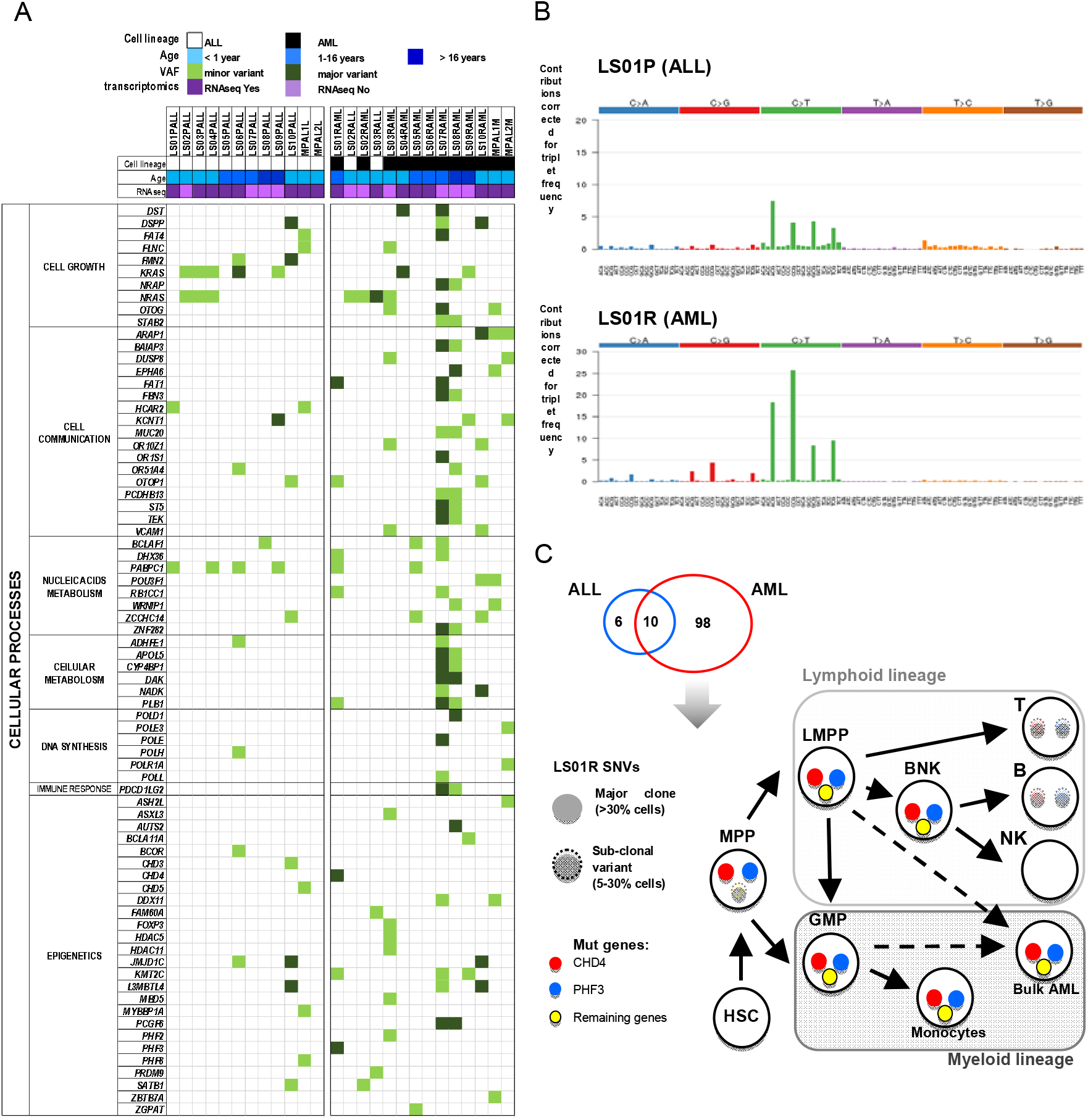
Molecular characterisation of lineage switch *MLL-AF4* leukaemias. (A) Whole exome sequencing (WES) data showing genes recurrently mutated within the analysed cohort, involved in the cell growth, communication and metabolism and genes mutated in single cases belonging to the same function protein complexes (e.g. DNA polymerases, epigenetic complexes). Data are presented according to the disease timepoint/cell lineage and age of the patient. Depicted are major single nucleotide variants (SNVs) that were found in >33% of reads and minor SNVs in <33% reads. (B) Clonal SNV patterns identified by whole genome sequencing (WGS) in LS01 ALL and AML samples, constructed from counts of each mutation-type at each mutation context, corrected for the frequency of each trinucleotide in the reference genome. (C) Comparison of the whole exome sequencing and RNAseq data obtained for LS01 patient identifies 6, 98, and 10 SNVs expressed only in ALL, AML, and shared between ALL and AML, respectively. 12 SNVs exclusive for the AML relapse, predicted (by Condel scoring) to have deleterious effects, were subjected to multiplex PCR followed by next-generation sequencing analysed within each of the purified haematopoietic sub-populations. Circles with solid colour indicate VAF >30%, light colour and dashed line indicates VAF 5-30%. Remaining genes (yellow circle) represent the 10 other SNVs (out of 12 SNVs) which showed the same pattern in the frequency of mutation acquisition.

Next, we examined the mutation patterns of patient LS01 in greater detail. ALL and AML contained 104 and 3196 SNVs with a variant allele frequency (VAF) ≥0.3 respectively, with only 22 shared SNVs between both samples. Globally the most prevalent type of SNV was the C to T transition in the DNA of both ALL and AML samples (Figure 6B). However, the contribution of underlying single base substitution (SBS) signatures differed between diagnosis and relapse. Three different signatures (SBS16, 5 and 1) explained about 60% of the SNPs found in ALL, while SBS1 seemed to explain more than 50% of all SNPs in the AML (Figure 6B) suggesting a mutational clock as the main driver of the evolution of relapse (24). Further inspection of the pattern revealed a mutational signature mainly comprising C to T transitions and to a lesser degree C to G transversions in NCG triplets, raising the possibility that thiopurine maintenance treatment may have increased the mutational burden, resulting in lineage switched relapse in this patient (25).

Twelve deleterious SNVs were identified as unique to the relapse sample of case LS01. The availability of viable cellular material allowed us to investigate the order of acquisition of these secondary mutations within the structure of the normal haematopoietic hierarchy. We sorted this sample to isolate HSC-, MPP-, LMPP- and GMP-like and later populations. Using a targeted deep sequencing approach we then examined these populations for the presence of those 12 SNVs. This analysis showed an increasing number of mutations during the differentiation from MPPs through LMPPs to GMPs. Amongst them, only *PHF3* and *CHD4* mutations were present within the purified CD34+CD38-CD45RA-CD90-MPP-like fraction with VAF≥0.3 (Figure 6C and Table S3). In contrast, LMPP- and GMP-like populations contained all 12 SNVs at high VAF (Table S3). These findings identify the mutation of *CHD4* and *PHF3* as the earliest genetic events during relapse evolution and suggest them as potential drivers of an *MLL-AF4* positive, non-lineage committed, pre-leukaemic precursor population. Subsequent accumulation of additional mutations likely establishes the fully developed leukaemia with more mature haematopoietic/myeloid immunophenotypes.

### Perturbation of CHD4 and PHF3 disrupts lymphoid development in MLL-AF4 expressing cells

The two earliest mutations within LS01 relapse were identified in the Nucleosome Remodelling and Deacetylation complex (NuRD) member *CHD4* and the plant homeodomain finger containing *PHF3*. PHF3 is a member of a family of transcriptional regulators that have been suggested to link the deposition of histone marks to the regulation of transcription (26). PHF3 itself has been recently identified as an inhibitor of transcription elongation by competing with TFIIS for binding to the C-terminal domain of RNA polymerase II (27). NuRD is a multiprotein transcriptional co-repressor complex with both histone deacetylase and ATP-dependent chromatin remodelling activity. It is a critical factor in the lymphoid lineage determination in part directed by the transcription factor IKZF1 (28-30). Both the CHD4 R1068H and PHF3 K1119I mutations affect highly conserved residues (Figures 7A, S8A) that are predicted by the Condel classifier (31) to disrupt protein function. Specifically, the CHD4 R1068H mutation has previously been linked to defects in cardiac development (32).

**Figure 7.**
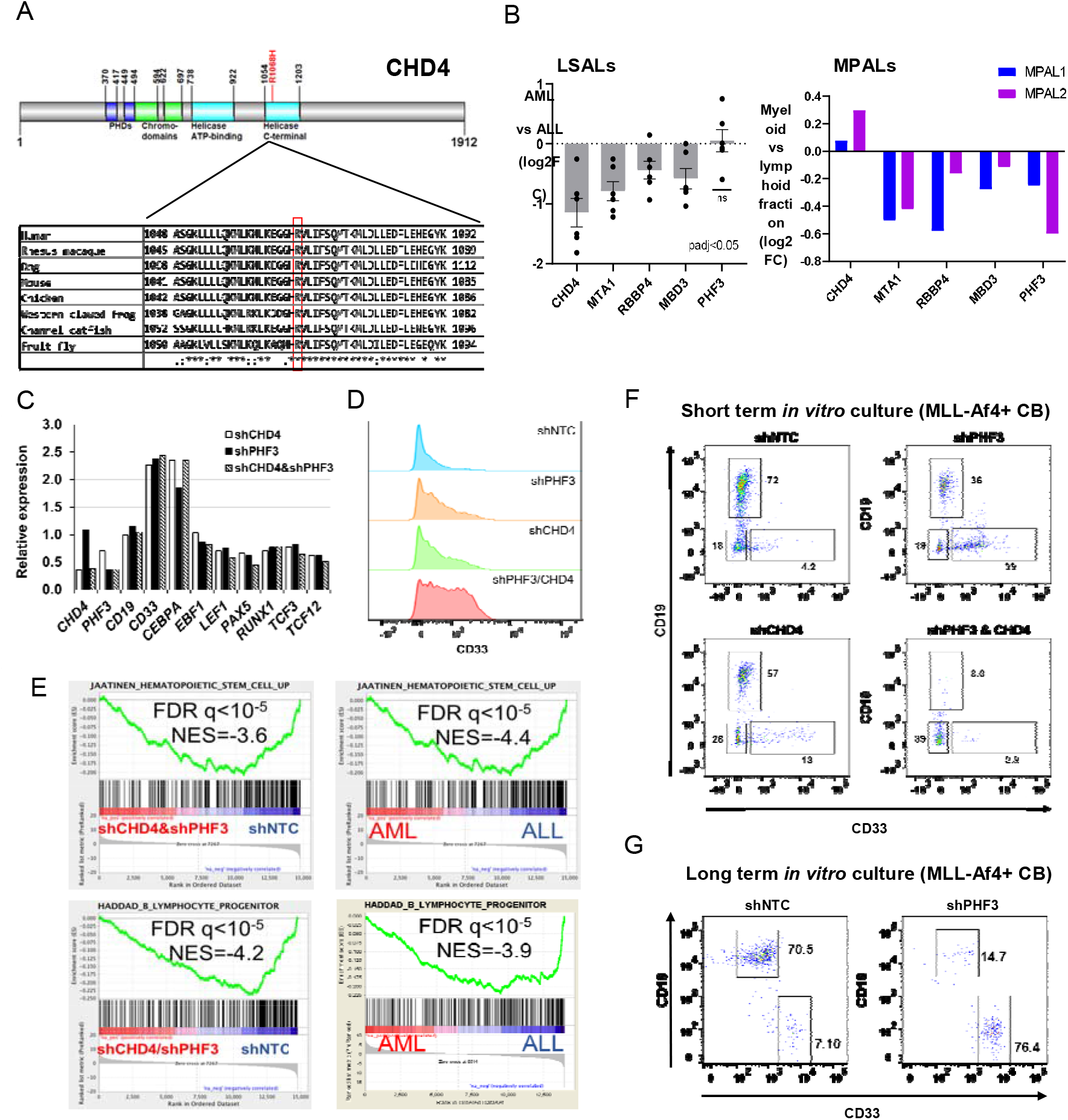
Epigenetic modulatory genes influence lineage specific expression profiles. (A) CHD4 scheme; the R1068H mutation is located in the critical helicase domain of CHD4 at a highly conserved residue. An * (asterisk) indicates positions which have a single, fully conserved residue, a : (colon) indicates conservation between groups of strongly similar properties - scoring > 0.5 in the Gonnet PAM 250 matrix, a. (period) indicates conservation between groups of weakly similar properties - scoring =< 0.5 in the Gonnet PAM 250 matrix. (B) Fold change in expression of NuRD complex members (*CHD4, MTA1, RBBP4, MBD3*) and *PHF3* following lineage switched relapse (left panel) and in MPAL cases (right panel). (C) Expression of lineage specific genes following knockdown of *PHF3, CHD4* or the combination, relative to non-targeting control construct in the *MLL-AF4* positive ALL cell line, SEM. (D) Flow cytometric analysis of the surface CD33 expression following knockdown of *PHF3, CHD4* or the combination in the SEM cell line. shNTC – non-targeting control. (E) Gene set enrichment analysis of RNA sequencing data derived from: knockdown of *PHF3* and *CHD4* in the SEM cell line (left panel) and lineage switch leukaemia cases (right panel). Shown is negative correlation of shCHD4/shPHF3 and relapse samples with Jaatinen haematopoietic stem cell signature (upper panel) and Haddad B-lymphocyte progenitor signature (lower panel). (F) Expression of lineage specific cell surface markers CD19 (lymphoid) and CD33 (myeloid) following culture of *MLL*-*Af4* transformed hCD34+ cord blood progenitor cells in lymphoid permissive conditions. Knockdown of *PHF3, CHD4* or the combination disrupts the dominant lymphoid differentiation pattern seen in non-targeting control (shNTC). (G) Assessment of *PHF3* knockdown influence on the surface marker expression after longer incubation period (33 days); *CHD4* knockdown impaired cellular survival upon longer *in vitro* culture.

Whilst mRNA expression levels of the mutant CHD4 R1068H were unaffected in case LS01, across the remaining cases analysed, expression of *CHD4* and additional NuRD complex members *MBD3, MTA1* and *RBBP4* was reduced following lineage switched myeloid relapse (Figures 7B). Furthermore, *CHD4* was affected in the MPAL myeloid populations by non-canonical alternative splicing leading to premature termination of translation, indicating that this particular pathway was severely disrupted across the cohort, irrespective of the putative cell of origin of relapse.

The relevance of *CHD4* and *PHF3* in regulating lymphoid versus myeloid lineage choice was further supported by ARACNE analysis using published ALL (33) and AML (34) expression datasets (n=216) which we used to reverse-engineer a mutual information network. This network was trimmed to represent only genes significantly associated with the difference between AML and ALL thus reflecting the likely influence of genes of interest upon genes associated with the difference between AML and ALL. This analysis found *CHD4* and *PHF3* to be the mutated genes with the highest number of edges within the network (*PHF3* – 21 edges, p=0.010; *CHD4* – 12 edges, p=0.0005, Figure S8B and Table S8), implicating them as causal to the lymphoid/myeloid distinction identified within primary ALL/AML.

To establish a direct functional link from these mutations to the loss of lymphoid lineage commitment in *MLL-AF4* ALL, we knocked down *CHD4* and *PHF3* in the *MLL-AF4* positive cell line, SEM. Depletion of *CHD4* and *PHF3* alone or in combination resulted in increased expression of the myeloid transcription factor *CEBPA* and reduced expression of lymphoid transcription factors including *LEF1, PAX5, TCF3* and *TCF12* (Figure 7C). These changes were accompanied by a more than twofold increase in CD33 expression in the two *MLL-AF4*-expressing ALL cell lines SEM and RS4;11, while CD33 levels in two *MLL-AF4*-negative ALL cell lines remained unaffected, supporting the importance of these epigenetic regulators specifically within the *MLL*-*AF4* context (Figures 7D and S8C). GSEA following *PHF3* and *CHD4* knockdown indicated loss of HSC and B-lymphocyte progenitor gene expression signatures (Figure 7E, left panel), similar to what was observed with the transcriptomes of the lineage switch leukaemia cases (Figure 7E, right panel). In line with these cell line experiments, knockdown of *CHD4* and *PHF3* in an ALL PDX generated from the first relapse of patient LS03 also resulted in a more than twofold increase in CD19/CD33 double and CD33 single positive cells, demonstrating that perturbation of these two genes is also able to change the immunophenotype of primary *MLL-AF4* ALL cells in a case susceptible to trans-differentiated relapse (Figure S8D).

In order to examine the role of additional mutations of chromatin modifiers found in our cohort and known to regulate lineage choice, we investigated the impact of the PRC1 members *PCGF6* and *AUTS2* on CD33 expression in SEM cells. *PCGF6* is mutated in LS07RAML and LS08RAML and has known roles in B lymphoid malignancy (35). *AUTS2* is both mutated in LS08RAML and highly expressed in all lymphoid populations examined (Figures 3A, 6A). STRING network analysis demonstrated close functional associations within PRC1 and NuRD complexes and their shared associations (Figure S8E). Mutual interactions include CBX2, a PRC1 complex member, which shows a similar expression pattern between lineage switch and MPAL cases (Figure S8F). While knockdown of *AUTS2* did not change CD33 levels, depletion of *PCGF6* also increased CD33 surface expression in SEM cells, further supporting the notion of epigenetic factors regulating lineage determination in ALL (Figures S8G). Furthermore, GSEA also indicated impaired function of PRC1 and PRC2 complexes in the AML relapse compared with the presentation ALL with down-regulation of their respective target genes (Figure S8H).

Given that the relapse-initiating cell can arise within an uncommitted, *MLL-AF4* translocated HSPC population, we went on to assess the impact of *CHD4* and *PHF3* function loss in a human cord blood model, which harbours a chimeric *MLL-Af4* fusion and can be differentiated both into myeloid and lymphoid lineages (36). Knockdown of either *CHD4* or *PHF3* under lymphoid culture conditions significantly impaired lymphoid differentiation potential, whilst co-knockdown of *CHD4* and *PHF3* disrupted differentiation entirely (Figures 7F, G and Table S9). Transcriptomic analysis of the sorted populations revealed that CD33 positive cells exhibited metagene expression pattern similar to *MLL*r AML, while the pattern describing CD19+ cells was most similar to *MLLr* ALL, thus confirming that changes in surface marker expression were associated with the corresponding changes in the transcriptomic profiles (Figure S8I).

Taken together, our data show the important role of the NuRD (CHD4), PHF3 and other (PRC1) repressive complexes in the epigenetic control of lymphoid lineage choice. In particular, dysregulation of *CHD4*/NuRD was mediated by mutation, down-regulation of expression and differential splicing across the cohort, irrespective of the cellular/clonal origin of relapse. These data support a role for these factors in the strong lineage determining capacity of *MLL-AF4* whilst their loss undermines both the execution and the maintenance of the lymphoid lineage fate.

## Discussion

This study describes impaired epigenetic control as being central to the phenomenon of lymphoid-myeloid lineage switch in *MLL-AF4*-positive leukaemia and identifies the cell of origin of relapse into AML. We found that the origin of relapse was heterogeneous. Relapse can directly evolve from pro-B-like ALL blast populations, which agrees with the general self-renewal capacity of ALL cells (37), but can also originate within the HSPC compartment. Indeed, analysis of both patient and xenotransplanted cell populations from diagnostic ALL identified *MLL-AF4* fusion transcripts in MPP- and HSC-like cells. This finding agrees with recently published data pointing at MPP cells as the origin of *MLL-AF4* leukaemia (38) and is in line with transcriptomic similarities between t(4;11) ALL and Lin-CD34+CD38-CD19-fetal liver cells, again suggesting an HSPC as the cell of origin (39).

Irrespective of the cellular origin of the relapse, lineage switching was associated with a major rewiring of gene regulatory networks. At the level of transcriptional control, the decision for lymphoid development relies not only on the activation of a lymphoid transcriptional program, but also on the silencing of a default myeloid program (40). That decision is enacted by lymphoid master regulators including EBF1, PAX5 and IKAROS, which represent genes commonly mutated in precursor B-ALL (41-43). Pax5^-/-^ pro-B cells which lack lymphoid potential, whilst capable of differentiating down erythro-myeloid lineages *in vitro*, still maintain expression of early B cell transcription factors *EBF1* and *E2A* (*TCF3*) (40). In contrast, we show that lineage switching *MLL-AF4* pro-B leukaemic relapse is associated with significant reduction in expression and binding of these earliest B lymphoid transcription factors. Their loss is linked with changes in the *MLL-AF4* transcriptional programme, notably within *HOXA* cluster genes (13, 14) which likely results in a wider reorganisation of malignant haematopoietic transcriptional networks, ultimately leading to a myeloid differentiation fate.

Similar to the *Pax5* knockout (40), loss of IKAROS DNA-binding activity prevents lymphoid differentiation (29). NuRD co-operates directly with IKAROS to repress HSC self-renewal and subsequent myeloid differentiation, permitting early lymphoid development (29, 44, 45). We found that the abrogation of this pathway through multiple mechanisms was central to the lineage switch from ALL to AML. Lineage switch was either associated with mutation, reduced expression or, in the case of two MPALs, alternative splicing of *CHD4* and other NuRD components. Long term knockdown of *CHD4* was not tolerated in our cord blood culture. This is in line with reports showing that complete loss of *CHD4* impairs leukaemic proliferation (46, 47), both myeloid and lymphoid differentiation of HSPCs and causes exhaustion of HSC pools (44), indicating that basal CHD4 expression is required for maintaining AML. Moreover, our observation that a 60% *CHD4* knockdown is associated with the activation of pluripotency gene signatures is in line with the finding that a partial inhibition of *CHD4* supported induction of pluripotency in iPSCs, while a complete deletion eliminated cell proliferation (48).

Whilst our study has investigated the rare clinical occurrence of lineage switching, recent studies have identified core NuRD and PRC1 complex members as being direct targets of *MLL*-*AF4* binding (16, 49). We therefore hypothesise that epigenetic regulator genes are co-opted during *MLL*-*AF4* leukemogenesis and mediate fundamental lineage specific decision-making processes, in this case the suppression of the myeloid lineage program. Multiple routes to their dysregulation may result in escape from this lineage restriction. Our finding that frontline chemotherapy itself may contribute to relapse highlights the urgent need to find alternative therapies for this high-risk leukaemia. Equally, however, the associated loss of B cell surface markers (*e*.*g*., CD19) provides an alternative mechanism for relapse following CAR-T cell or blinatumomab therapy (50, 51) in addition to mutations, alternative splicing (52, 53) and T cells trogocytosis (54). Whilst these therapies target lineage specific surface markers, lineage-switched (pre-)leukaemic progenitor populations escape epitope recognition and provide a potential clonal source for the relapse. Given the increasing use of advanced immunological therapies, a detailed understanding of the molecular processes underlying lineage determination and switching will be critical for developing new strategies to avoid this route to clinical relapse.

## Methods

### Patient samples and data

Patients were diagnosed by local haematology specialists according to contemporary clinical diagnostic criteria based on morphology and immunophenotypic analysis. All patient samples were collected at the point of diagnosis, remission following treatment or relapse and stored with written informed consent for research in one of six centres (Newcastle Haematology Biobank, Newcastle, UK; University Hospital Schleswig-Holstein, Kiel, Germany; Dmitry Rogachev National Medical Research Center of Pediatric Hematology, Oncology and Immunology, Moscow, Russia; Haematological Malignancy Diagnostic Service, Leeds, UK; Princess Maxima Center for Pediatric Oncology, Utrecht, The Netherlands; Cincinnati Children’s Hospital Medical Center, Cincinnati, USA). Mononuclear cells were isolated from bone marrow or peripheral blood by density centrifugation followed by immediate extraction of DNA or RNA, or cryopreservation in the presence of 10% v/v DMSO.

Samples were requested and used in accordance with the ethical approvals granted to each of the local/institutional ethical review boards (NRES Committee North East - Newcastle & North Tyneside 1, UK, reference 07/H0906/109+5; Medical Faculty Christian-Albrechts University, Kiel, reference A 103/08; Dmitry Rogachev National Medical Research Center, Moscow, references MB2008: 22.01.2008, MB2015: 22.01.2015, ALL-REZ-2014: 28.01.2014; Haematological Malignancy Research Network, Yorkshire, UK, reference 04/Q1205/69; Haematological Malignancy Diagnostic Service, Leeds, UK, reference 14/WS/0098; Erasmus MC METC, Netherlands, reference MEC-2016-739; IRB of Cincinnati Children’s Hospital, USA, reference 2010-0658) and in accordance with the Declaration of Helsinki. Each patient/sample was allocated an anonymised reference and no identifiable information was shared.

### DHS library generation, sequencing, and mapping

DHS analysis was performed as described previously (55). DNase I (Worthington, Cat# LS006328) digestion was performed using ∼5 million patient sample cells using 8 units (presentation) or 14 units (relapse) for 3 min at 22°C in a 1 mM CaCl_2_ supplemented buffer. Nuclear proteins were digested with 1 mg/ml Proteinase K overnight at 37°C. DNase I digestion products were size-selected on an agarose gel, cutting below 150 bp. High-throughput sequencing libraries were prepared from 10 ng of size-selected material, using the Kapa Hyperprep kit as per manufacturer’s instruction. Libraries were sequenced with 50 bp single-end reads on an Illumina HiSeq 2500 sequencer according to manufacturer’s instructions.

Fastq files were generated using bcl2fastq (1.8.4) and subsequently aligned to the hg19 assembly (NCBI Build 37) with the use of bowtie2 (2.1.0), with –very-sensitive-local as a parameter. Read coverage generation and peak detection were carried out using MACS 1.4.1 using --keep-dup=all –g hs -w -S. Pairwise comparisons were performed as previously described (55). Digital footprinting was carried out using the Wellington package using default parameters (56). Differential footprinting analysis was carried out on footprints using the Wellington-bootstrap (57) package with default parameters. Average profiles and heatmaps were obtained using the functions dnase_average_profile and dnase_to_javatreeview from the Wellington package. Heatmaps were plotted using Java TreeView.

### Exome sequencing

Germline DNA from cases LS08 and LS09 were extracted from formalin fixed paraffin embedded remission bone marrow using QIAamp DNA FFPE Tissue Kit (Qiagen, Cat#56404). Other DNA samples were extracted from either bone marrow or peripheral blood using AllPrep DNA/RNA Mini Kit (Qiagen, Cat#80204), QIAamp DNA Mini Kit (Qiagen, Cat#51306), or innuPREP DNA/RNA Mini Kit (Analytik Jena, Cat#845-KS-2080050), according to manufacturers’ instructions. The exons were captured using SureSelect XT2 Human All Exon V6 (Agilent), and sequenced by paired-end 75 bp sequencing on HiSeq4000 (Illumina), resulting in roughly 45 million reads per sample. DNA from the myeloid and lymphoid cellular compartments derived from MPAL patients samples, were pre-processed with KAPA HyperPlus Kit (Roche) followed by exons enrichment with KAPA HyperCapture Kit (Roche), and sequenced by paired-end 300 bp sequencing on NovaSeq6000 (Illumina), resulting in roughly 25 million reads per sample.

Raw reads were aligned to human reference genome (hg19 or hg38 for lineage switch or MPAL patients, respectively) using Burrows-Wheeler Aligner (BWA) 0.7.12 and were processed using the Genome Analysis Toolkit (GATK, v3.8 or 4.1). MuTect (v1.1.7) and MuTect2 (4.1) were used to identify somatic variants for each matched sample pair. Variants were annotated using Ensembl Variant Effect Predictor (VEP, version 90).

### RNA sequencing

Total RNA was extracted with AllPrep DNA/RNA Mini Kit (Qiagen, Cat#80204), innuPREP DNA/RNA Mini Kit (Analytik Jena, Cat#845-KS-2080050), or TRIzol (Thermo Fisher Scientific, Cat# 15596026) followed by RNeasy Mini Kit (Qiagen, Cat#74106) from either bone marrow or peripheral blood, according to manufacturers’ instructions. Messenger RNA was captured using NEBNext Ultra Directional RNA Kit in combination with NEBNext poly(A) mRNA Magnetic Isolation Module or KAPA RNA HyperPrep Kit with RiboErase (HMR) in case of lineage switch or MPAL patients respectively, and submitted for paired-end 150 bp sequencing on HiSeq4000 (Illumina) or paired-end 300 bp sequencing on NovaSeq6000 (Illumina) depending on the analysed patients group. For each sample, transcript abundance was quantified from raw reads with Salmon (version 0.8.2) using the reference human transcriptome (hg38) defined by GENCODE release 27. An R package Tximport (version 1.4.0) was used to estimate gene-level abundance from Salmon’s transcript-level counts. Gene-level differential expression analysis was performed using DESeq2 (version 1.16.1). Differential splicing events were identified in both presentation/relapse pairs or lymphoid/myeloid cellular fractions, using pipeline described previously (20).

### Whole genome sequencing

Presentation, remission and relapse DNA samples from case LS01 were sequenced by Illumina UK and analysed using the remission sample as the matching normal. Sequencing reads were aligned to the human GRCh37.1 reference genome using ISAAC (58). Identification of somatic SNVs and small somatic indels (<50 bp) was performed by Strelka (59). Large structural variants (including deletions, inversions, duplications and insertions all >50bp and translocations) were called by Manta (60).

### Nested multiplex PCR and targeted sequencing

The 12 mutation candidate driver genes and *MLL-AF4* LS01RAML were amplified using gDNA as the template by nested multiplex PCR method. The primers were designed using Primer Express (Applied Biosystems) software.

PCR amplifications were carried out using Phusion® High-Fidelity PCR Master Mix with HF Buffer (NEB, Cat#M0531L) according to the manufacturer’s instructions (25 µl reaction), plus 80 – 200 nM of each primer set. The first multiplex reaction parameters were: one cycle at 98°C for 2 min, thirty cycles of 98°C for 10 s, 63°C for 30 s, and 72°C for 30 s, followed by one cycle at 72°C for 10 min. The products were diluted 500-fold, and 1 µl used as the template for the second PCR reactions (25 µl reaction). The second/nested multiplex reaction parameters were: one cycle at 98°C for 2 min, twenty cycles of 98°C for 10 s, 65°C for 30 s, and 72°C for 30 s, followed by one cycle at 72°C for 10 min. The amplicons that were taken forward for next-generation targeted sequencing had additional CS1 (ACACTGACGACATGGTTCTACA) and CS2 (TACGGTAGCAGAGACTTGGTCT) Fluidigm tag sequences on the nested PCR primer forward and reverse, respectively. The amplicons were barcoded using Fluidigm Access Array Barcode Library for Illumina Sequencers (Cat#100-4876) by taking 0.8 µl multiplex PCR products (multiplex group A-C), 4 µl Fluidigm barcode primer (400 nM final concentration), 10 µl of 2X Phusion Master Mix (NEB, Cat#M0531L), and 3.6 µl H_2_O (20 µl reaction). The barcoding PCR reaction parameters were: one cycle at 98°C for 2 min, six cycles of 98°C for 10 s, 60°C for 30 s, and 72°C for 1 min, followed by one cycle at 72°C for 10 min. The products were run on 2% agarose gel and extracted using the QIAquick Gel Extraction Kit (Qiagen, Cat#28706). The purified products were submitted for paired-end 300 bp sequencing on MiSeq (Illumina), resulting in >1,000 coverage per gene.

### Lymphoid differentiation of transduced MLL-Af4 cord blood cells

*MLL*-*Af4* cord blood cells (36) were transduced with short hairpin constructs targeting CHD4, PHF3 or NTC control (as described in supplementary methods) and co-cultured with MS-5 stroma cells in lymphoid culture conditions. Single and triple transduced populations were identified using the construct specific fluorophores and lineage specific surface markers assessed as a proportion of the total transduced leukocyte population.

### MLL-Af4 stem cell expression analysis

Following myeloid or lymphoid culture of *MLL/Af4* transduced CD34+ cord blood cells, CD19+CD33-, CD19-CD33+ and CD19-CD33-populations were flow sorted, lysed and RNA extracted using RNeasy Micro Kit (Qiagen), according to the manufacturer’s instructions. Input RNA was equilibrated to a starting input cell number of 300 cells per population before cDNA and sequencing library production were performed using SMARTSeqv4 (Clontech) and NexteraXT (Illumina) kits, according to manufacturer’s instructions. The resultant libraries were submitted for paired-end 150 bp sequencing on a NEXTSeq500 (Illumina). For each sample, transcript abundance was quantified from raw reads with Salmon (version 0.8.2) using the reference human transcriptome (hg38) defined by GENCODE release 27. An R package Tximport (version 1.4.0) was used to estimate gene-level abundance from Salmon’s transcript-level counts. Gene-level differential expression analysis was performed using DESeq2 (version 1.16.1) prior NMF analysis as described above.

## Supporting information

Tirtakusuma_2021_Table S1

Tirtakusuma_2021_Table S2

Tirtakusuma_2021_Table S3

Tirtakusuma_2021_Table S4

Tirtakusuma_2021_Table S5

Tirtakusuma_2021_Table S6

Tirtakusuma_2021_Table S7

Tirtakusuma_2021_Table S8

Tirtakusuma_2021_Table S9

## Data availability

Exome sequencing data and genome sequencing data presented in this manuscript have been deposited in the NCBI Sequence Read Archive (SRA) under project numbers PRJNA547947 and PRJNA547815 respectively. Immunoglobulin and TCR sequencing data have been deposited in NCBI SRA under project number PRJNA511413. RNA sequencing data and DNase hypersensitivity sequencing data were deposited in Gene Expression Omnibus under accession numbers GSE132396 and GSE130142 respectively. All deposited data will be publically available following publication of the manuscript. Requests for additional specific data/materials should be made to Olaf Heidenreich.

## Acknowledgements

We thank Jon Coxhead and Raf Hussain at the Newcastle University core genomics facility as well as Marc van Tuil at the Princess Maxima Center Diagnostic department for development of sequencing strategies. We acknowledge the Newcastle University Flow Cytometry Core Facility and Tomasz Poplonski at the Princess Maxima Center Flow Cytometry core facility for their assistance with the generation of flow cytometry data and cell sorting strategies as well as the Newcastle University Bioinformatics Support Unit for helping to develop the analysis approach for sequencing data. We thank Ruben van Boxtel and Eline Bertrums at the Princess Maxima Center for the generation of single cell derived clones, derived from early progenitors sorted from MPAL patient samples. We thank Monique den Boer, Frank van Leeuwen and Ronald Stam for critically reading the manuscript.

This study makes use of data generated by the St. Jude Children’s Research Hospital – Washington University Pediatric Cancer Genome Project and the Therapeutically Applicable Research to Generate Effective Treatments (TARGET) initiative, phs000218, managed by the NCI (see supplementary methods).

## Author contributions

Conceptualization, O.H., S.B., C.B.;

Methodology, O.H., C.B., R.T., K.S., P.M., S.B., A.P., C.M., A.K., Z.K., J.B., V.B., M.H., R.M., J.V., J.M.A., S.L.;

Software Programming, S.N., J.H.K., V.V.G., A.K., D.W., Pi.C.;

Formal Analysis, S.N., J.H.K., V.V.G., A.K., Z.K., J.B., D.W., Pi.C., C.B., O.H.;

Investigation, R.T., K.S., P.M., A.P., C.M., H.J.B., A.K., S.A., M.R.I., P.E., H.M., A.E., N.M.S., S.E.F., Y.S., D.P., Pi.C.;

Resources, F.V., E.Z., S.L., J.S., A.S., J.C.M., L.J.R., C.E., O.A.H., S.Ba., R.S., N.M., M.C., V.B., R.M., M.W., C.J.H., C.A.C., Pe.C., M.H., D.S., Y.O., M.J.T., P.N.C., J.C.M., C.B., O.H.;

Data Curation, S.N., D.W., Pi.C.; Writing, S.B., O.H., C.B., R.T., K.S.;

Supervision, O.H., S.B., J.M.A., J.V., C.B., ;

Funding Acquisition, O.H., J.V., S.B., C.B., P.N.C., J.M.A., E.Z.

