## Supplementary methods and figures for "Epigenetic regulator genes direct lineage switching in *MLL-AF4* leukaemia"

**Running title:** Lineage switching in *MLL*-*AF4* leukaemias

**Keywords:** *MLL*-*AF4*; *KMT2A*-*AFF1*; acute lymphoblastic leukaemia; nucleosome remodelling and deactylation complex (NuRD); chromatin remodelling

### Corresponding authors

Dr Simon Bomken

Wolfson Childhood Cancer Research Centre

Translational and Clinical Research Institute

Level 6 Herschel Building

Brewery Lane

Newcastle University

Newcastle upon Tyne

NE1 7RU, UK

E mail:

Professor Olaf Heidenreich

Princess Maxima Center for Pediatric Oncology

Heidelberglaan 25

3584 CS Utrecht

The Netherlands

E mail:

### Supplements – Table of contents

**Supplementary methods:** pages 4-12

**Supplementary references:** pages 13

**Supplementary figures 1-8:** pages 14-25

### Supplementary methods

#### Cell Lines

SEM, 697, REH and RS4;11 cells were grown in RPMI-1640 with 10% fetal bovine serum (FBS, Sigma) ] at 37°C in a humidified 5% CO2 incubator. All cell lines contain no PHF3, CHD4 (with exception of the REH cell line - missense mutation at p.R61Q), PCGF6, AUTS2 mutation according to the CCLE (<https://portals.broadinstitute.org/ccle>) database.

*MLL-Af4* transduced cord blood cells (1) were cultured in IMDM with 10% FBS, 2 mM Glutamine supplemented with recombinant human SCF, IL-3, IL-6, FLT-3L, and TPO (10 ng/ml each) at 37°C in a humidified 5% CO_2_ incubator. To prime towards lymphoid differentiation, the cells were co-cultured with MS-5 (RRID:CVCL_2128, murine) stroma cells in α-MEM with 10% FBS, 2 mM glutamine supplemented with SCF, FLT-3L, and IL-7 (10 ng/ml each). The cells were demi-populated weekly.

#### In Vivo Mouse Studies

*In vivo* studies were conducted in accordance with the UK Animals (Scientific Procedures) Act 1986 under project licences PPL60/4552 and PPL60/4222 following institutional ethical review.

NOD.Cg-*Prkdc^scid^ Il2rg^tm1Wjl^*/SzJ (NSG, Charles River Labs and bred in-house) mice (both sexes) aged 8 -10 weeks old were transplanted intra-femorally under isoflurane anaesthesia with 10^6^ cells from sample LS01PALL. Sample LS01RAML was transplanted into 5 NSG mice without evidence of engraftment.

MISTRG mice, expressing four human cytokines (M-CSF^h^; IL-3/GM-CSF^h^; hSIRPA^tg^;TPO^h^;Rag2^-^;γc^-^, Jax Lab (2), bred in house) were transplanted intra-hepatically. Three day old mice (n = 2) were sub-lethally irradiated (1.5 Gy) 24 h before the transplantation. Each mouse received 10^6^ mononuclear cells from sample LS01RAML.

Mice were humanely killed before or when they displayed end points specified by the licenses (tumours reached 1.5 cm in diameter, lost >10% weight in 3 consecutive days or 20% at any time, or displayed signs of ill). PDX cells were harvested from engrafted spleen (NSG) or abdominal masses (MISTRG).

#### Flow cytometry and cell sorting

One million cells (of all tested cell lines, cord blood or LS03ALL PDX cells) were used for flow cytometry analysis of CD19 (BV421) and CD33 (APCCy7). The antibody details are listed in the Key Resources Table. Cells were collected and washed in a 3 ml wash buffer containing 0.2% BSA in PBS, re-suspended in 100 µl wash buffer and 5 µl each antibody was added, followed by incubation at RT for 20 min in the dark. Cells were washed and re-suspended in a 0.5 ml wash buffer. Flow cytometry was performed on FACSCanto II from Becton Dickinson (BD) and data analysed with FlowJo (Treestar).

Haematopoietic hierarchy analysis was performed from 10^7^ cells of primary and PDX samples. The cells were collected and incubated for 30 min in a wash buffer containing 0.5% FBS, 2 mM EDTA in PBS. Following the incubation, the cells were washed and resuspended in a wash buffer. Fluorescence activated cell sorting (FACS) was performed using a FACS Aria Fusion (BD) or ASTRIOS EQ (Beckman Coulter). In order to separate early progenitors or lymphoid and myeloid fractions, MPAL patient samples were sorted from 10^7^ or 3*10^6^ cells for MPAL1 and MPAL2 respectively. Sorted cells were collected into 1.5 ml microfuge tubes containing 500 µl RPMI with 10% fetal calf serum or SFEM II media in case of cell lines or primary samples, respectively. In order to analyse single cell derived clones, sorted early progenitor cells derived from MPAL samples were deposited onto 384-well plates containing 75ul of SFEM II media (supplemented with SCF, FLT3L, IL6, IL3 and TPO). Single cells were grown into 60-90% confluency colonies followed by RNA/DNA extraction and MLL-AF4 detection by PCR.

#### Single cell sorting and whole genome amplification (WGA)

Single cells were labelled (see flow cytometry and cell sorting section above) and sorted into 96 well plate (Eppendorf twin.tec PCR Plate 96 full skirt) using FACSAria Fusion (BD). The plates were filled with 3.5 µl PBS/well prior to the sorting. Each plate included forty-five single cell samples, two 0 cells as negative control, and one bulk (300 – 1,000 cells) as positive control sample. The cells were prepared from as many as 10^5^ cells in 500 µl buffer (0.5% FBS, 2 mM EDTA in PBS).

Cell sorting was performed using the “single cell” setting on FACSAria Fusion. Following the sorting, the plate was centrifuged at 900 g for 1 min, snap-frozen on dry ice, and then kept at -20°C until the whole genome amplification (WGA) procedure.

WGA was performed using REPLI-g Single Cell Kit (Qiagen, Cat#150345). The cells were lysed by adding 1.5 µl Buffer D2 and incubation at 65°C for 10 min in a thermal cycler (HYBAID PCRExpress). Subsequently, the reaction was terminated by adding 1.5 µl Stop Solution and stored on the ice. The mixture of 20 µl REPLI-g sc Reaction Buffer and REPLI-g sc DNA Polymerase was added and incubated at 30°C for 8 h. The reaction was terminated by heat inactivation at 65°C for 3 min. The products were diluted 100 fold with TE Buffer ahead of PCR amplification.

#### Long-distance inverse PCR experiments

*MLL* rearrangement sequences were identified using long-distance inverse PCR (LDI-PCR) as previously described (3, 4).

#### PCR-based clonality immunoglobulin gene targets

Clonality analysis based on immunoglobulin gene rearrangements was performed according to the BIOMED-2 Concerted Action BMH-CT98-3936 using standardised protocols and primer sets (5). Duplicate samples were analysed by GeneScan analysis on a Life Technologies 3100 platform.

#### Non-negative matrix factorization analysis

Non-negative matrix factorisation (NMF) was performed using data sets obtained with permission. Data from Bolouri *et al.* (6) were generated by the Therapeutically Applicable Research to Generate Effective Treatments (TARGET) initiative, phs000218,
managed by the NCI. The data used for this analysis are available [<https://www.ncbi.nlm.nih.gov/projects/gap/cgi-bin/study.cgi?study_id=phs000218.v21.p7>]. Information about TARGET can be found at <http://ocg.cancer.gov/programs/target>. Data from Andersson *et al.* (7) was obtained with permission of the Pediatric Cancer Genome Project from the European Genome-Phenome Archive (EGA study accession EGAS00001000246). Data from Novoshtern *et al.* (8) was obtained openly from Gene Expression Omnibus (GEO dataset GSE24759). NMF was used to extract metagenes from three leukaemic gene expression datasets (6, 7, 8). A matrix of gene expression data combining per gene read counts of samples from each dataset was created, normalized and variance-stabilized transformed (vst) using DESeq2. Batch effect between leukaemia derived datasets (6, 7) was minimized using the remove unwanted variation method from RUVSeq (9) on upper-quartile normalized counts whilst retaining variation associated with biological covariates of interest. To remove genes that did not vary sufficiently across the model dataset, for RNAseq data, we capped expression values at 20 and 10,000 read counts and only included genes with > 10-fold and >1000 read counts between minimum and maximum. For the Affymetrix U133A microarray, we capped expression values at 0.01 and 10,000 units and only probes with > 3-fold and >5 units difference between minimum and maximum were considered in the NMF analysis. To identify the most robust combination of metagenes and subgroups/clusters in the model datasets, consensus bootstrapped NMF clustering was performed on filtered gene count matrix as previously described (10). Briefly, we performed NMF and K-means clustering, testing all combinations of 2-15 metagenes and clusters with bootstrapped resampling method (n=100) to test for reproducibility. Cluster stability measures (Cohen’s kappa, average silhouette scores) were assessed to determine optimal combinations of metagenes and clusters. Samples assigned to the same cluster fewer than 90% of replicates were removed from the dataset. All genes in the refined model dataset were then column rank normalised and metagenes scores were recalculated.

Metagenes were projected onto our RNAseq datasets (both lineage switch cases and *MLL-Af4* cord blood samples) by the pseudoinverse method as outlined in (11, 12). All analysis and figures were generated using R/Bioconductor and by using a modified version of the NMF scripts provided by (11).

#### Reverse Engineering of Transcriptional Networks

A mutual information network was developed such that the connections between genes within the network represented an inferred likelihood of a causal relationship within ALL/AML. The network was further refined by retaining only those nodes (genes) and edges (causal connections) which are significantly associated, in expression terms, to the distinction between normal myeloid and lymphoid cells. Thus, the centrality of the mutations within the network reflects the estimated extent of their causal influence upon a lymphoid/myeloid distinction within the framework of primary ALL/AML.

Mutual Information networks were reverse engineered from 216 published AML/ALL Affymetrix HGU133p2 expression profiles (GSE11877 & GSE24006) processed using Rma (affy package R/Bioconductor) using the ARACNe2 algorithm (13) set to adaptive partitioning with a DPI tolerance of 0.1 and a p-value threshold of 0.01 and 1000 bootstraps. The probes/genes were filtered to remove those that were insufficiently variable as per the Aracne method. Probes had to be greater than 3 fold expression and 300 delta between the min and max excluding the 5% most extreme values to be included. Probes without >20 intensity in greater than 20% of samples were also removed. All probes which fulfilled these requirements were included in the construction of this network (n=9780).

#### Nodes within the network were annotated with a metagene value calculated using Non-Negative Matrix Factorisation (k=2) and reflecting the differences between myeloid and lymphoid cells (GSE24759) and the other. A metagene cutoff of 0.13 (based on a calculated alpha of 0.1) was applied to trim the network. Cytoscape (14) was used to merge nodes where probe sets represented common genes and to calculate centrality statistics for each of the mutated genes of interest. Nodes of interest (i.e. mutated nodes) were ranked according to their centrality (e.g. degree). Nodes were sized and coloured according to their differential expression between AML and ALL types; size = significance, colour = log2 fold change.

#### PHF3, CHD4, PCGF6, and AUTS2 shRNAs

shRNAs against PHF3 (TRCN0000019118, TRCN0000019114, TRCN0000274376), CHD4 (TRCN0000380981, TRCN0000021363, TRCN0000021360), PCGF6 (TRCN0000229804, TRCN0000073109, TRCN0000073109), AUTS2 (TRCN0000119058, TRCN0000304019, TRCN0000304081), or non-targeting control (ATCTCGCTTGGGCGAGAGTAAG) were cloned into pLKO5d.SFFV.miR30n (15). Each target gene was linked with different fluorescent protein, including dTomato (PHF3 and AUTS2), eGFP (CHD4 and PCGF6), and RFP657 (non-targeting control).

The vector contained BsmBI at the cloning sites. The oligonucleotides were designed with the appropriate complementary overhang sequences of BsmBI-cleaved vector. They were ligated and transformed into STBL3 chemically competent cells (Invitrogen) according to manufacturer’s instructions and plated on agar plates containing 100 µg/ml ampicillin incubated for 16 h at 37°C. Single colonies were inoculated for DNA preparation, and extracted using EndoFree Plasmid Maxi Kit (Qiagen, Cat#12362). All clones were verified by sequencing prior to lentivirus production.

#### Lentivirus Production and Transduction

Viral particles were produced using calcium phosphate precipitation method on 293T cells.The cells were grown in 100 mm tissue culture dishes at a concentration of 1-2 x 10^6^ cells in 10 ml medium the day prior to co-transfection. Equimolar amounts of envelope plasmid pMD2.G, packaging plasmid pCMVΔR8.91, and the shRNA vector pLKO5d.SFFV.miR30n were mixed and the volume adjusted with HEPES buffer solution (2.5 mM HEPES containing deionized water at pH7.3) to 250 µl. A volume of 250 µl of 0.5 M CaCl_2_ was added to the mixture. This solution was added to 500 µl of 2X HeBS (0.28 M NaCl, 0.05 M HEPES and 1.5 mM Na_2_HPO_4_ in deionized water at pH 7.00) and mixed by vortexing. It was incubated at RT for 30-40 min to allow the formation of the calcium phosphate precipitate, before adding dropwise on the 293T cells. After 16 h, the cells were washed with 10 ml PBS and added with 10 ml culture media. They were incubated at 37°C in a humidified atmosphere with 5% CO_2_ for the next two days. Lentivirus particles were collected by centrifuging the supernatant at 400 g for 10 min at 4°C and filtered through Acrodisc Syringe 0.45 µm filters. They were stored in aliquots at -80°C. Cell lines and PDX samples were transduced with lentivirus as previously described (16).

#### qRT-PCR

One million cells were collected and RNA was extracted using RNeasy Mini Kit (Qiagen, Cat#74106). cDNA was synthesised from 1 µg RNA using RevertAid H Minus First Strand cDNA Synthesis Kit (Thermo Fisher Scientific, Cat#K1632 ). The product was diluted by adding 80 µl H_2_O (17). Sequences of primers used in this study were designed using Primer Express (Applied Biosystems) software.

List of antibodies and oligonucleotides used in the study.

| REAGENT or RESOURCE | SOURCE | IDENTIFIER |
| --- | --- | --- |
| **Antibodies** |  |  |
| CD3-APC-H7 | BD Biosciences | Cat#641397; RRID:AB_1645731 |
| CD3-FITC | BD Biosciences | Cat#345763 |
| CD3-BV650 | Biolegend | Cat#300468 |
| CD10-BV650 | BD Biosciences | Cat#563734; RRID:AB_2738393 |
| CD11b-APC/Fire750 | Biolegend | Cat#301351 |
| CD14-FITC | BD Biosciences | Cat#555397; RRID:AB_395798 |
| CD14-BV605 | Biolegend | Cat#367125 |
| CD16-FITC | BD Biosciences | Cat#335035 |
| CD19-PE-CF594 | BD Biosciences | Cat#562294; RRID:AB_11154408 |
| CD19-APCCy7 | Biolegend | Cat#363010; RRID:AB_2564193 |
| CD19-BV421 | Biolegend | Cat#302234; RRID:AB_11142678 |
| CD19-PECy7 | Biolegend | Cat#363011 |
| CD20-PE | BD Biosciences | Cat#345793 |
| CD20-FITC | Biolegend | Cat#302303 |
| CD33-APC | BD Biosciences | Cat#345800 |
| CD33-APCCy7 | Biolegend | Cat#366614; RRID:AB_2566416 |
| CD33-BV421 | Biolegend | Cat#303416; RRID:AB_2561690 |
| CD34-PerCPCy5.5 | BD Biosciences | Cat#347222 |
| CD34-APCCy7 | Biolegend | Cat#343514; RRID:AB_1877168 |
| CD34-APC | Biolegend | Cat#343607 |
| CD38-PeCy7 | BD Biosciences | Cat#335825 |
| CD38-PE | Biolegend | Cat#303506 |
| CD45RA-BV510 | Biolegend | Cat#304142; RRID:AB_2561947 |
| CD56-FITC | BD Biosciences | Cat#345811 |
| CD64-BV711 | Biolegend | Cat#305041 |
| CD90-A700 | Biolegend | Cat#328120; RRID:AB_2203302 |
| CD90-PerCPCy5.5 | Biolegend | Cat#328118; RRID:AB_2303335 |
| CD90-BV421 | Biolegend | Cat#328122 |
| CD117-PE | BD Biosciences | Cat#332785 |
| CD117-BV605 | BD Biosciences | Cat#562687; RRID:AB_2737721 |
| CD123-BV421 | BD Biosciences | Cat#306018; RRID:AB_10962571 |
| HLA-DR-BV786 | Biolegend | Cat#307642; RRID:AB_2563461 |
| HLA-DR-A700 | Biolegend | Cat#560743; RRID:AB_1727526 |
| HLA-DR-PerCPCy5.5 | Biolegend | Cat#307629 |
| **Oligonucleotides** |  |  |
| shPHF3-1 | GPP, Broad Institute | TRCN0000019118 |
| shPHF3-2 | GPP, Broad Institute | TRCN0000019114 |
| shPHF3-3 | GPP, Broad Institute | TRCN0000274376 |
| shCHD4-1 | GPP, Broad Institute | TRCN0000380981 |
| shCHD4-2 | GPP, Broad Institute | TRCN0000021363 |
| shCHD4-3 | GPP, Broad Institute | TRCN0000021360 |
| shPCGF6-1 | GPP, Broad Institute | TRCN0000229804 |
| shPCGF6-2 | GPP, Broad Institute | TRCN0000073109 |
| shPCGF6-3 | GPP, Broad Institute | TRCN0000073109 |
| shAUTS2-1 | GPP, Broad Institute | TRCN0000119058 |
| shAUTS2-2 | GPP, Broad Institute | TRCN0000304019 |
| shAUTS2-3 | GPP, Broad Institute | TRCN0000304081 |
| shNTC (ATCTCGCTTGGGCGAGAGTAAG) | (16) | N/A |
| PHF3 Fw (ATGGACCTGGGCTTGAACTG) | This paper | N/A |
| PHF3 Rev (TGGTGGTGCACTTTCAGGAG) | This paper | N/A |
| CHD4 Fw (TGCTGACACAGTTATTATCTATGACTCTGA) | This paper | N/A |
| CHD4 Rev (ACGCACGGGTCACAAACC) | This paper | N/A |
| PCGF6 Fw (GGGAAATCCGACGTGCAAT) | This paper | N/A |
| PCGF6 Rev (GGAGAAACCACAAGACCATAATGA) | This paper | N/A |
| AUTS2 Fw (AAAAGGACCCGAGGTTGACA) | This paper | N/A |
| AUTS2 Rev (GCGATGTGAACATGCATAGCA) | This paper | N/A |
| GAPDH Fw (GAAGGTGAAGGTCGGAGTC) | This paper | N/A |
| GAPDH Rev (GAAGATGGTGATGGGATTTC) | This paper | N/A |
| CD19 Fw (TGACCCCACCAGGAGATTCTT) | This paper | N/A |
| CD19 Rev (CACGTTCCCGTACTGGTTCTG) | This paper | N/A |
| CD33 Fw (CTCGTGCCCTGCACTTTCTT) | This paper | N/A |
| CD33 Rev (CCCGGAACCAGTAACCATGA) | This paper | N/A |
| CSF3R Fw (CCCAGGCGATCTGCATACTT) | This paper | N/A |
| CSF3R Rev (AACAAGCACAAAAGGCCATTG) | This paper | N/A |
| KIT Fw (GGACCAGGAGGGCAAGTCA) | This paper | N/A |
| KIT Rev (GATAGCTTGCTTTGGACACAGACA) | This paper | N/A |

### Supplementary figures

**Figure S1. Genomic characterization of the *MLL-AF4* lineage switch cases.** (A) Sequencing of *MLL-AF4* fusion breakpoints from DNA (LS02, LS03, LS04, LS07, LS08, LS09) or RNA (LS05, LS06, LS10, MPAL1, MPAL2). (B) Whole genome sequencing data of LS01 showing karyotype (outer circle), copy number changes (log2 depth ratio in 1Mb windows, loss <2 green dots, gain >2 red dots) and structural variants (translocations - red connecting lines, deletions – blue lines, inversions, purple lines). (C) Schematic representation of identified fusion variants, located within the major *MLL* breakpoint region, present in analysed t(4;11) cases (detailed breakpoint description presented at Fig.S1 and Table S1). Red connecting line indicates MLL-AF4 translocation positions of each gene partner.

**Figure S2. Cellular origin of the *MLL-AF4* leukaemia.** (A) Immunoglobulin rearrangement on ALL and AML LS01 according to the standard protocol BIOMED-2. Clonal peaks are coloured in dark blue, as indicated in the multiplex amplification FR1, FR2, and FR3 of the VH segments LS01PALL (left panel). The multiplex results were confirmed by single amplification VH-JH primer sets, which further showed two clonal VH3, one clonal VH1, and clonal VH4 rearrangement. No clonal peaks are seen in LS01RAML (right panel). (B) Presentation ALL sample only produces ALL in all mouse strains transplanted. In contrast, relapse AML sample does not engraft in lymphoid supportive NSG mice and produces only AML in myeloid supportive MISTRG mice. (C) Post-transplantation flow cytometric analysis of LS01PALL produced in NSG mice (top panel) and LS01RAML produced in MISTRG mice (bottom panel).

**Figure S3. Evaluation of *MLL-AF4* presence in HSPC populations.** (A) Flow cytometry plots of single sorted HLA-DR+CD14+CD11c+ monocytes LS01PALL. The two *MLL-AF4* positive cells, cells 11 and 18, are highlighted in red throughout the sorting strategy (lower panel). (B) Amplification of the *MLL-AF4* in cells 11, 18 and bulk LS01 sample shows the expected 299 base pair band. (C) Flow cytometry plots showing sorting strategy for lymphoid and myeloid populations of MPAL1 and MPAL2. (D) Bulk sort plots for HSC and MPP populations (upper panel) and single cell plots for Lin-CD34+ populations (lower panel) sorted from MPAL1 sample. (E) Amplification of the *MLL-AF4* in single- cell derived clones Lin-CD34+, lymphoid, and myeloid fractions of MPAL1 patient.

**Figure S4. Transcriptional profiles of ALL and AML lineage switch leukaemia.**
(A) Non-negative matrix factorisation analysis of paired presentation ALL cases and relapse AML cases, against normal haematopoietic precursors (Novershtern *et al*, 2011). Presentation ALL cases demonstrate a high expression of B cell metagene signature, whilst relapse AML cases continue to express this signature to a variable degree (upper panel). Relapse AML cases demonstrate high expression of myeloid metagene signature, at a comparable level to that of GMP (lower panel). (B) Non-negative matrix factorisation analysis of paired presentation ALL cases and relapse AML cases, against ALL and AML cases, with and without *MLL* rearrangement (Andersson *et al* 2015, Bolouri *et al* 2018). Heatmap shows clustering of presentation cases (dark blue) with B-ALL *MLL-AF4* and relapse cases (dark green) with AML with *MLL*r. (C) Differential promoter accessibility is associated with higher expression of *PAX5*, *LEF1*, and *CD79A* in presentation ALL cases, and of *CSF3R*, *KIT* and *CSF2RA* in relapse AML cases. TSS – transcriptional start site. (D) Gene set enrichment analysis indicates impaired DNA repair and cell cycle progression in AML relapse. NES – normalised enrichment score.

**Figure S5. Impact of leukaemia lineage switch on *MLL-AF4*-regulated genes.** (A) HOXA cluster analysis includes a general reduction on HOXA3-10 in relapse AML cases. (B) Differential chromatin accessibility and expression across the HOXA cluster at ALL presentation and AML relapse of case LS01, demonstrating differential DNase hypersensitivity of HOXA cluster. (C) Differential expression across the HOXA cluster in five cases of lineage switch from ALL (upper panel for each case) and AML (lower panel). (D) The list of direct *MLL-AF4* target genes obtained by overlaying SEM cells and t(4;11) patient cells ChIPseq data (Kerry *et al.*, 2017) and differentially expressed genes in our lineage switch and MPAL cases (Venn diagram, upper panel). Highly enriched genes are further listed on the lower panel. (E) GSEA showing loss of chromatin modifying enzymes signature in AML relapse.

**Figure S6. High resolution DNase hypersensitivity sequencing.** (A) UCSC genome browser screenshot for *RUNX1* focused on an AML-associated DHS with C/EBP occupation as indicated by high resolution DHS-seq and Wellington analysis. FP - footprint. (B) Heat maps showing distal DHS regions specific for ALL presentation on a genomic scale. Red and green indicate excess of positive and negative strand cuts, respectively, per nucleotide position. Sites are sorted from top to bottom in order of decreasing Footprint Occupancy Score. *De novo* motif discovery in distal DHSs unique to ALL as compared to AML relapse, as shown on the table, right panel.

**Figure S7. Alternative splicing analysis.** (A) RNAseq read counts across all analysed patient samples. (B) Qualitative analysis of exon-exon spanning reads from LS01, LS03 & LS04 samples. Upper panel represents distribution of the unfiltered and unnormalized reads, lower panel represents filtered and voom normalized reads. Expression (abundance) of the exon-exon junctions is presented as log2-transformed counts per million (CPM). (C) Abundance of non-differential versus differential exon-exon junctions in the transcriptome of LSALs (left panel) and MPAL series of samples (right panel). According to limma/diffSplice approach, junctions were divided into non-differential (upper panel) and differential (bottom panel) splicing events. In each boxplot, the horizontal line represents the median of distribution, box shows the interquartile range, and whiskers are the minimum and maximum. p-values were calculated using the two-sided Mann-Whitney U test. (D) Venn diagrams showing distribution of identified retained intron (RI) across both LSAL and MPAL patients with corresponding GO terms describing cellular components affected in the AML relapse of lineage switched patients. (E) Venn diagrams showing distribution of identified differential exon-exon junctions (DEEj), significantly expressed in the AML relapse or myeloid compartment of MPAL patients with corresponding GO terms describing affected Reactome pathways and cellular components. GO terms analysis has been performed with <https://biit.cs.ut.ee/gprofiler/gost> under the highest significance threshold, with multiple testing correction (g:SCS algorithm).

**Figure S8. The functional role of epigenetic modifiers on leukaemia lineage switching.** (A) PHF3 scheme; the K1119I mutation is located at a highly conserved residue. An * (asterisk) indicates positions which have a single, fully conserved residue, a : (colon) indicates conservation between groups of strongly similar properties - scoring > 0.5 in the Gonnet PAM 250 matrix, a . (period) indicates conservation between groups of weakly similar properties - scoring < 0.5 in the Gonnet PAM 250 matrix. (B) Mutual information sub-networks illustrate high centrality of mutated genes *PHF3* (left panel) and *CHD4* (right panel) within the AML/ALL transcriptional network. Size and colour of nodes reflect differential expression between AML and ALL. (C) Myeloid marker CD33 expression level upon knockdown of *CHD4*, *PHF3*, and non-targeting control (NTC) in *MLL*r (t(4;11)) cell line RS4;11 and non-*MLL*r cell lines, 697 and REH cells. (D) CD19 and CD33 surface expression level change upon knockdown of *CHD4* and *PHF3* in the PDX sample generated from the first relapse of patient LS03 (ALL relapse). (E) Protein network analysis of differentially expressed NuRD and PRC1 members, showing interactions between proteins (and clusters, depicted as blue and red colour); interaction strength is shown as increasing thickness of the joining line (<https://string-db.org/>). (F) Fold change expression level changes of PRC1 members following lineage switched relapse and in MPAL cases. (G) Expression of lineage specific surface markers following *PCGF6* knockdown in SEM cells. Knockdown of *PCGF6* results in upregulation of *CD33*, *CSFR3* and *KIT* mRNA (upper panel). Surface CD33 expression increases substantially with shPCGF6_1 and shPCGF6_3, compared with shNTC (lower panel). (H) GSEA showing increased expression of PRC2 target genes and impaired function of PRC1 complex in the relapse AML samples. (I) Boxplot of NMF projections of CD19+ cord blood (CB_19pos) and CD33+ cord blood (CB_33pos) populations relative to the derived AML with *MLL*r metagene (red boxes) and B precursor ALL with *MLL*r metagene (turquoise boxes).

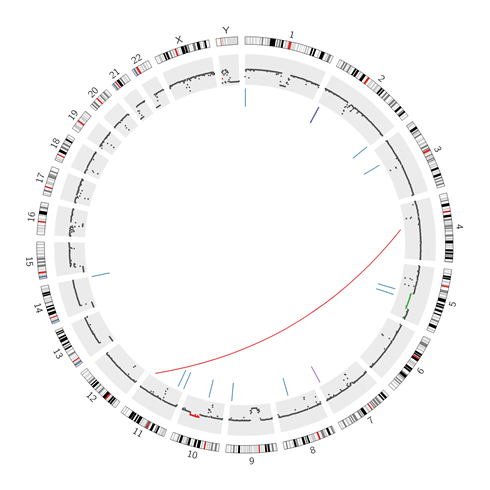

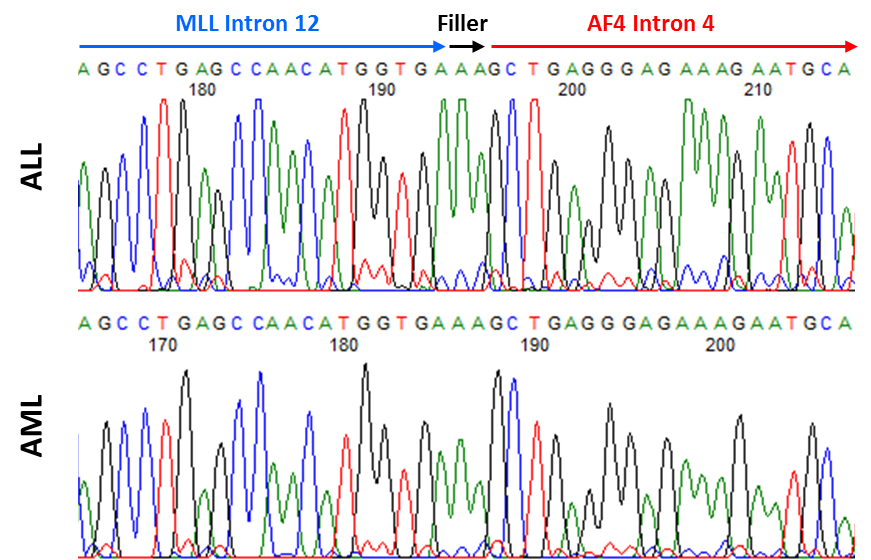

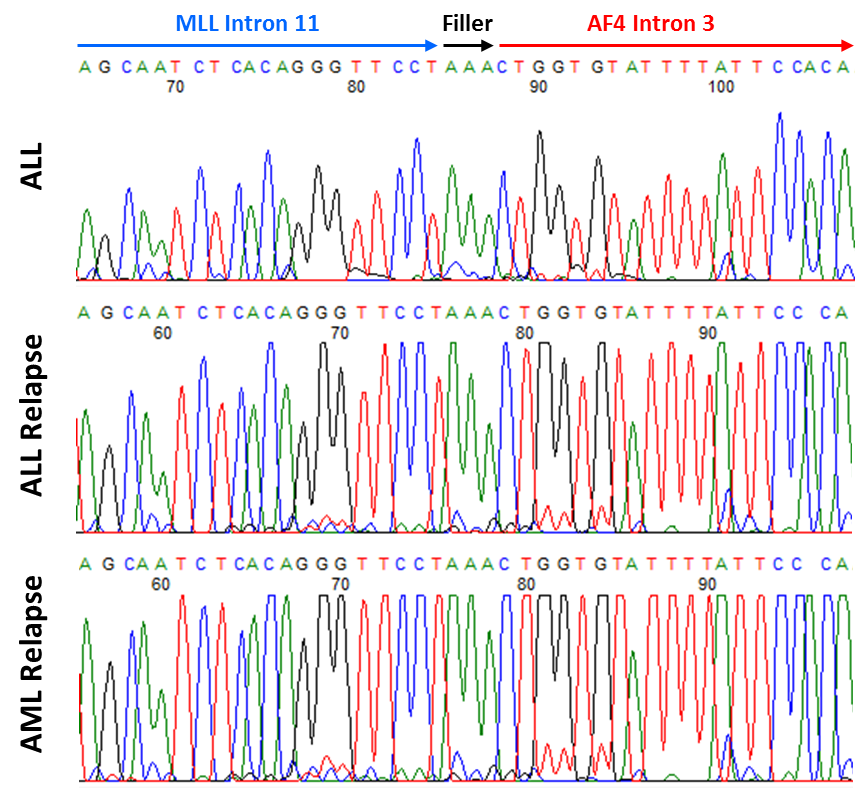

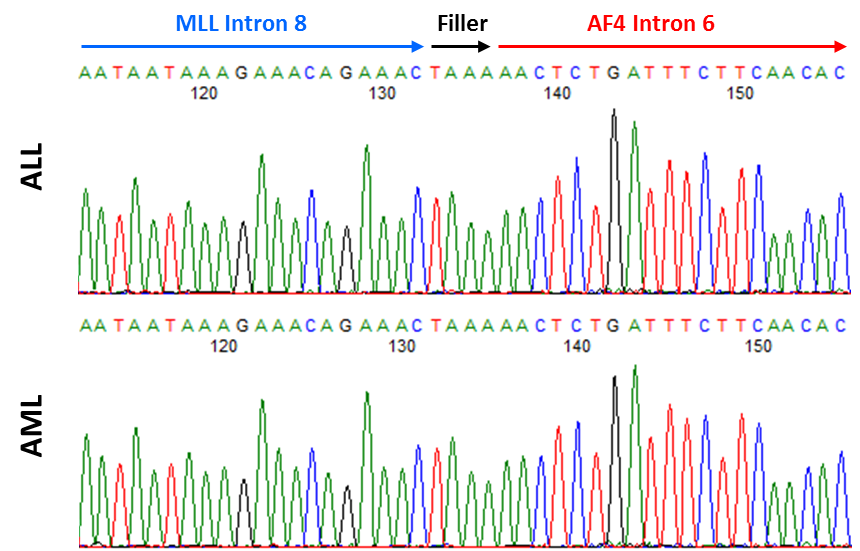

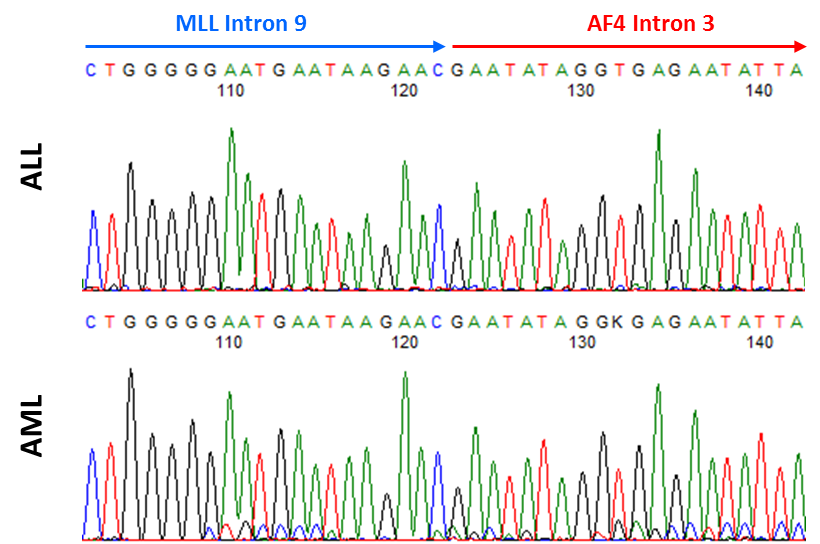

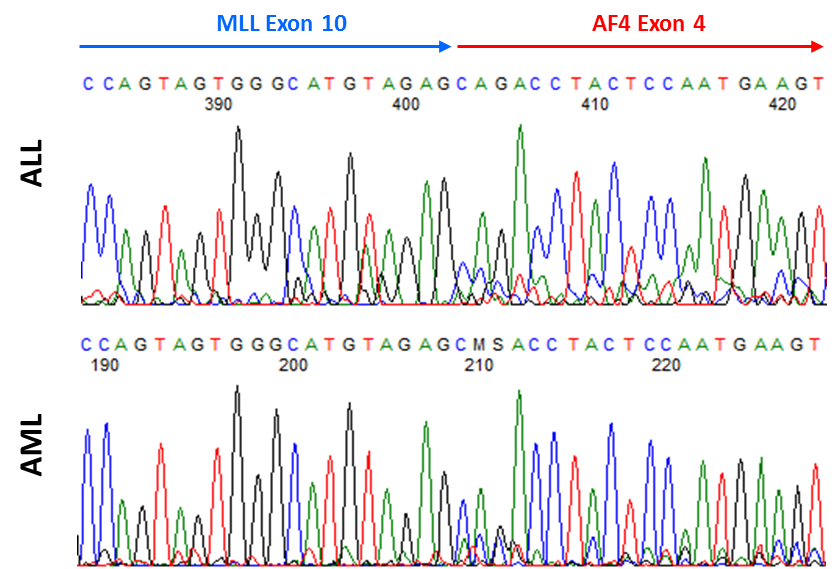

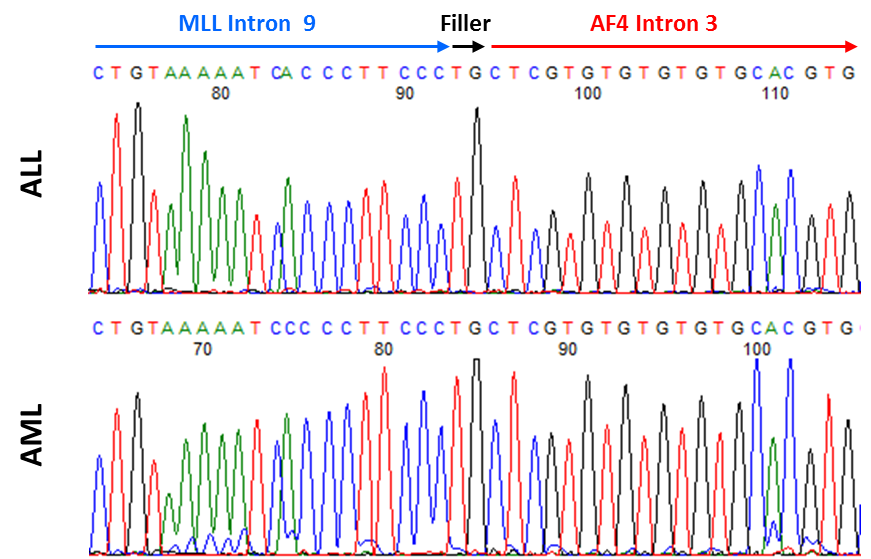

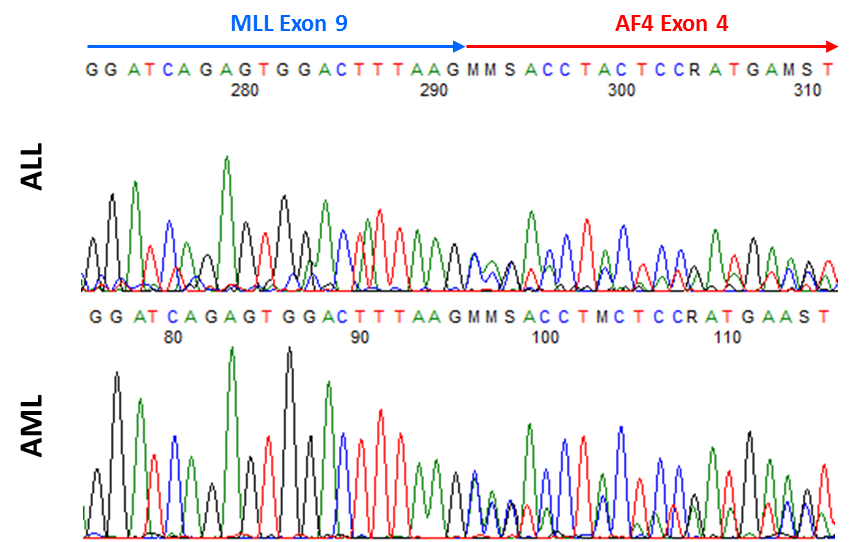

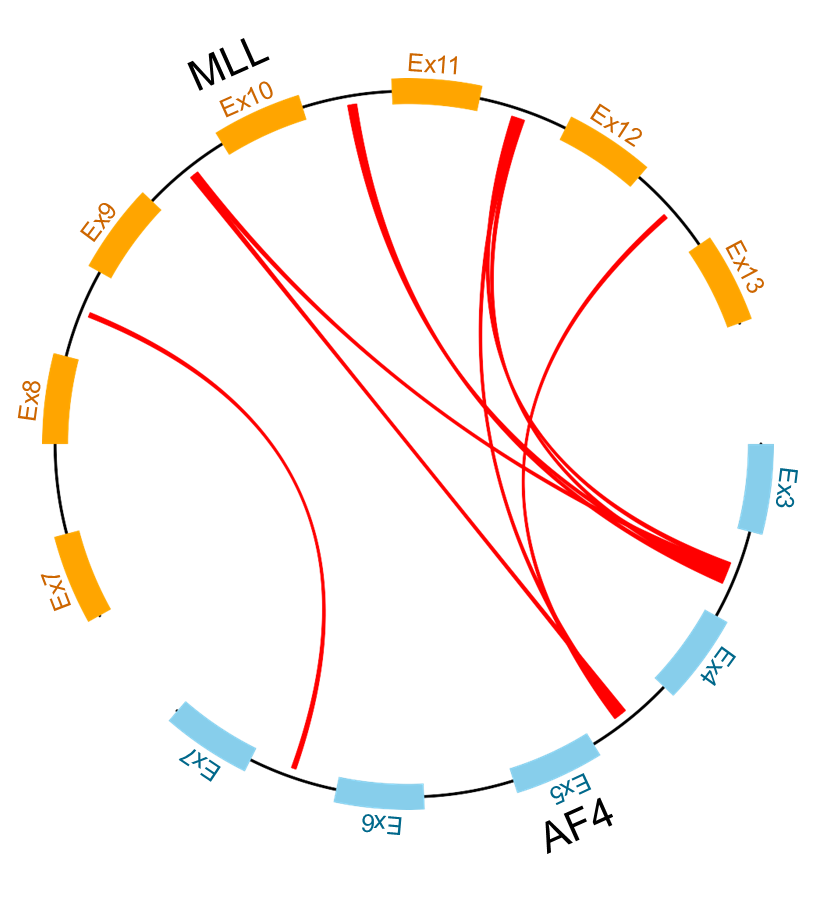

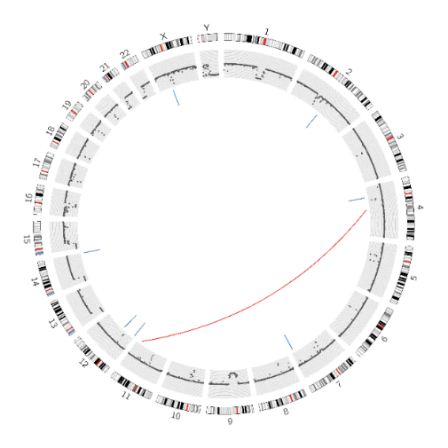

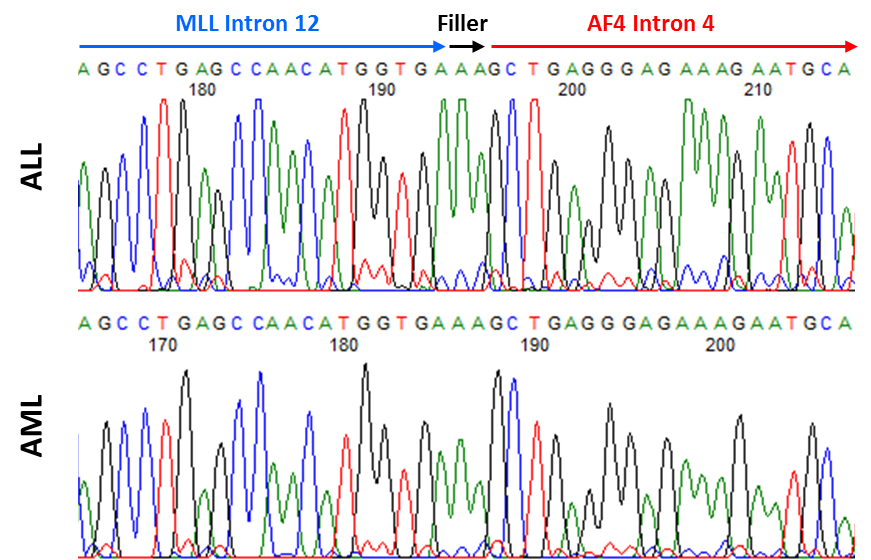

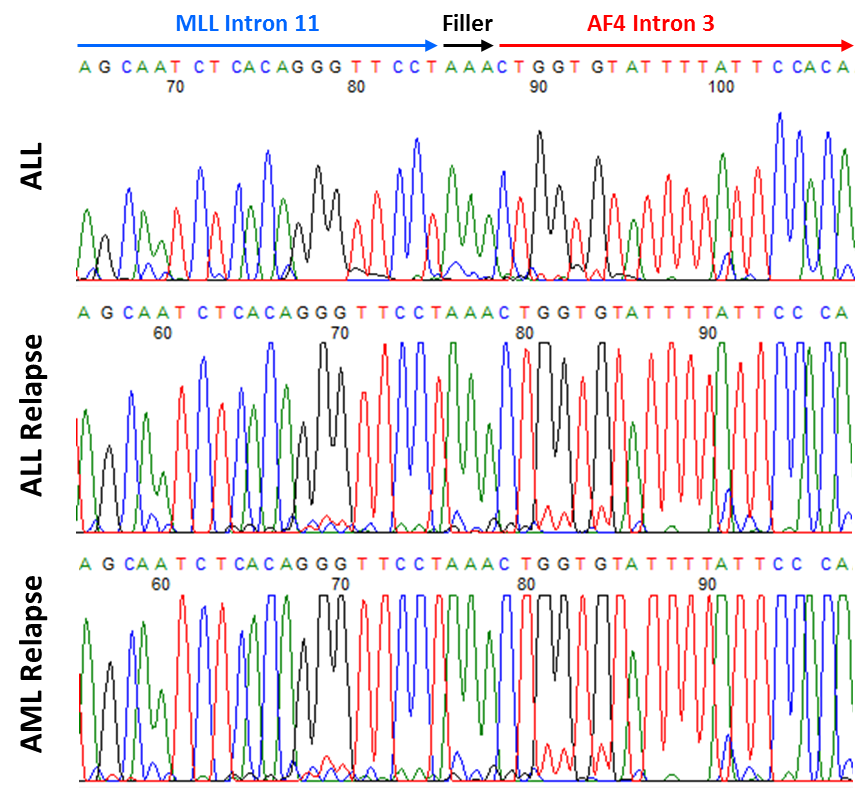

**Relapse AML**

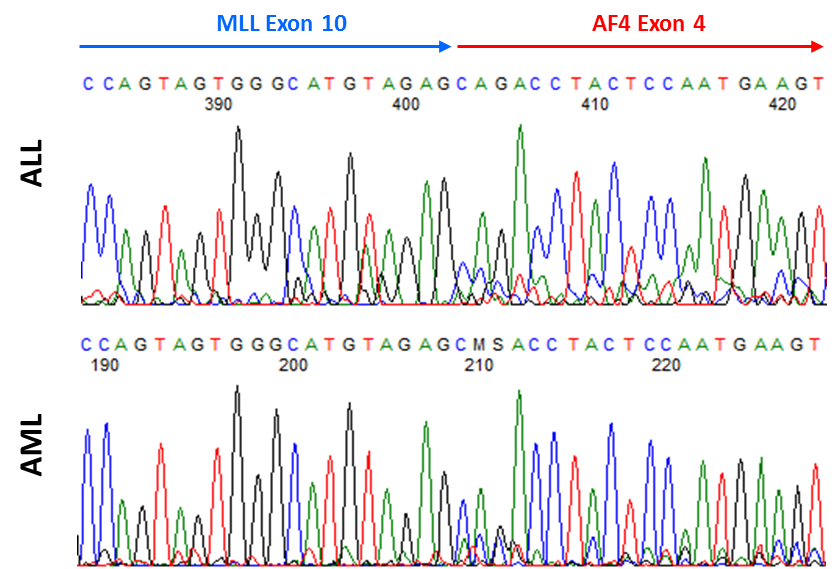

**ALL**

**AML**

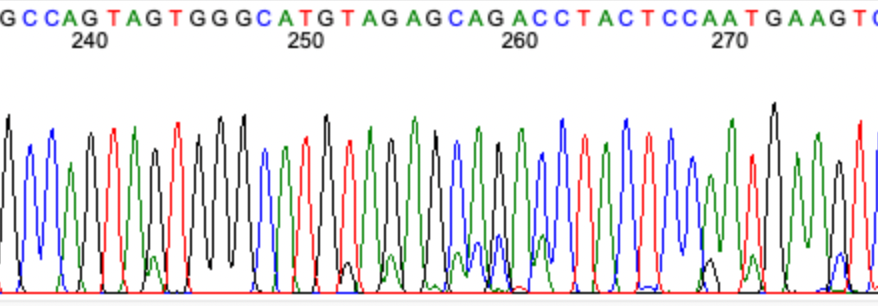

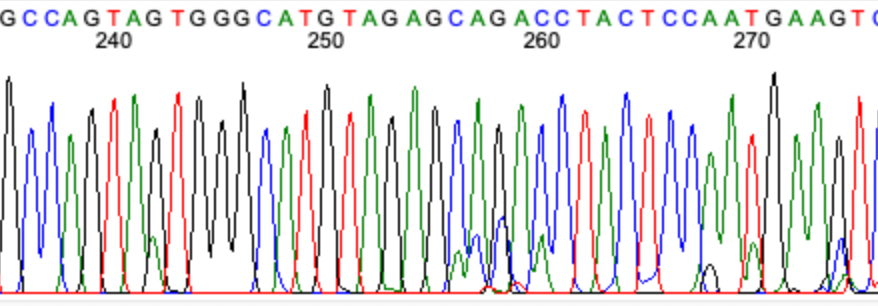

LS10

LS04

LS07

LS09

LS06

MPAL1

LS08

LS05

C

**Presentation ALL**

LS03

LS02

Figure S1

MPAL2

A

B

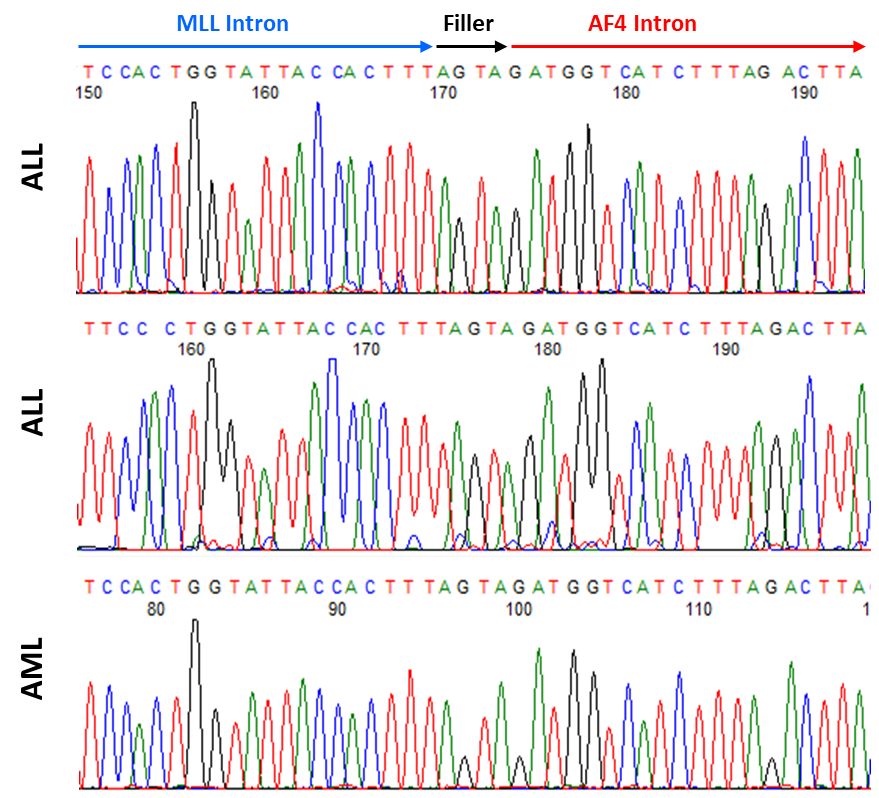

**ALL**

**ALL Relapse**

**AML Relapse**

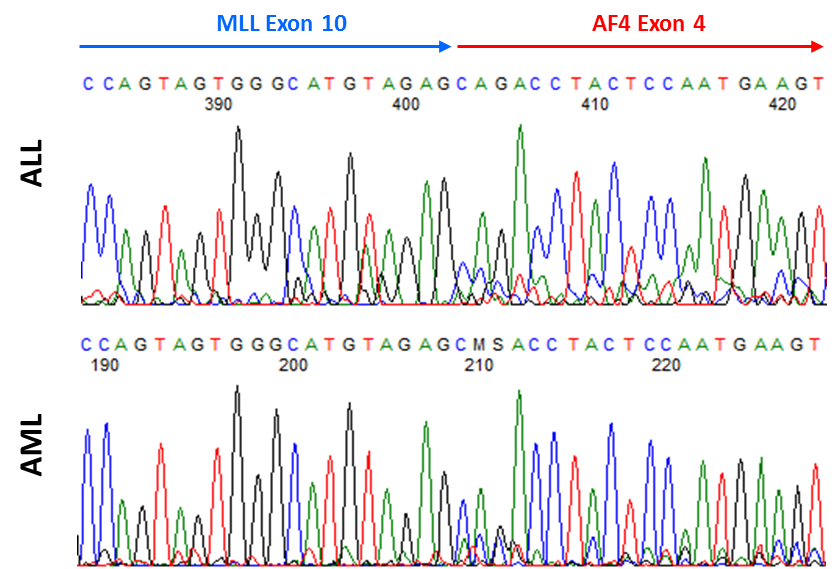

**Lymphoid fraction**

**Myeloid fraction**

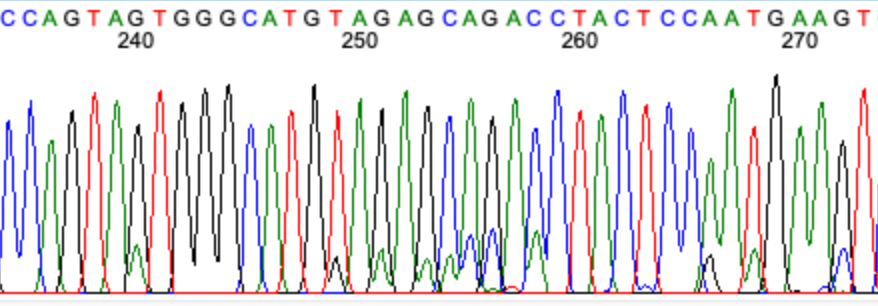

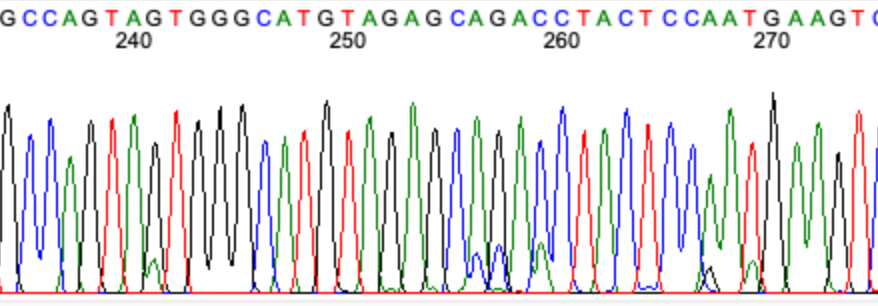

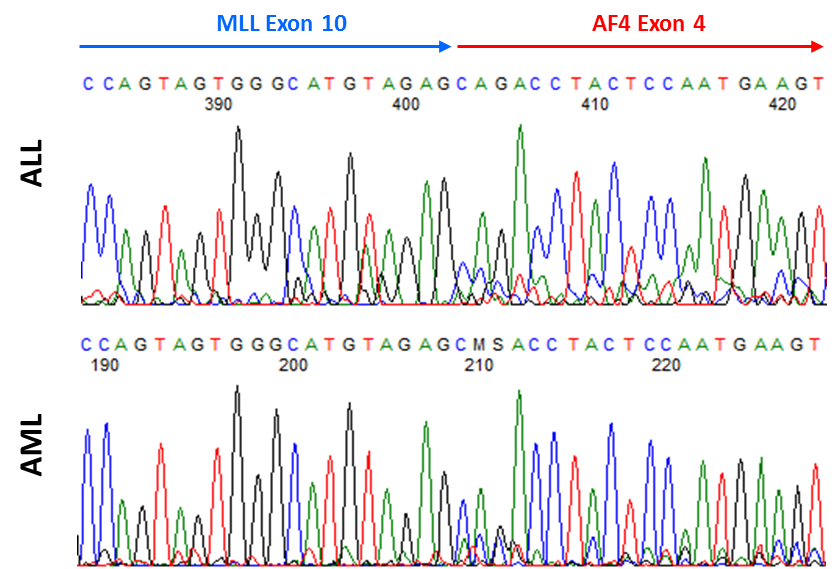

**Myeloid fraction**

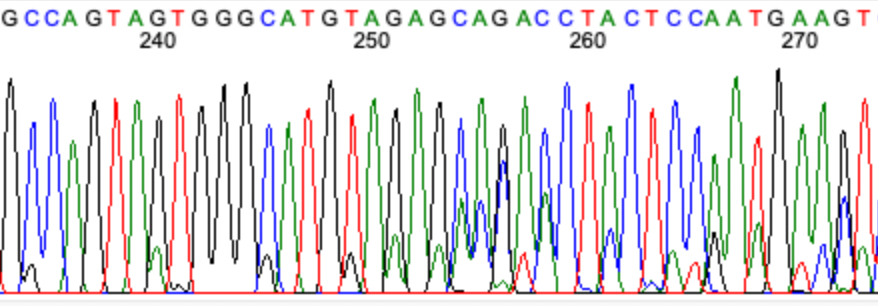

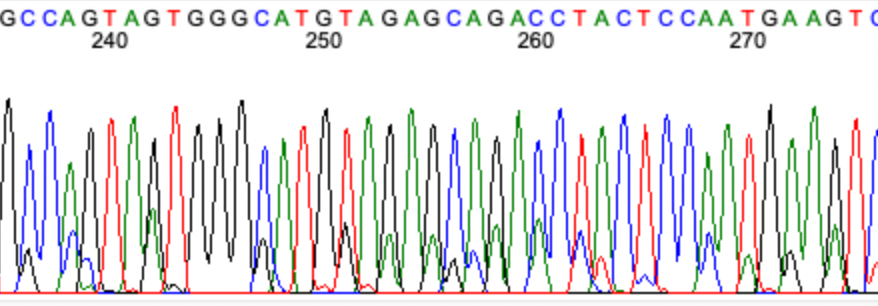

**Lymphoid fraction**

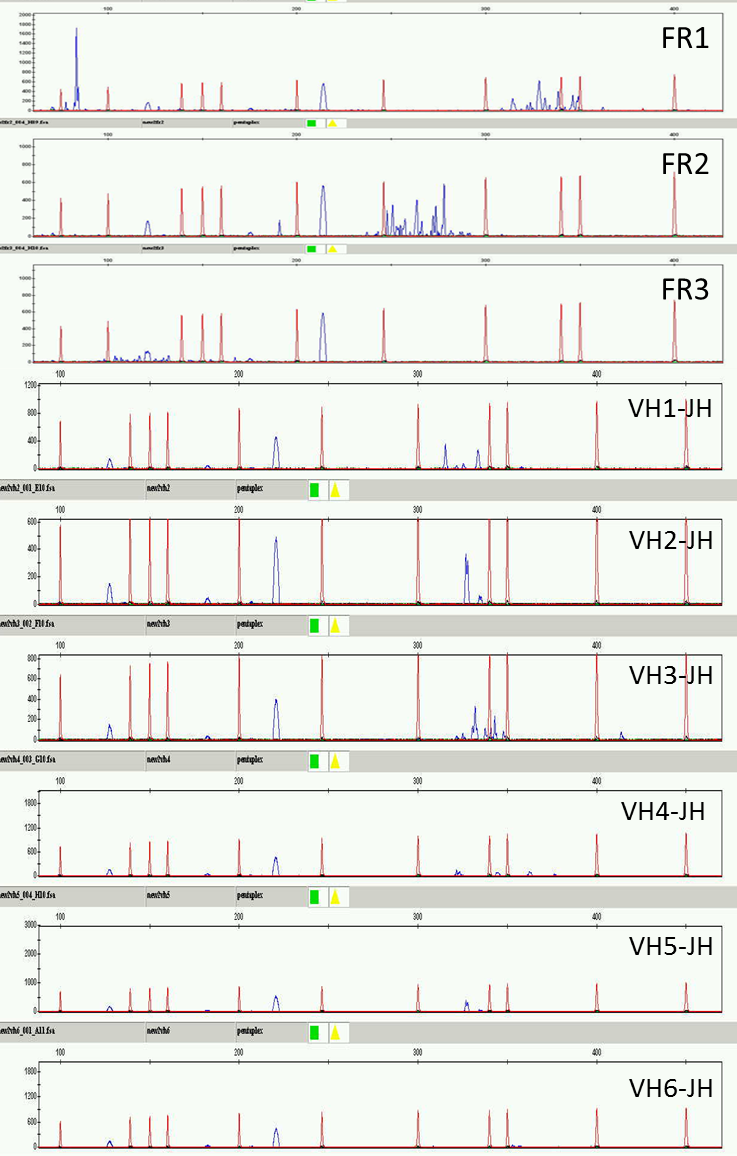

LS01AML

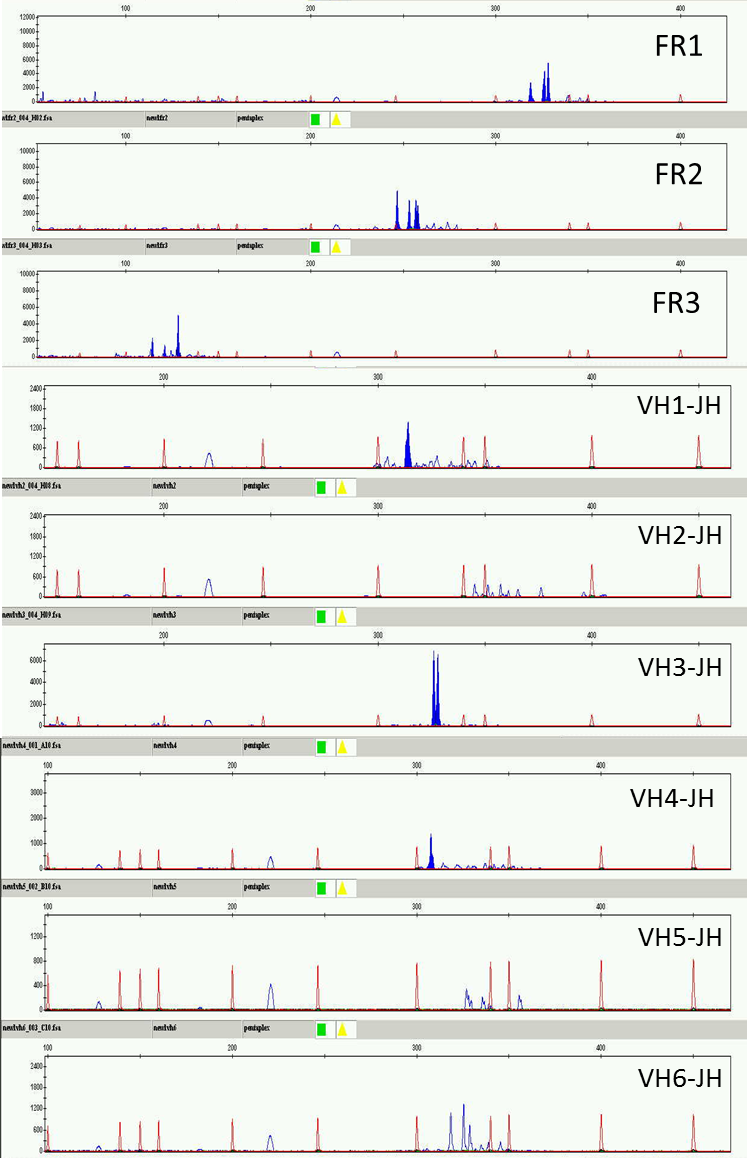

LS01ALL

Figure S2

A

C

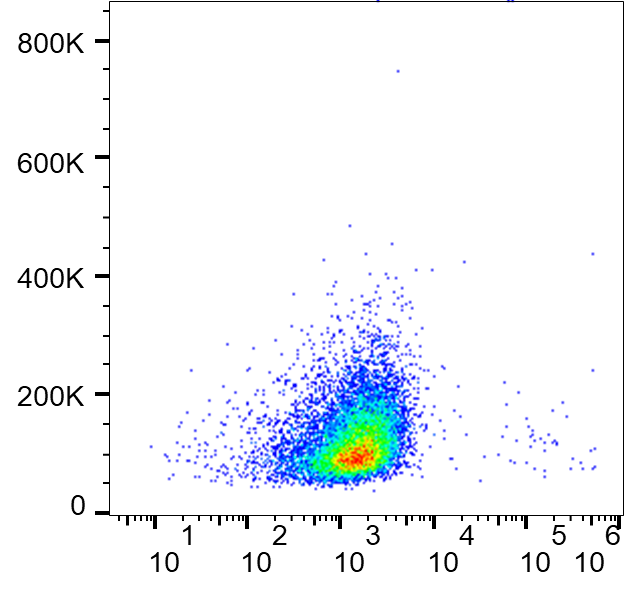

**CD34**

**SSC-A**

**CD19**

**CD33**

**LS01PALL**

**NSG**

**NSG**

**MISTRG**

**MISTRG**

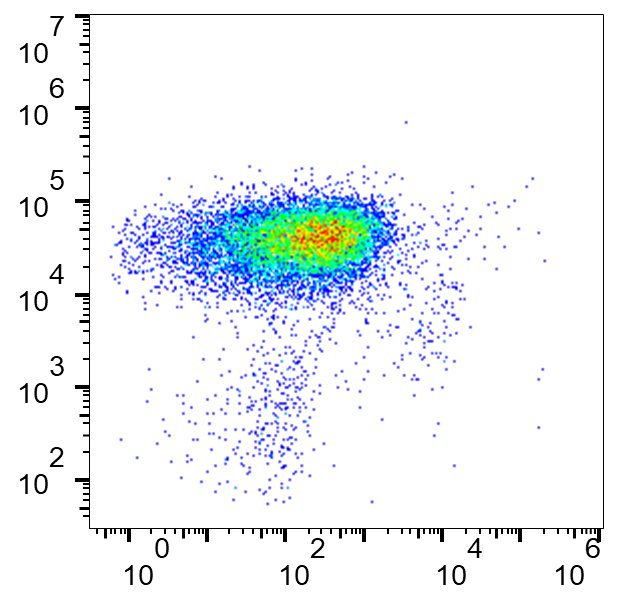

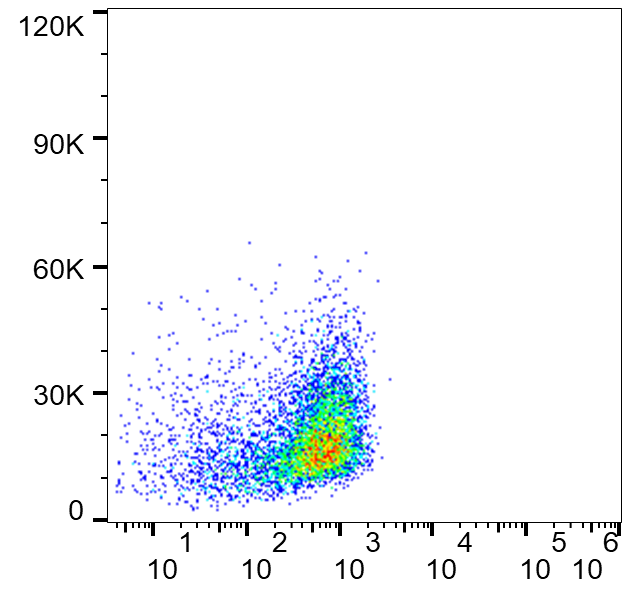

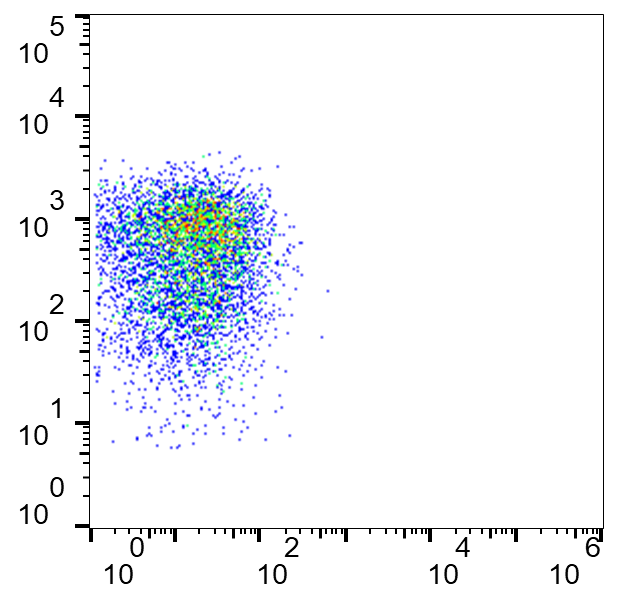

**LS01RAML**

**SSC-A**

**CD19**

**MISTRG**

**MISTRG**

**CD34**

**CD33**

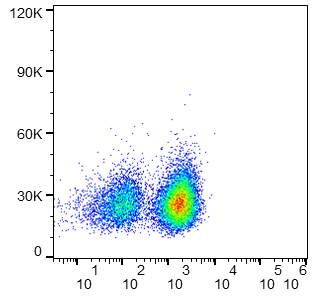

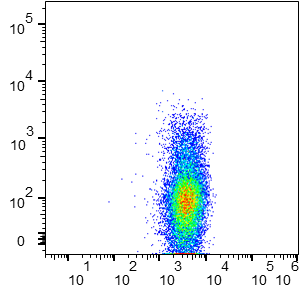

B

**NSG**

**LS01PALL**

$$100\% engraftment$$

**(6/6 mice)**

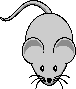

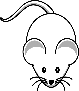

**NSG**

**(0/5 mice)**

$$50\% engraftment$$

**LS01RAML**

**MISTRG**

**(4/4 mice)**

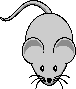

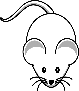

*Transplantation*

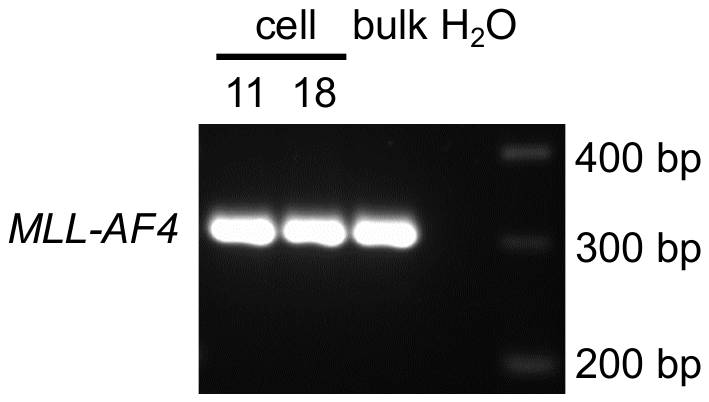

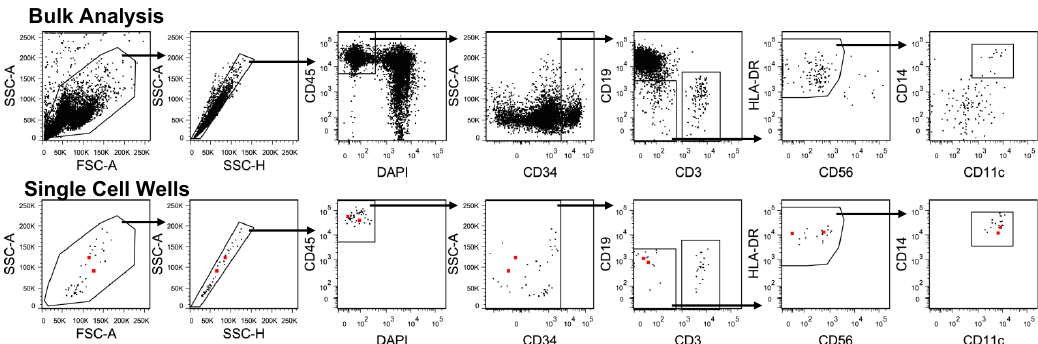

**Single Cell sort:**

**Lin(-)CD34(+)**

**Bulk sort:**

**HSCs vs MPPs**

E

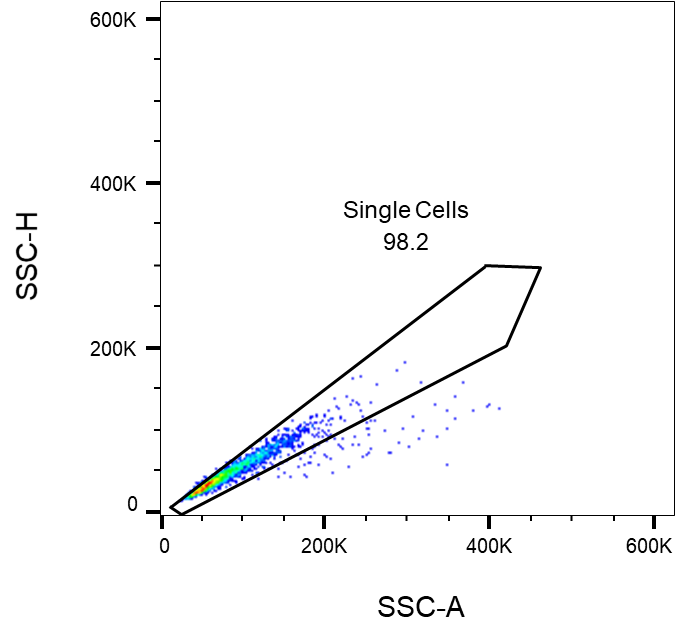

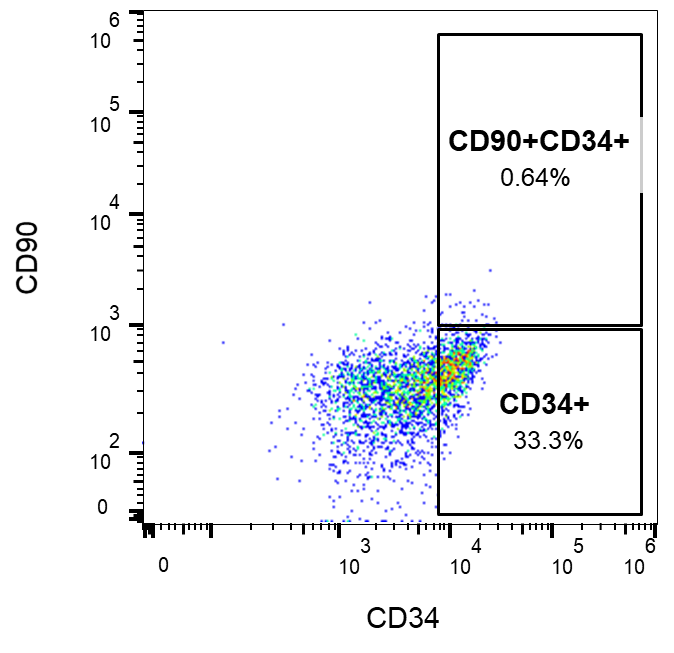

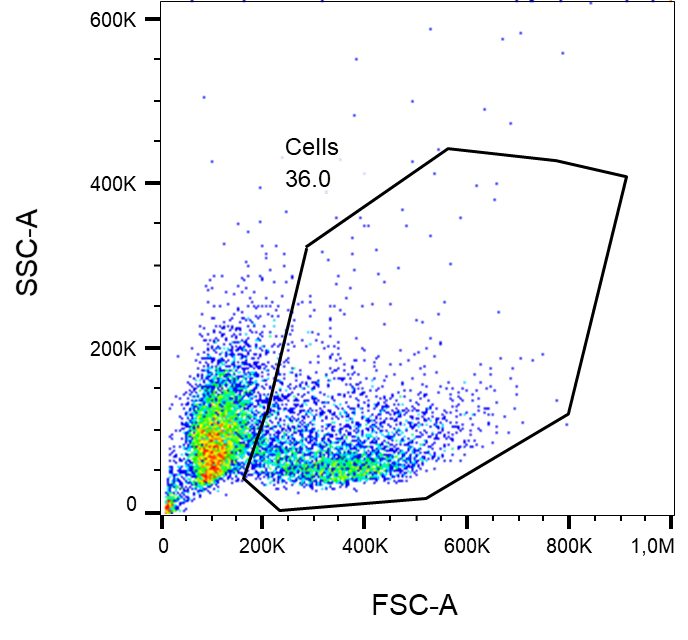

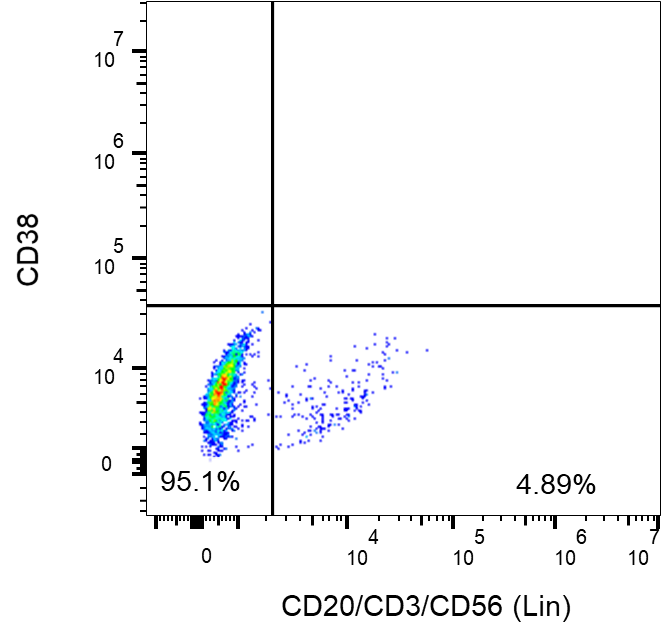

**HSCs**

**MPPs**

**MPAL2**

**MPAL1**

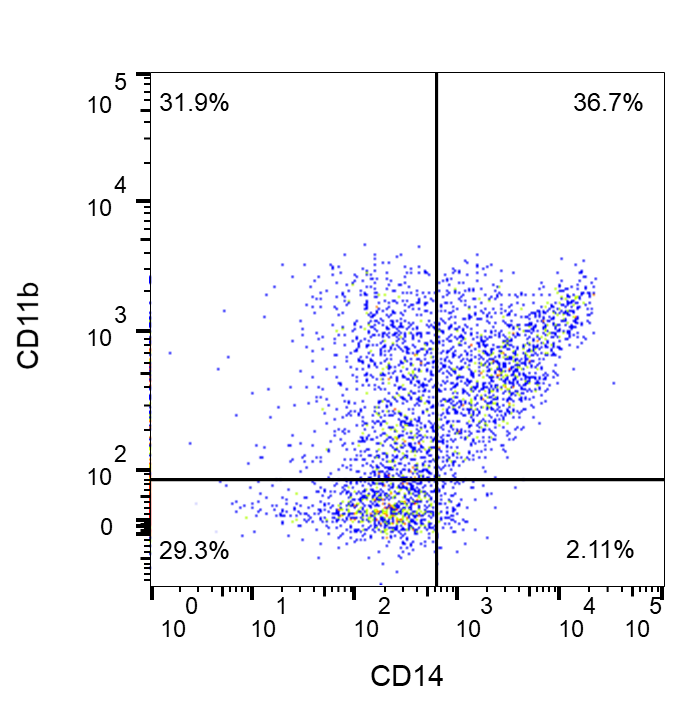

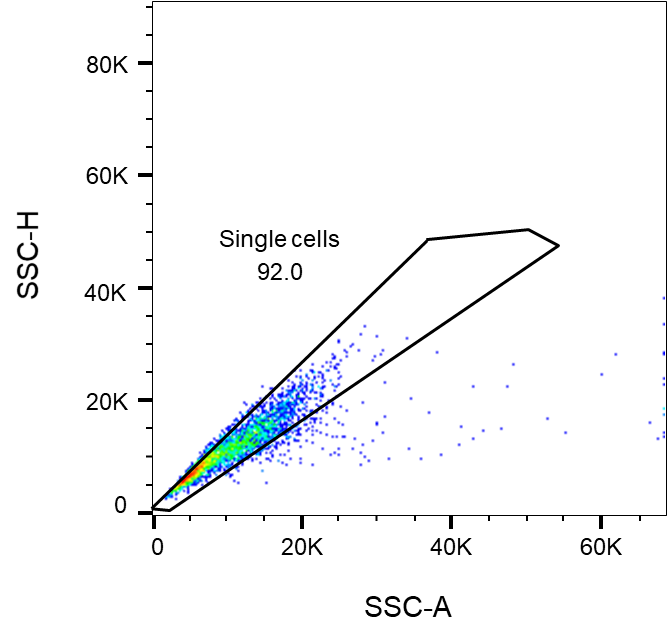

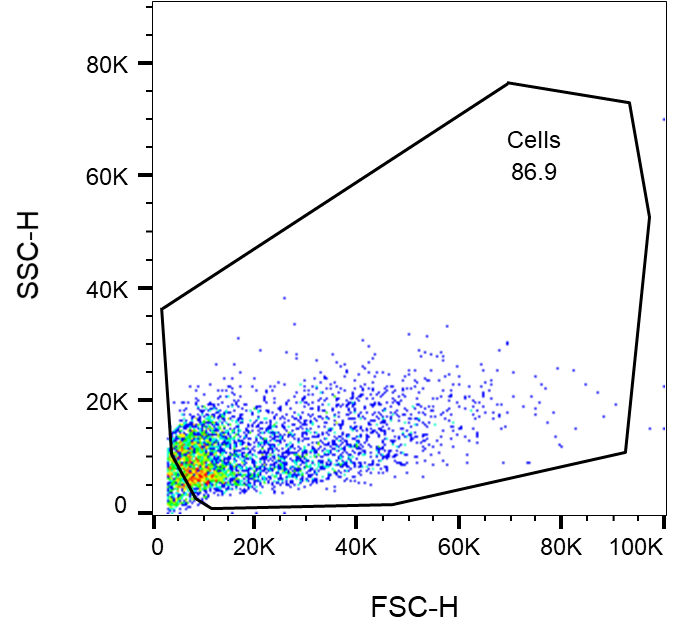

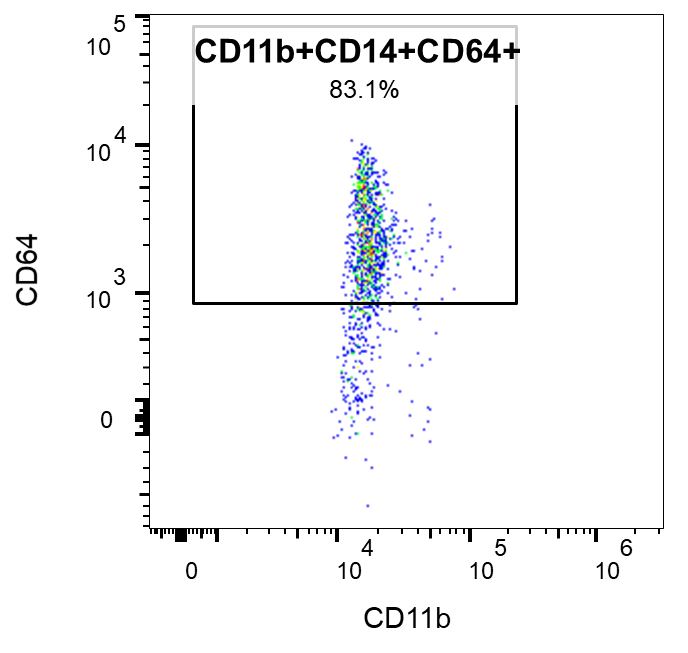

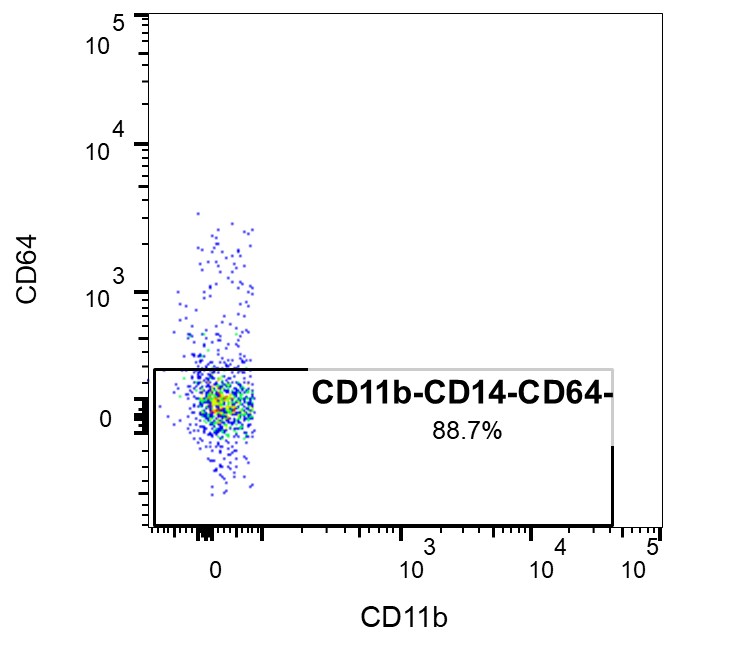

**Lymphoid**

**Myeloid**

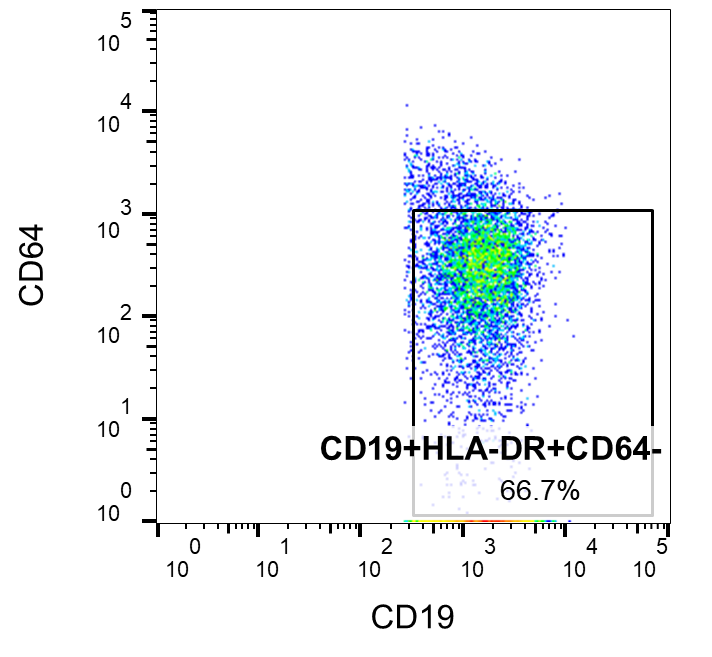

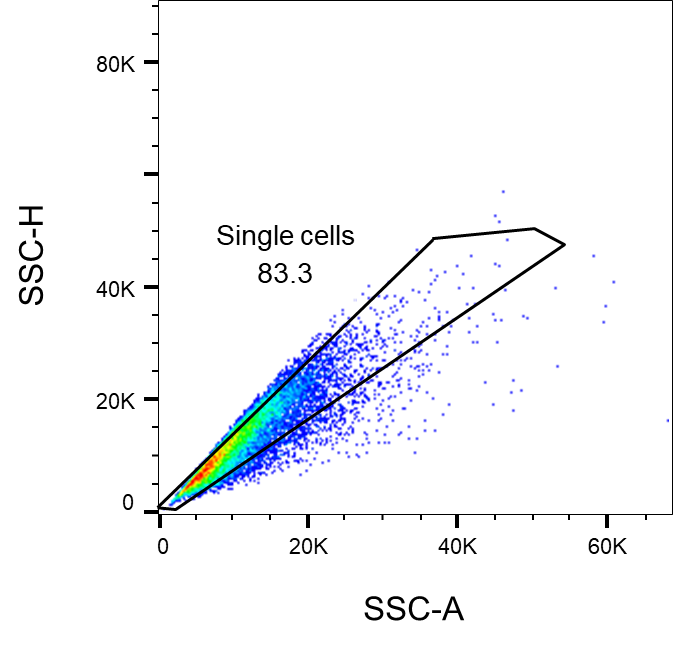

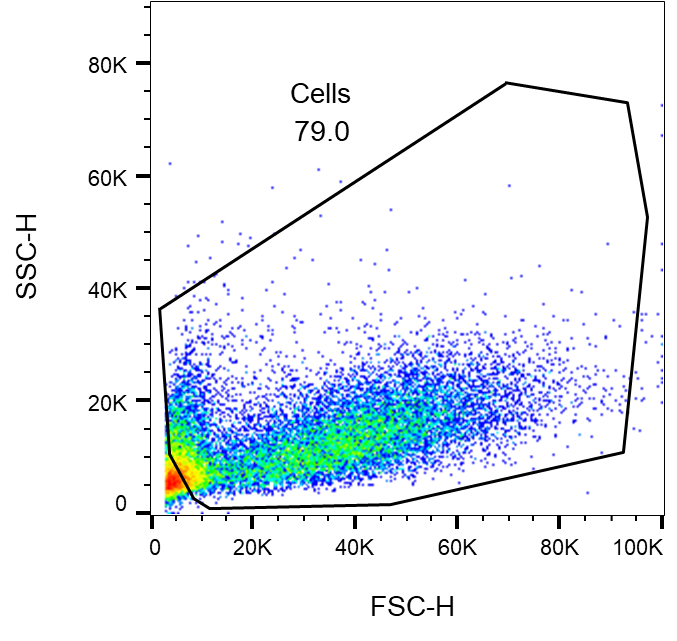

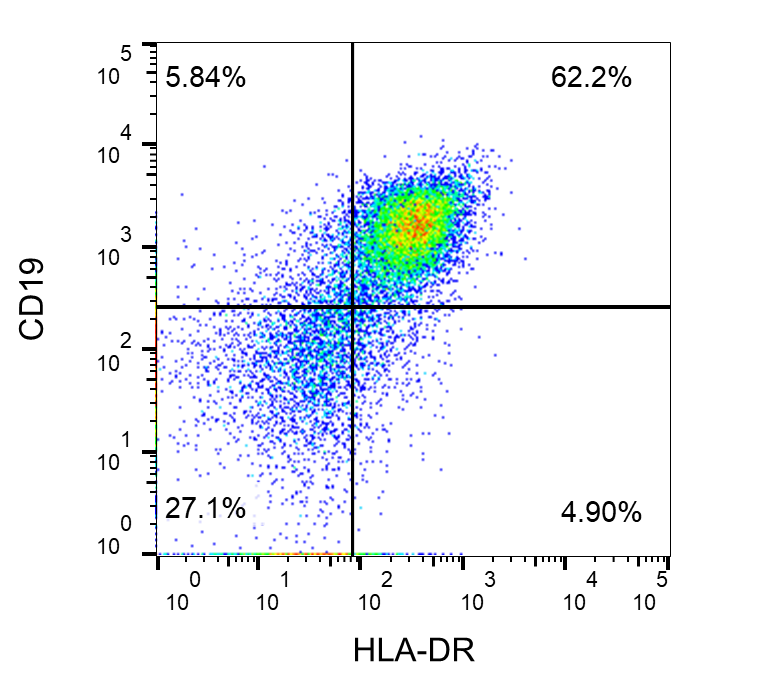

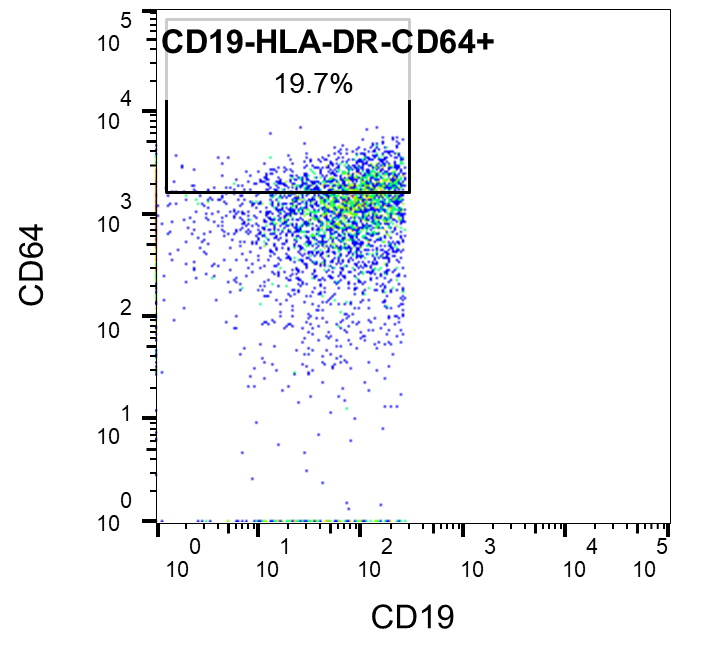

**Lymphoid**

**Myeloid**

A

B

Figure S3

C

D

Single cell deposition

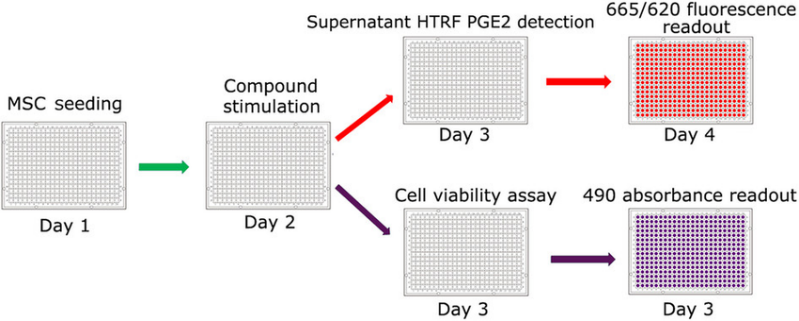

Single cell expansion

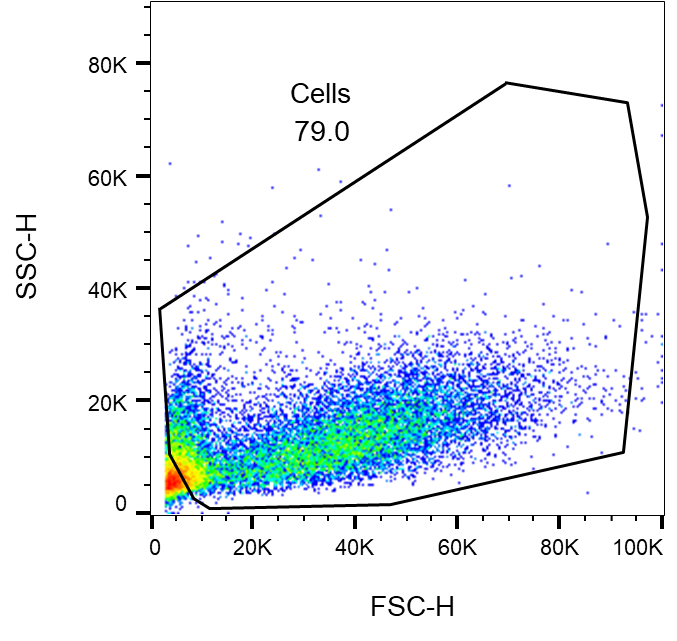

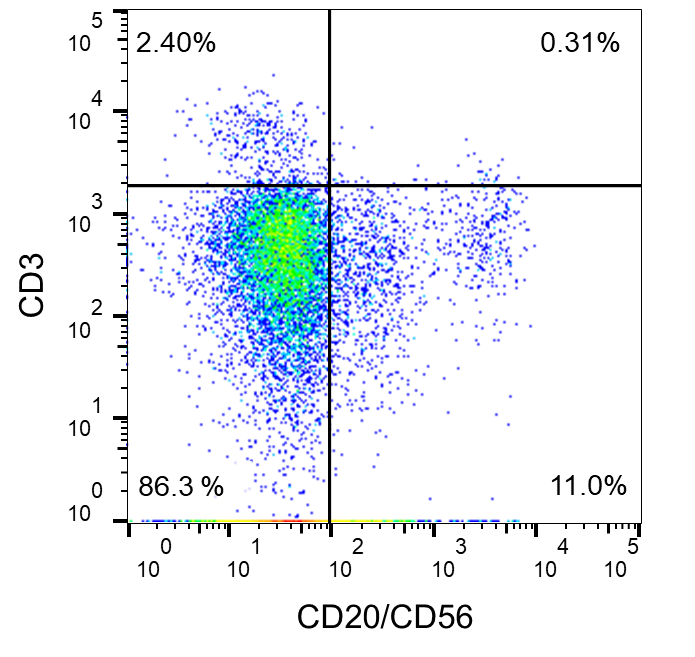

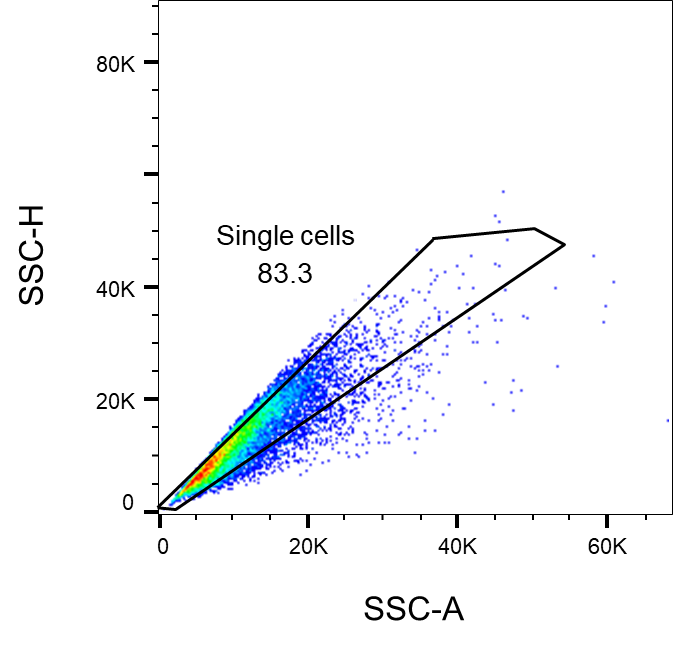

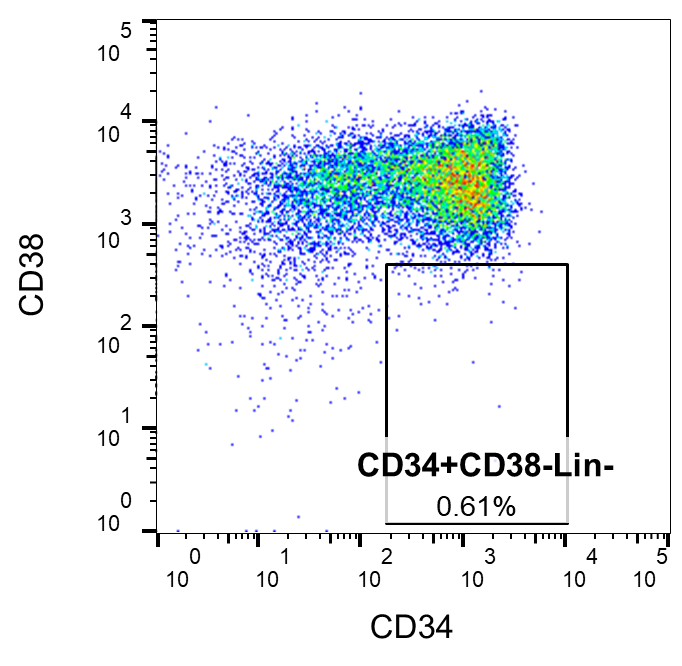

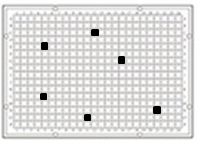

**HSCs+MPPs**

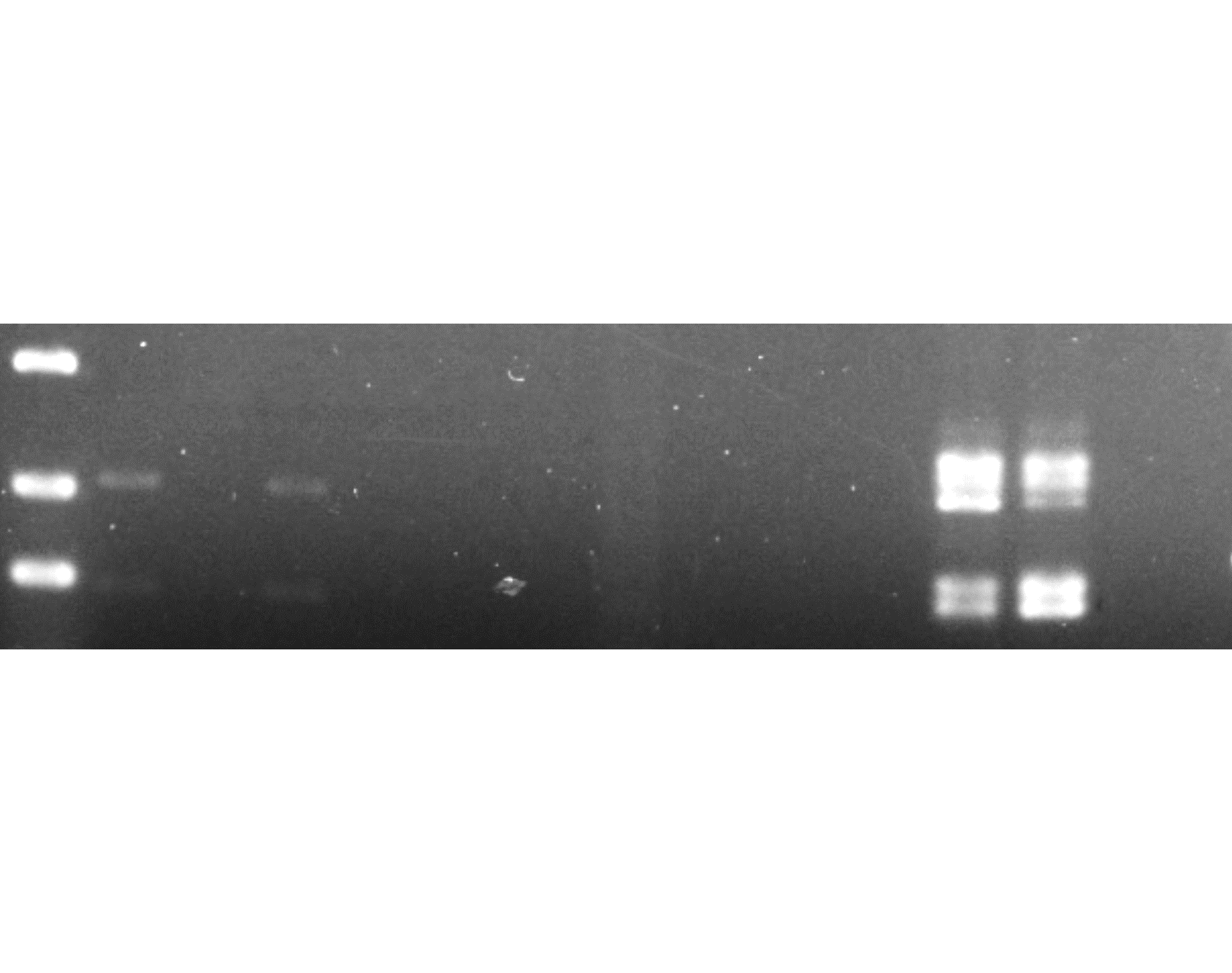

**MPAL1**

**Lymphoid**

**Myeloid**

H_2_O ctr

700 bp

500 bp

400 bp

**single-cell derived clones (Lin-CD34+)**

1	2	3	4	5	6	7	8	9	10

B

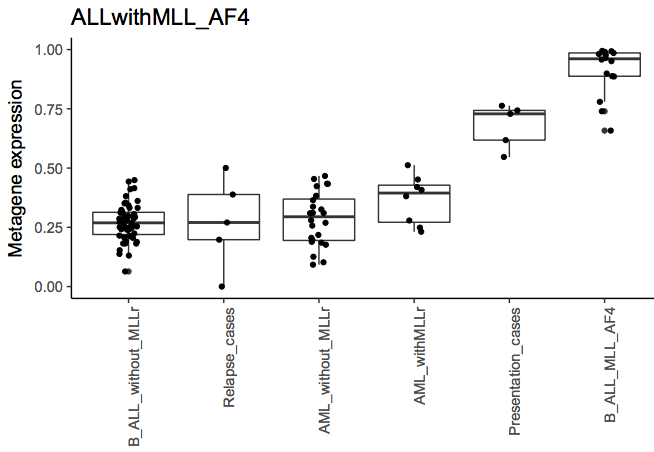

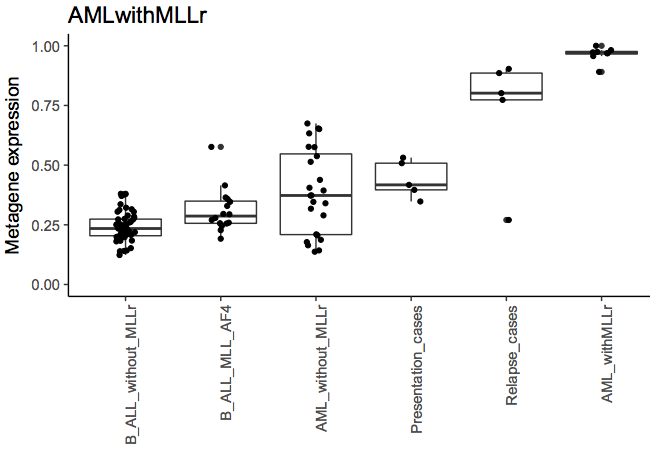

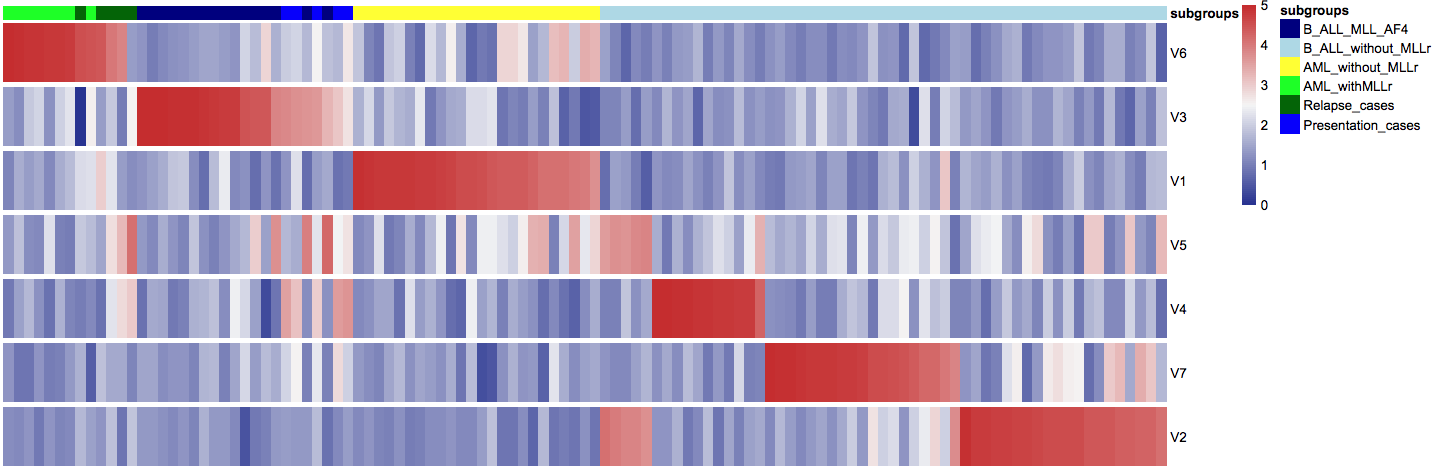

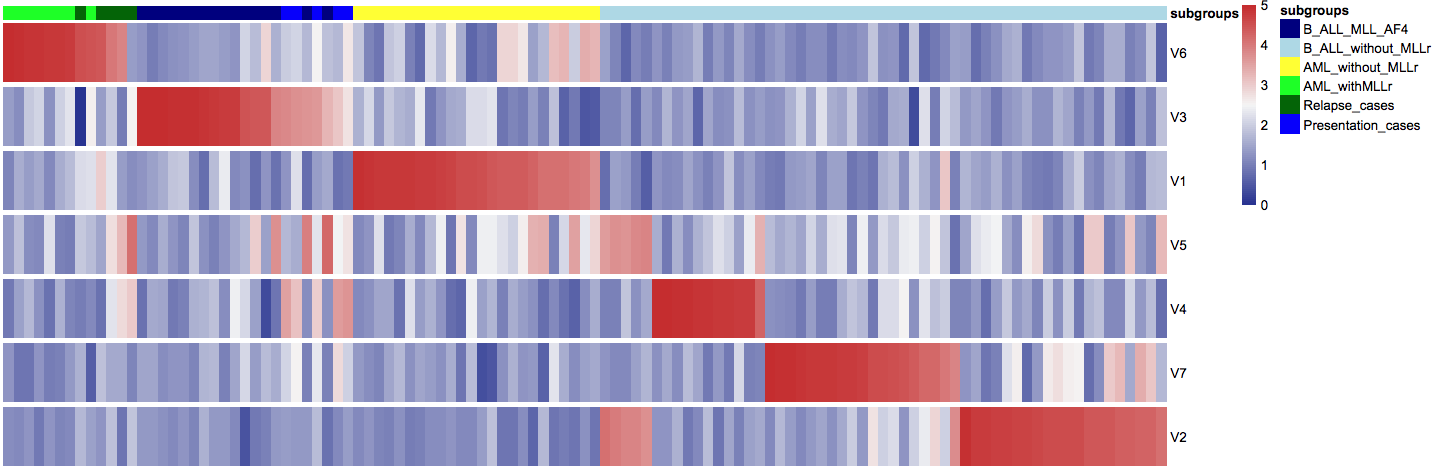

**subgroups**

**V6	V3	V1	V5	V4	V7	V2**

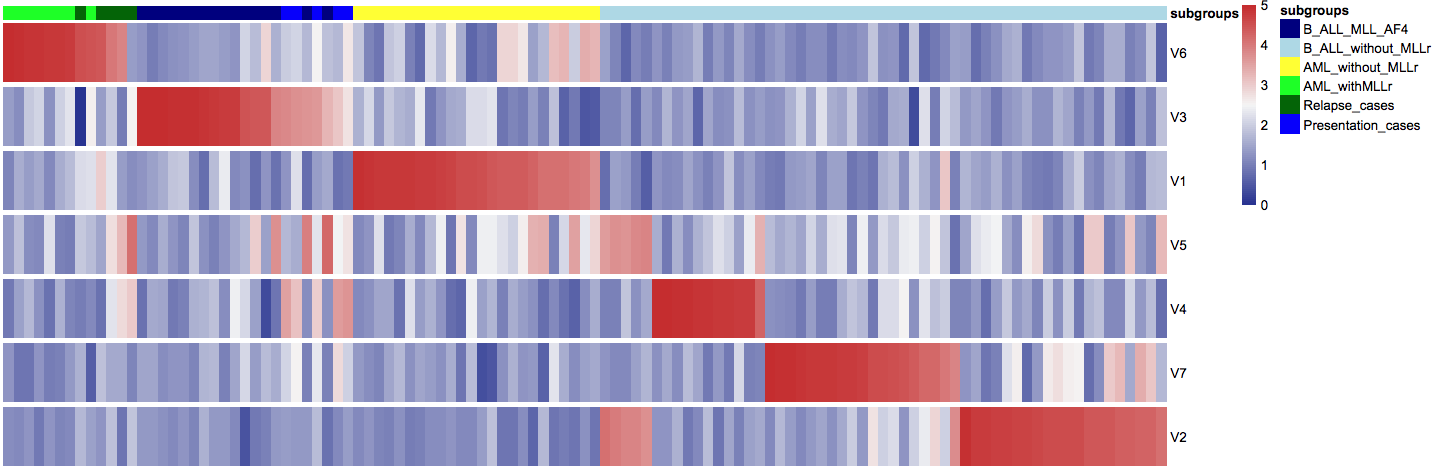

B_ALL_without MLLr

Relapse_cases

Presentation_cases

B_ALL_MLL_AF4

AML_withMLLr

AML_without_MLLr

B_ALL_without MLLr

Relapse_cases

AML_without_MLLr

AML_withMLLr

Presentation_cases

B_ALL_MLL_AF4

A

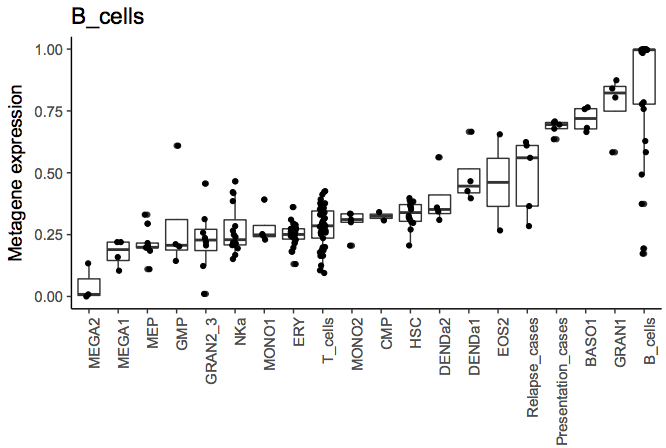

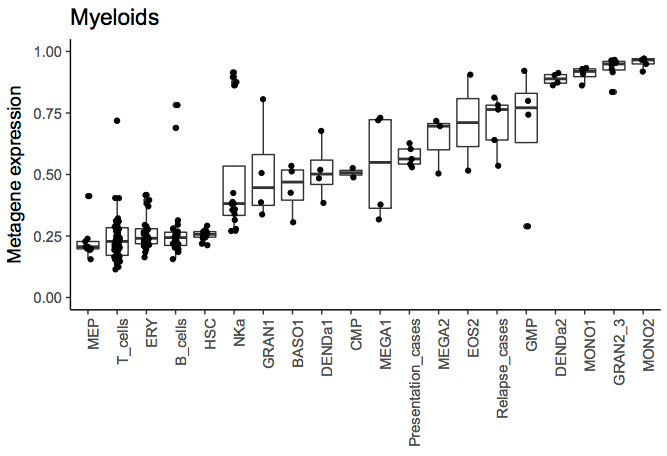

Figure S4

C

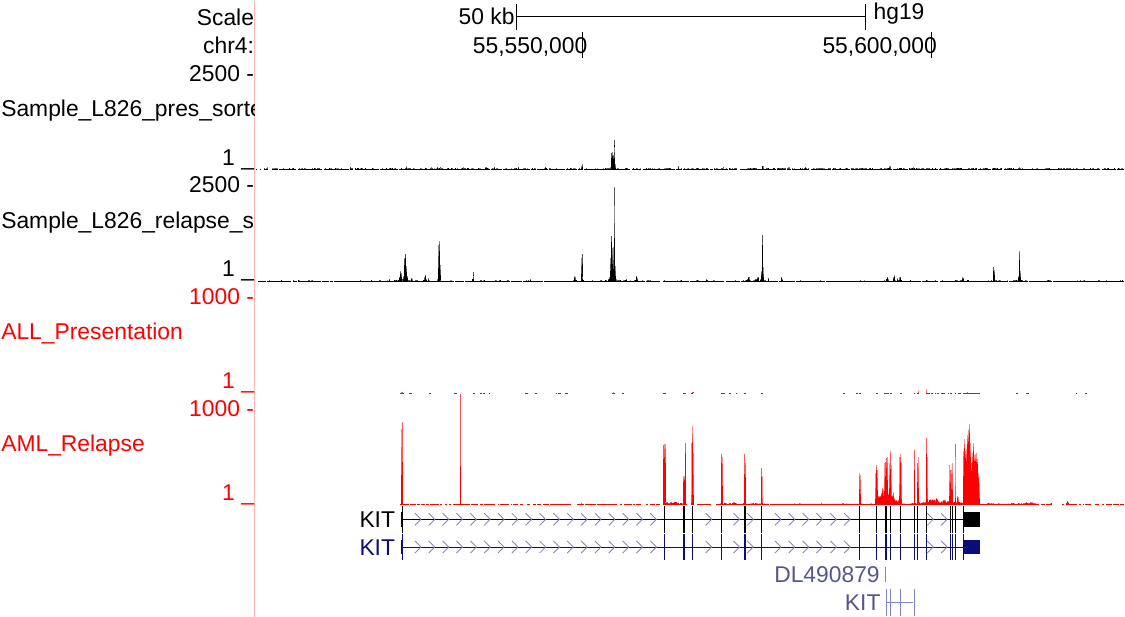

*KIT*

TSS

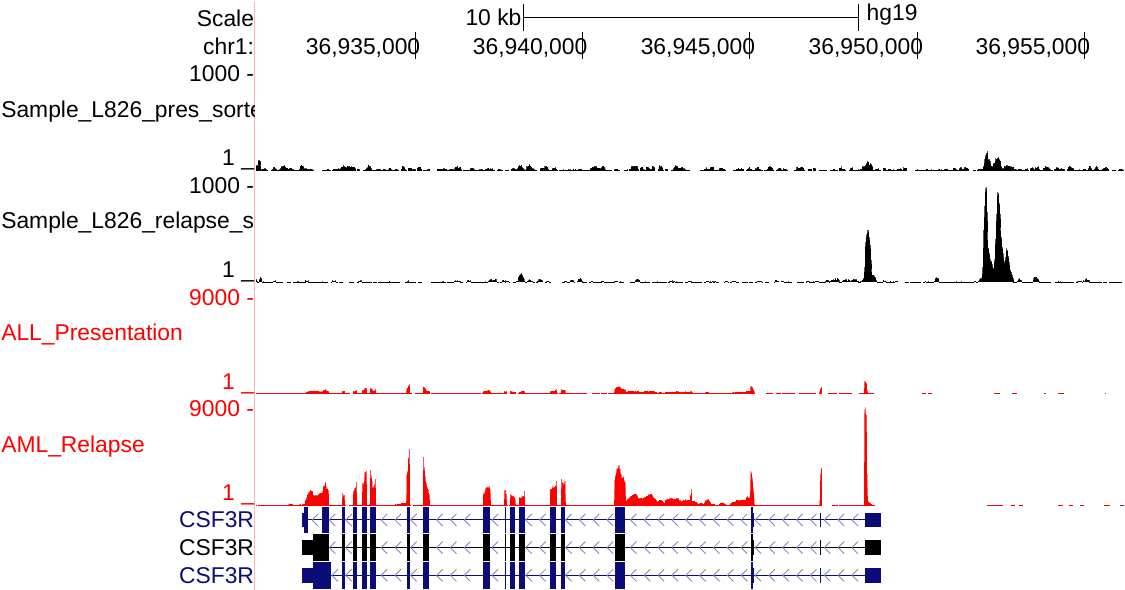

*CSF3R*

RNA-seq AML

DHS-seq ALL

TSS

DHS-seq AML

RNA-seq ALL

RNA-seq AML

DHS-seq ALL

DHS-seq AML

RNA-seq ALL

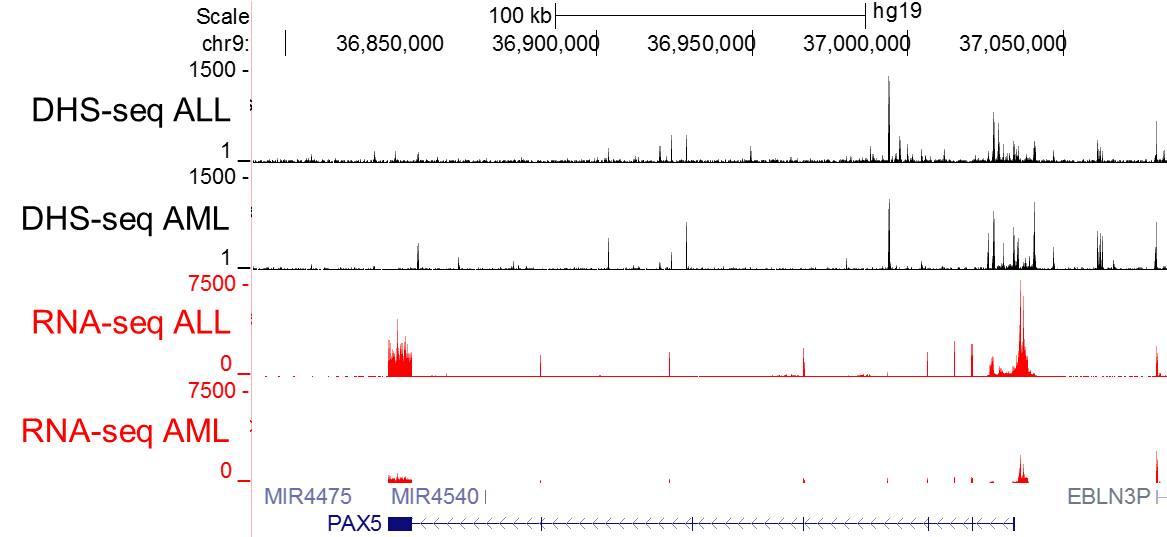

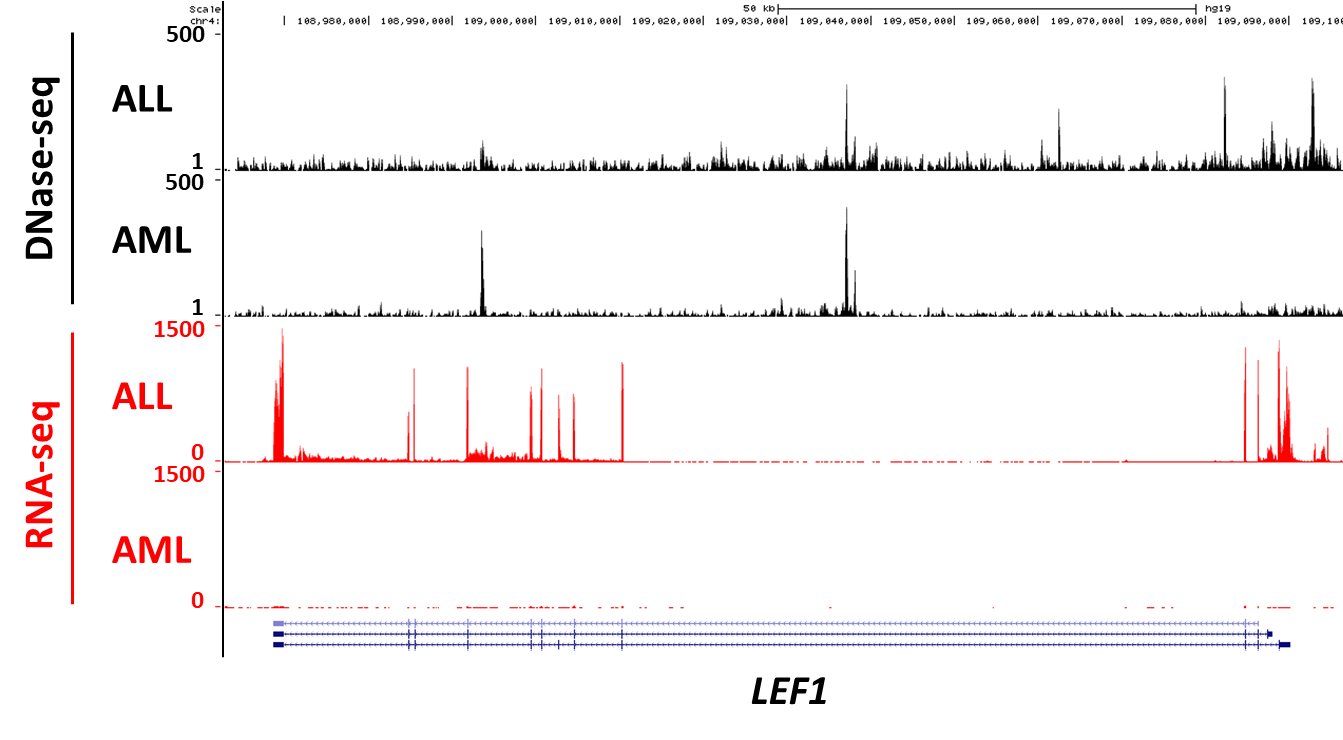

LEF1

TSS

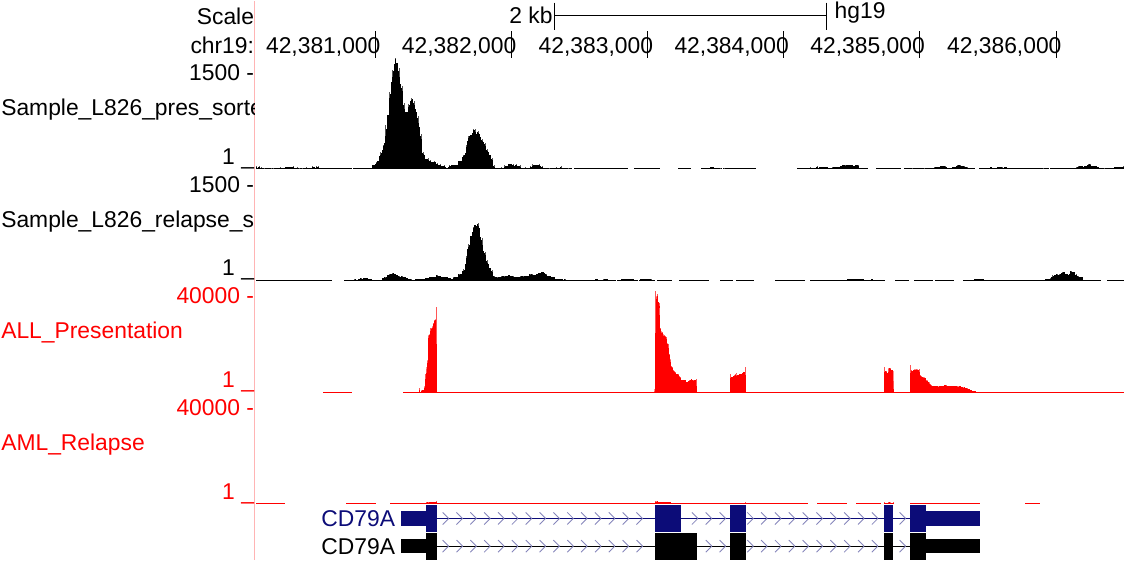

*CD79A*

TSS

TSS

*LEF1*

*PAX5*

RNA-seq AML

DHS-seq ALL

DHS-seq AML

RNA-seq ALL

RNA-seq AML

DHS-seq ALL

DHS-seq AML

RNA-seq ALL

D

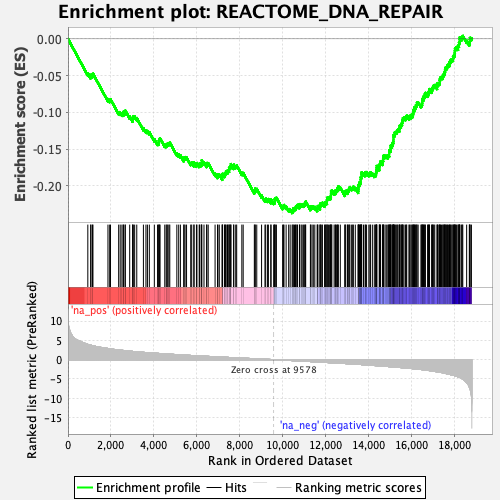

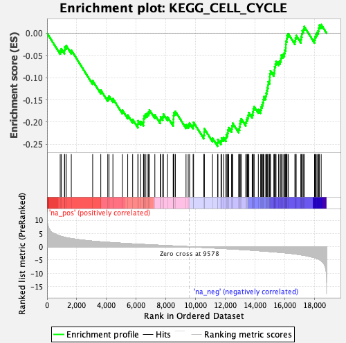

relapse	diagnostic

NES=-4.95

FDR<10^-4^

relapse	diagnostic

NES=-3.36

FDR<10^-4^

TSS

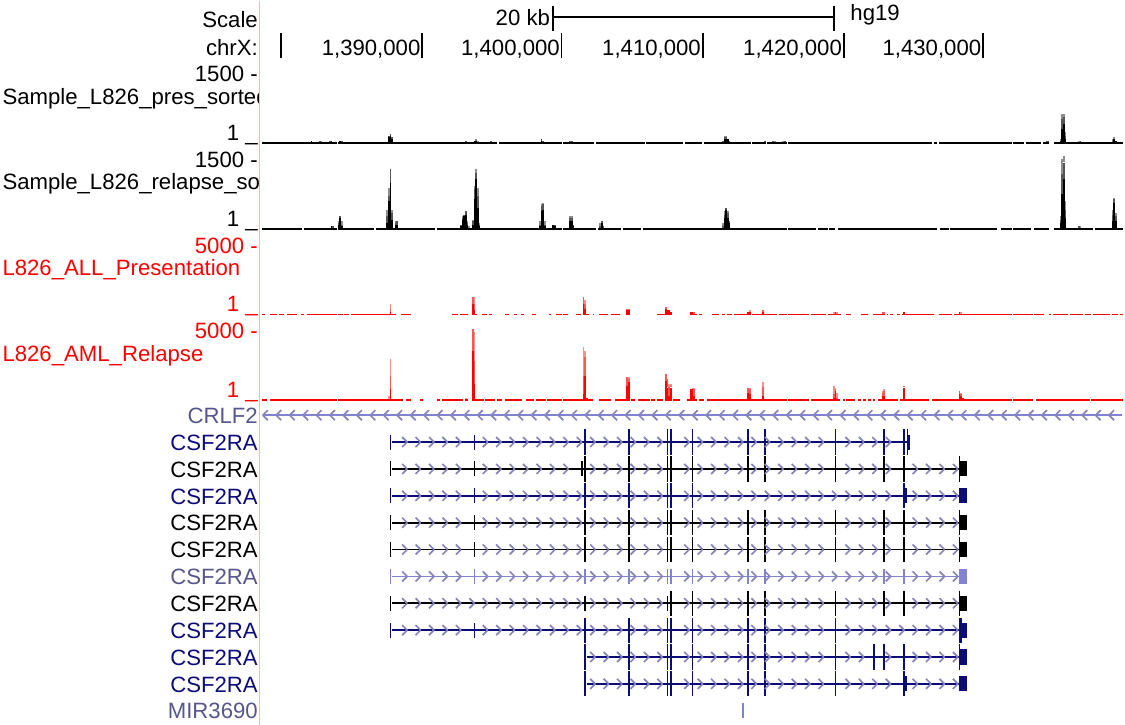

TSS

*CSF2RA*

RNA-seq AML

DHS-seq ALL

DHS-seq AML

RNA-seq ALL

E

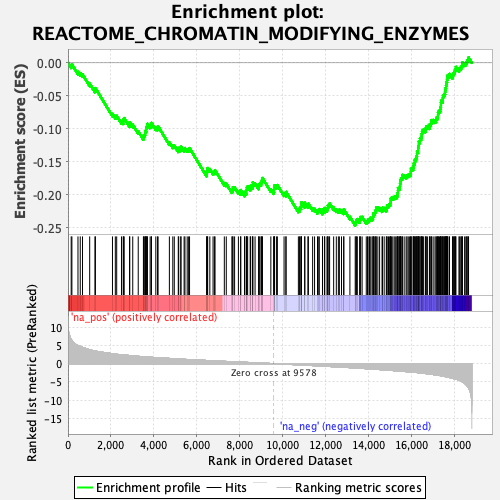

relapse	diagnostic

NES=-4.57

FDR<10^-4^

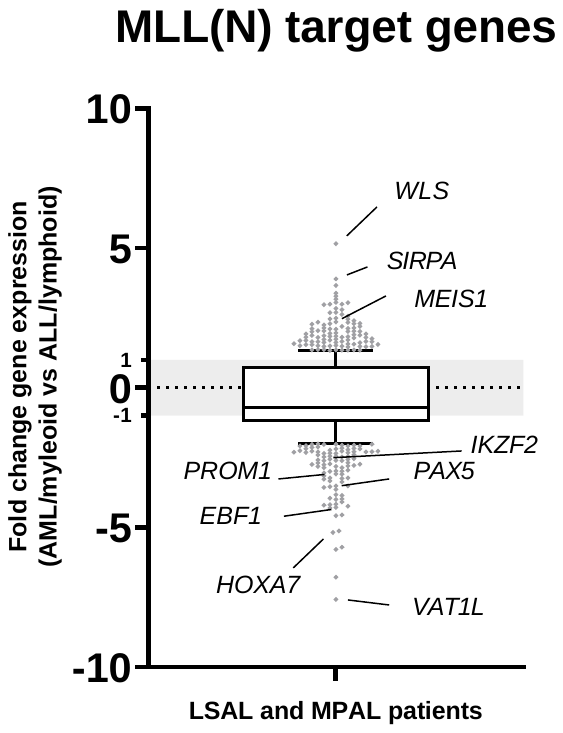

**MLL-AF4 target genes**

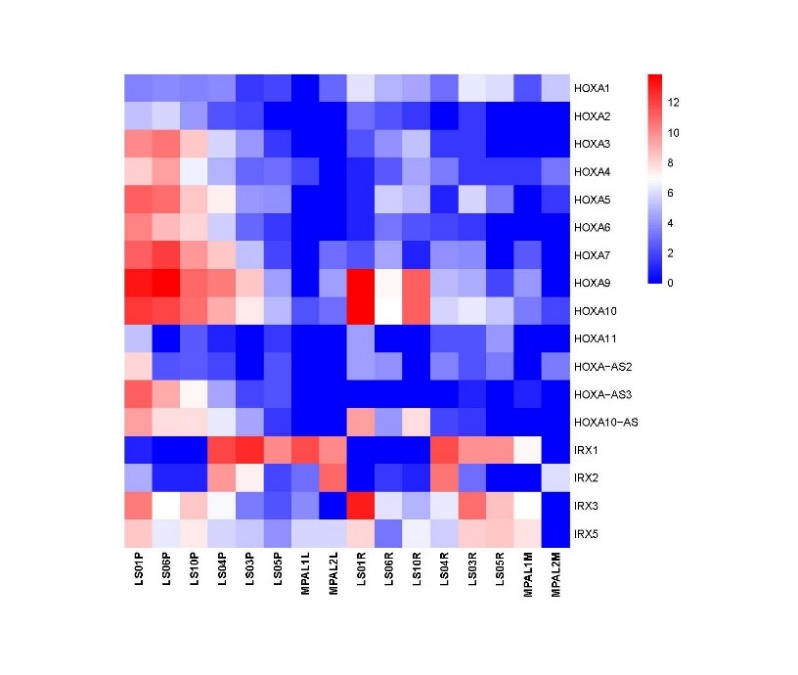

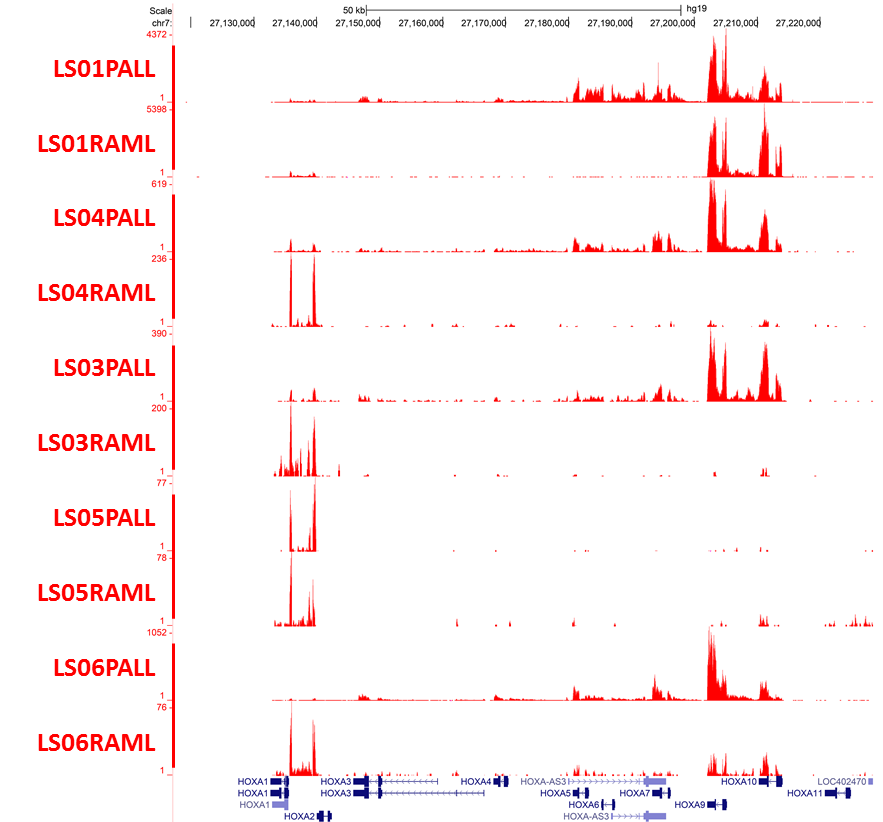

*HOXA cluster*

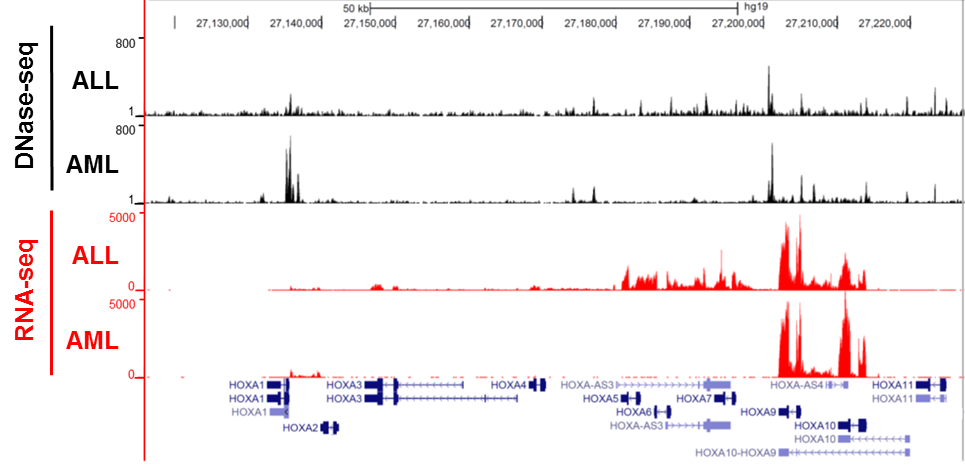

A

B

*HOXA cluster*

C

**ALL**

**AML**

**Log2 values**

D

**423**

**MLL(N) ChIPseq
by Kerry J.et at., 2017**

**RNAseq**

**LSALs and MPALs data**

**15670**

2620

**(total 4039)**

**(total 23503)**

996

4217

**Differentially expressed genes in AML/myeloid fraction**

**(total 5208)**

Figure S5

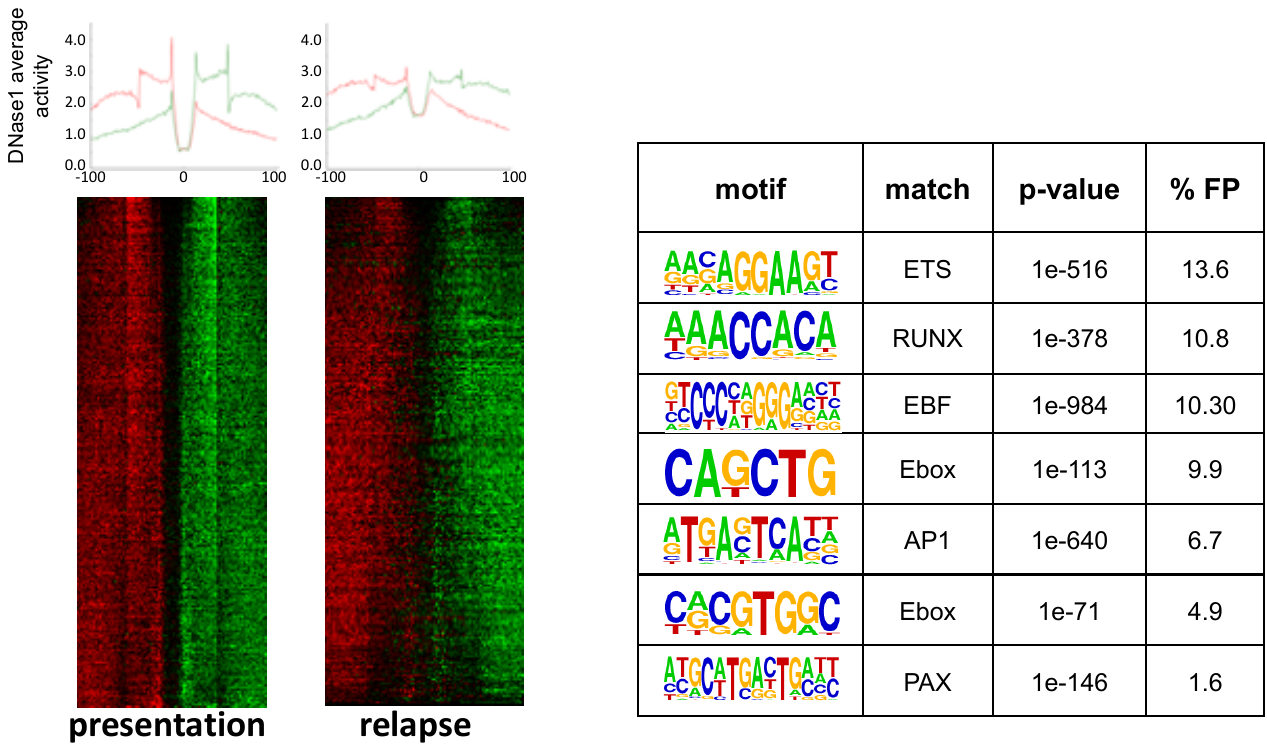

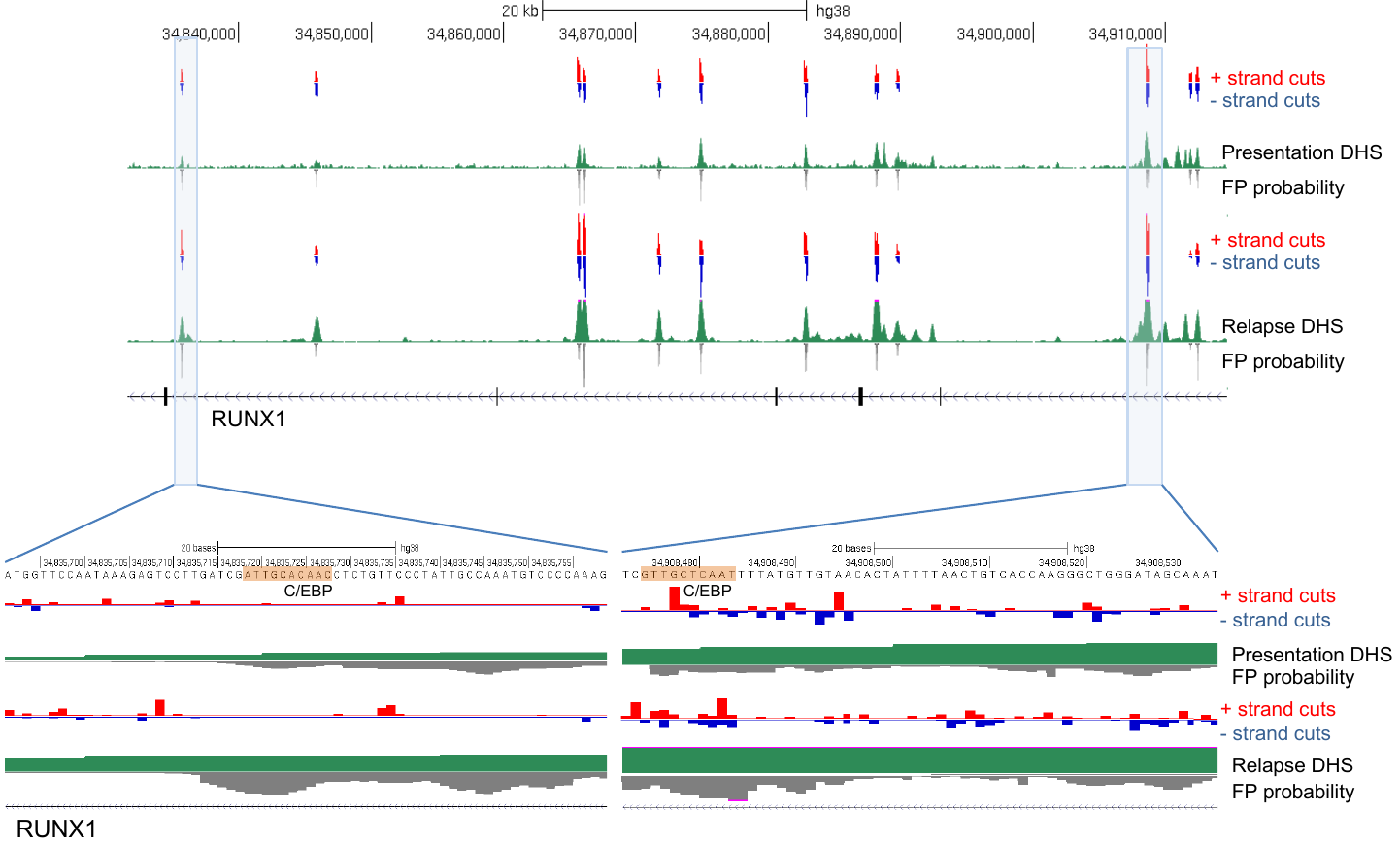

Figure S6

B

A

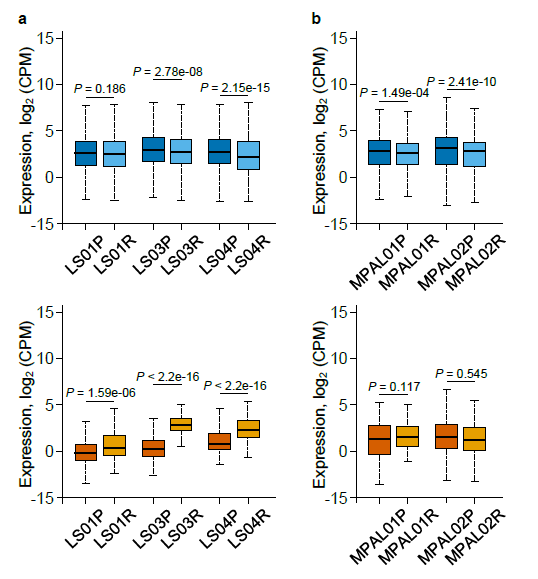

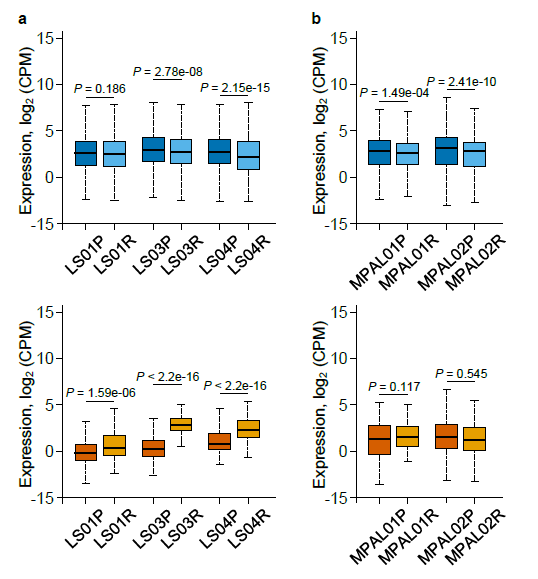

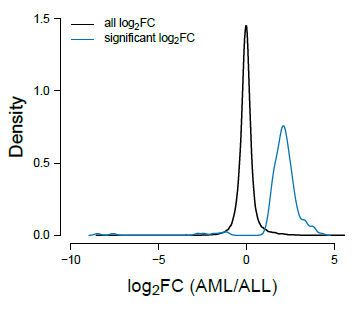

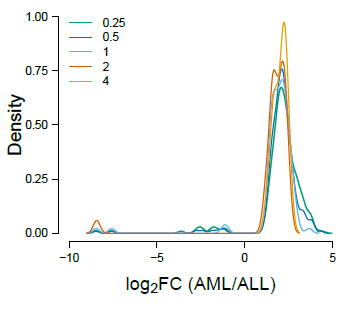

**MPALs**

**LSALs**

**6**

**97**

**160**

*PTK2B*

*EWSR1*

*MPRIP*

*GNAQ*

*ARFGAP2*

*ZEB2*

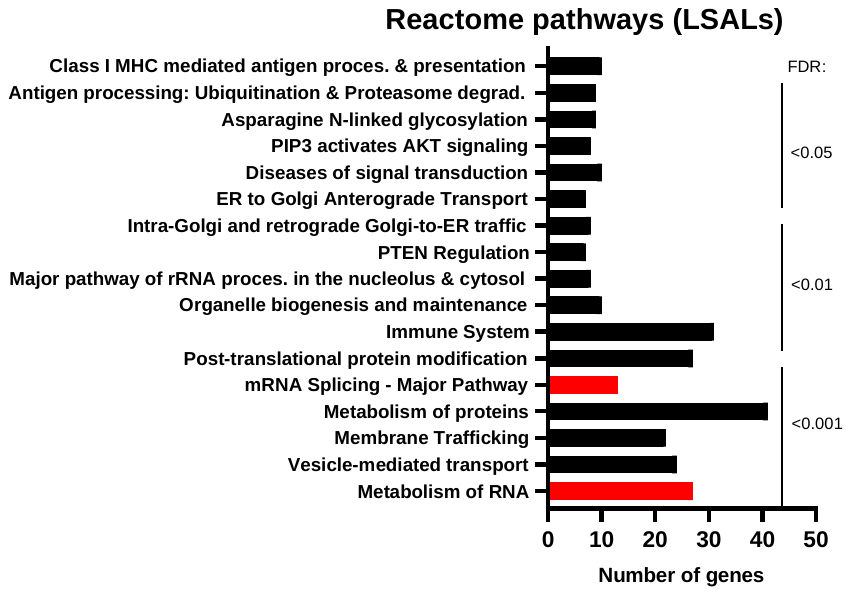

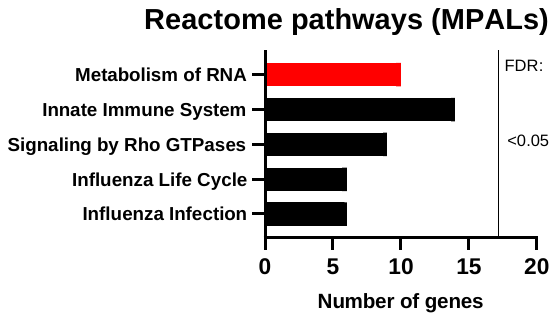

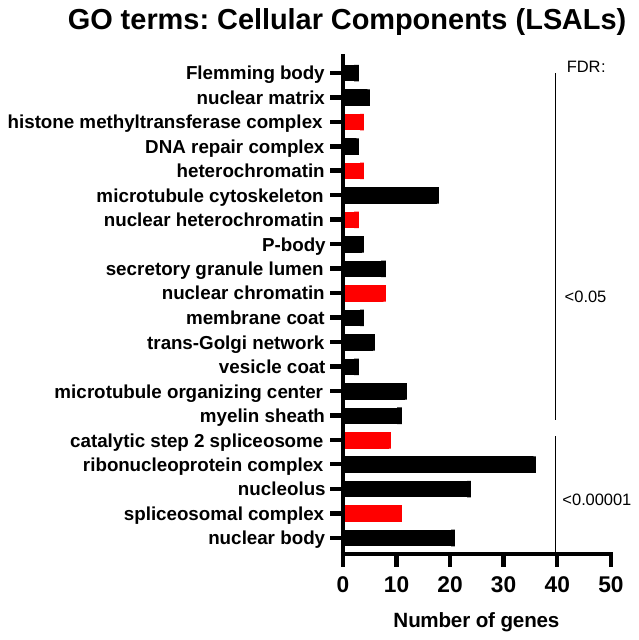

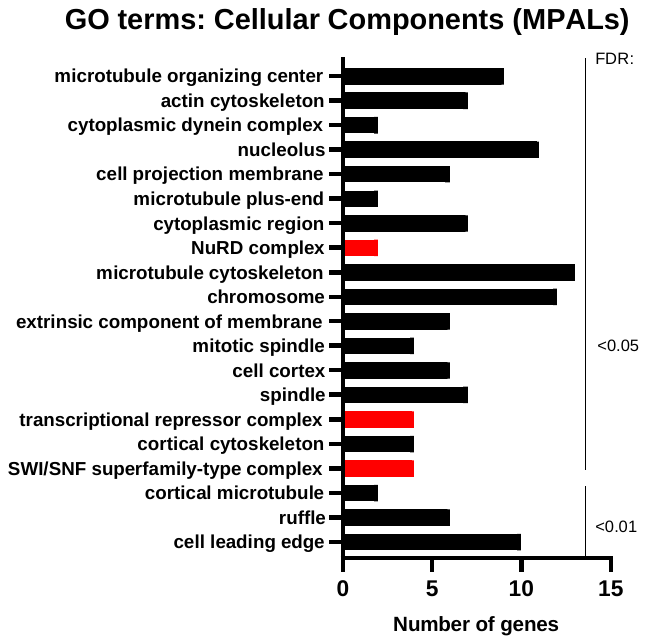

E

C

B

A

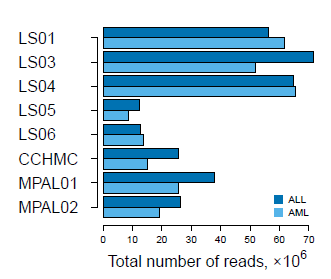

LS01

LS03

LS04

L505

LS06

LS10

MPAL1

MPAL2

Figure S7

**MPALs**

**LSALs**

**653**

**ALL**

**AML**

**343**

**1634**

**(2287)**

**(1977)**

**19**

**6**

**18**

**lymphoid**

**myeloid**

**(25)**

**(37)**

D

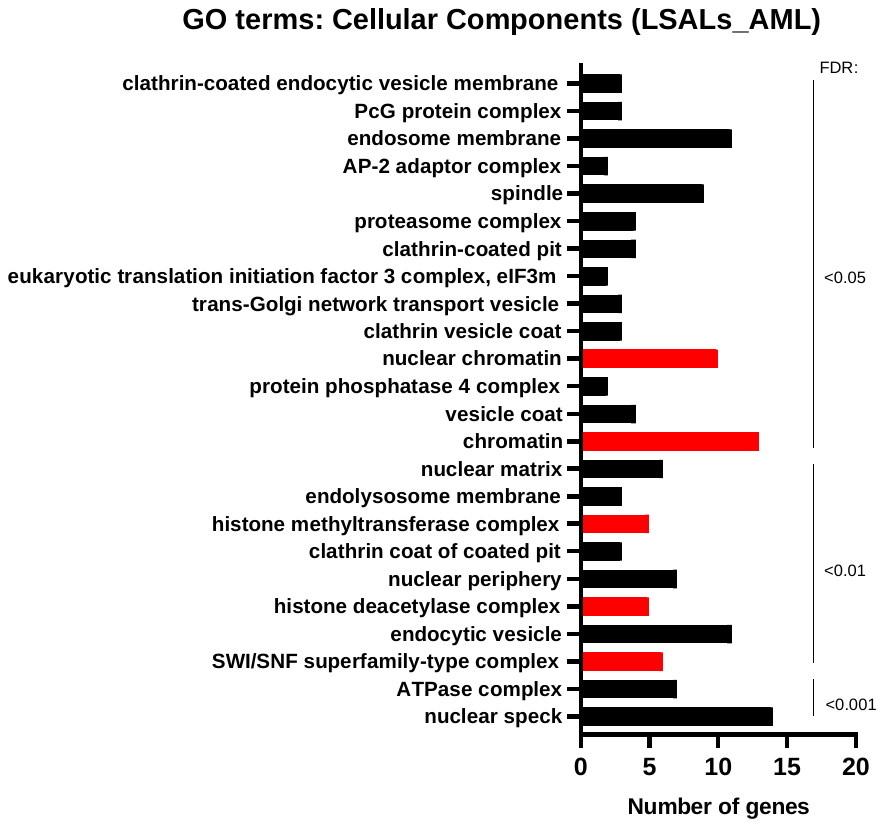

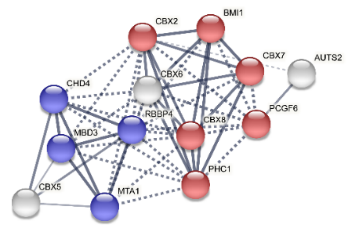

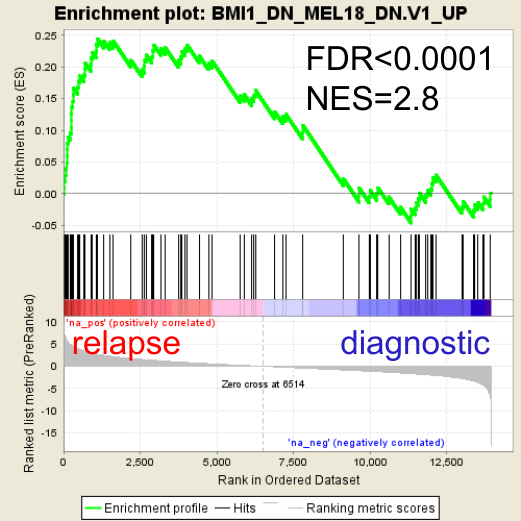

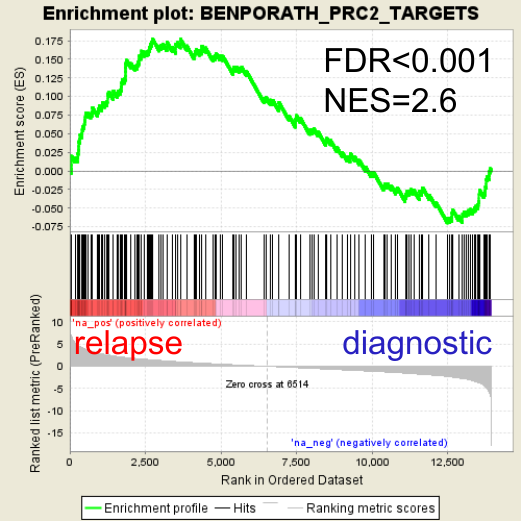

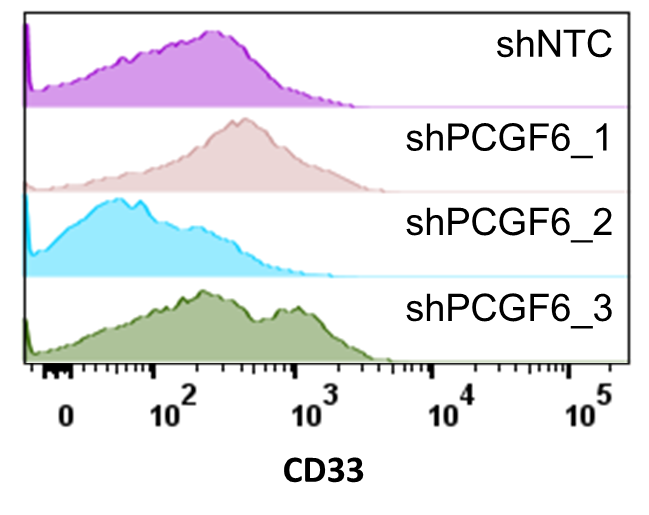

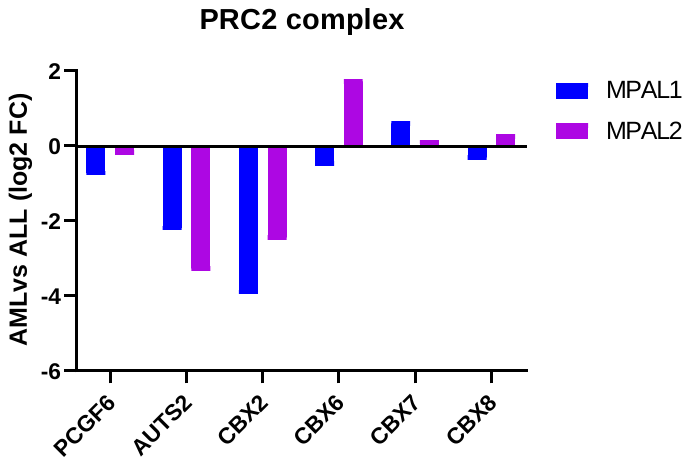

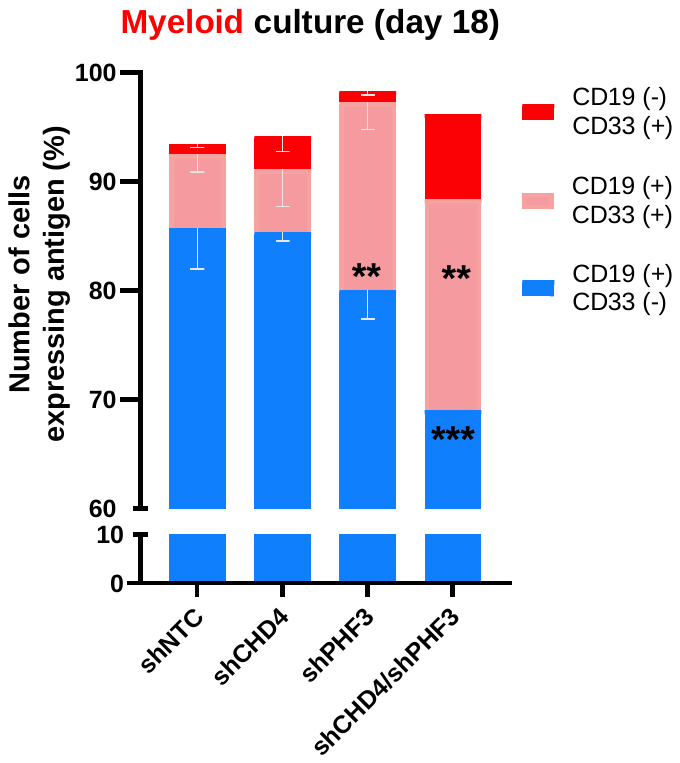

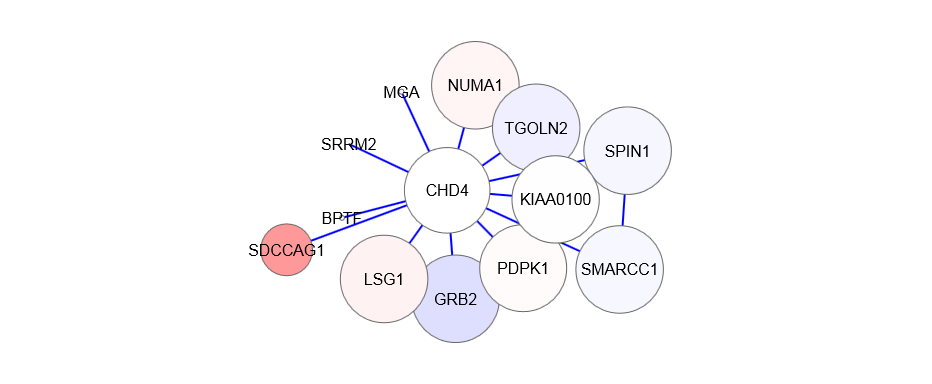

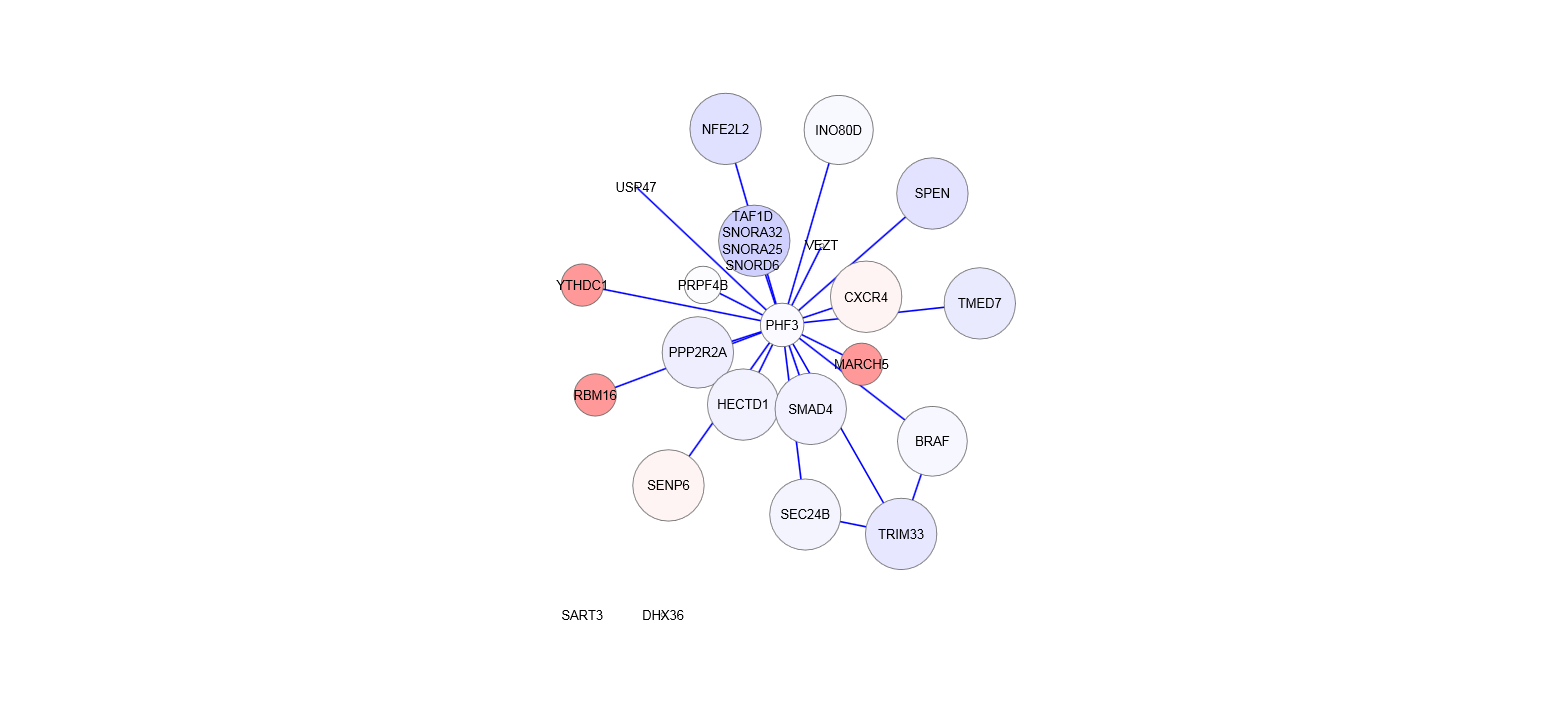

adjusted p (-logP)

log2 fold change

-11	0	6

I

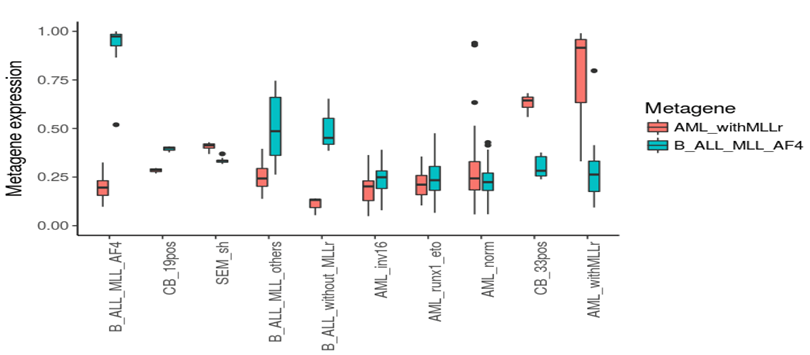

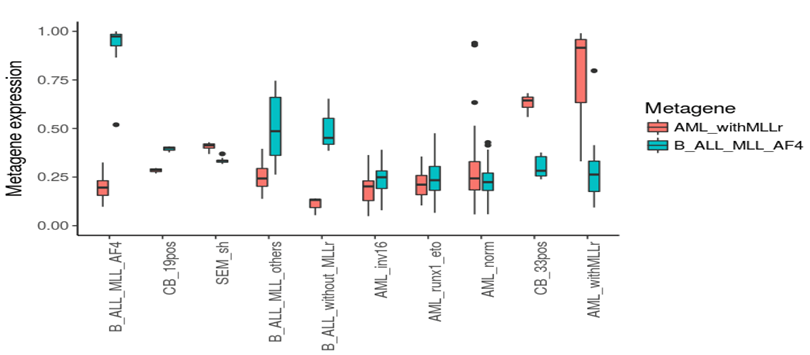

**ALL**

**AML**

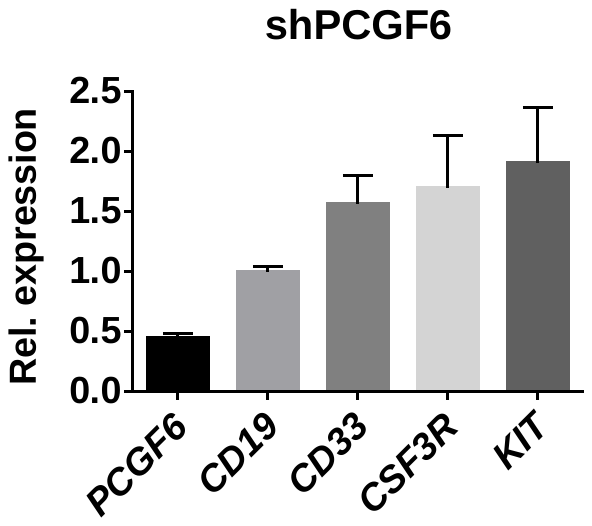

**shPCGF6**

H

**LS03 ALL (PDX)**

E

**CD19+CD33-**

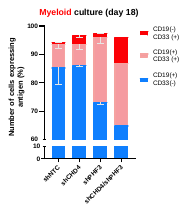

**CD19-CD33+**

**CD19+CD33+**

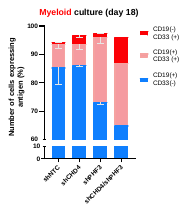

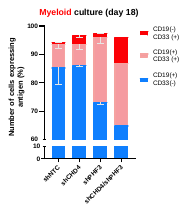

**Cell numbers (%)**

**LSALs**

**MPALs**

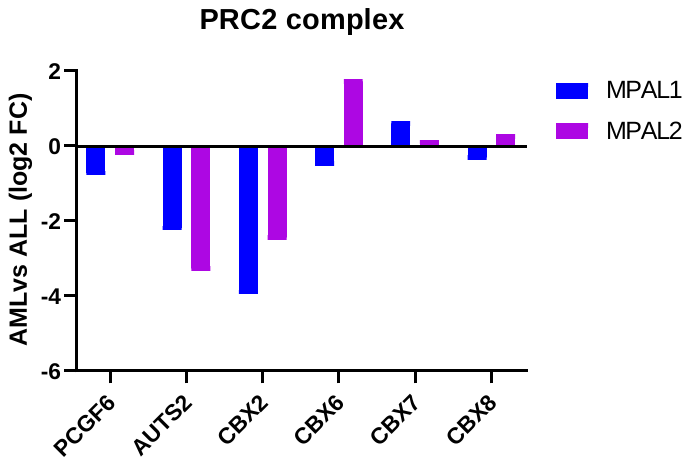

**Myeloid vs lymphoid (log2FC)**

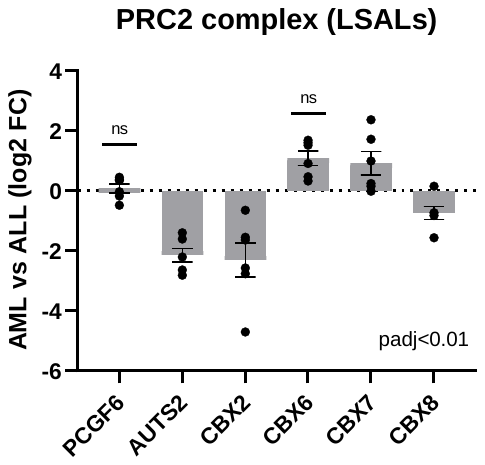

**AML vs ALL (log2FC)**

F

D

**FSC-A**

**697**

**REH**

**RS4;11**

**shNTC**

**shCHD4**

**shPHF3**

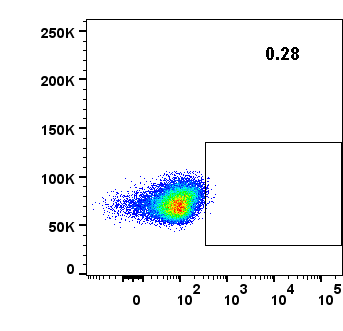

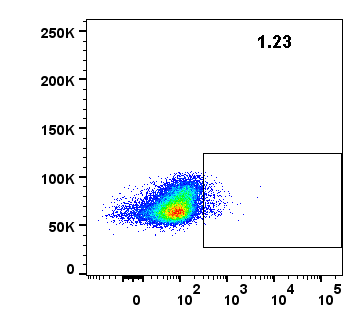

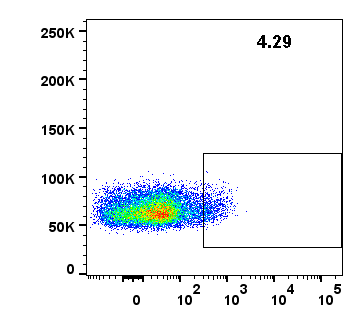

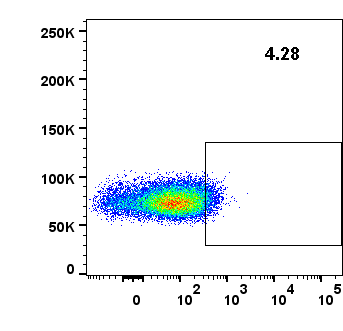

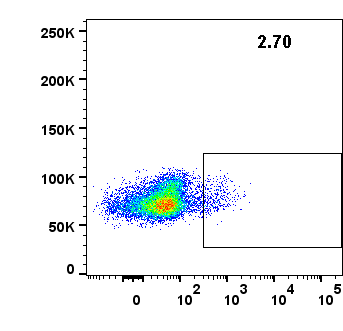

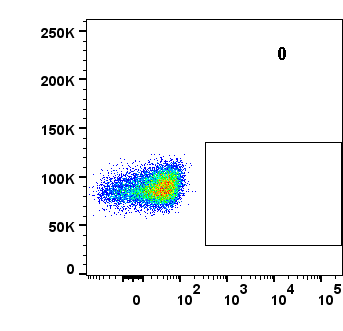

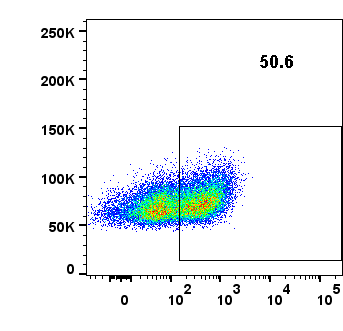

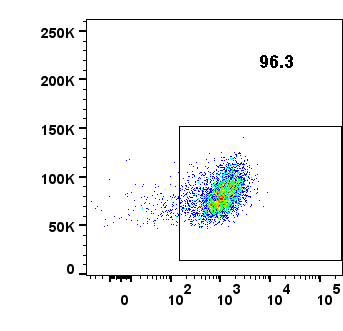

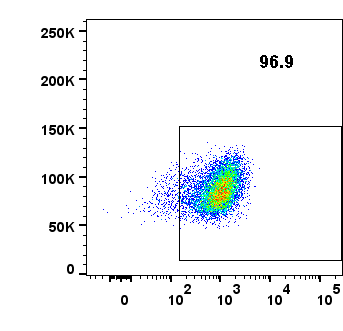

**CD33**

**0.28**

**4.28**

**0**

**2.70**

**4.29**

**1.23**

**50.6**

**96.3**

**96.9**

B

**PHF3**

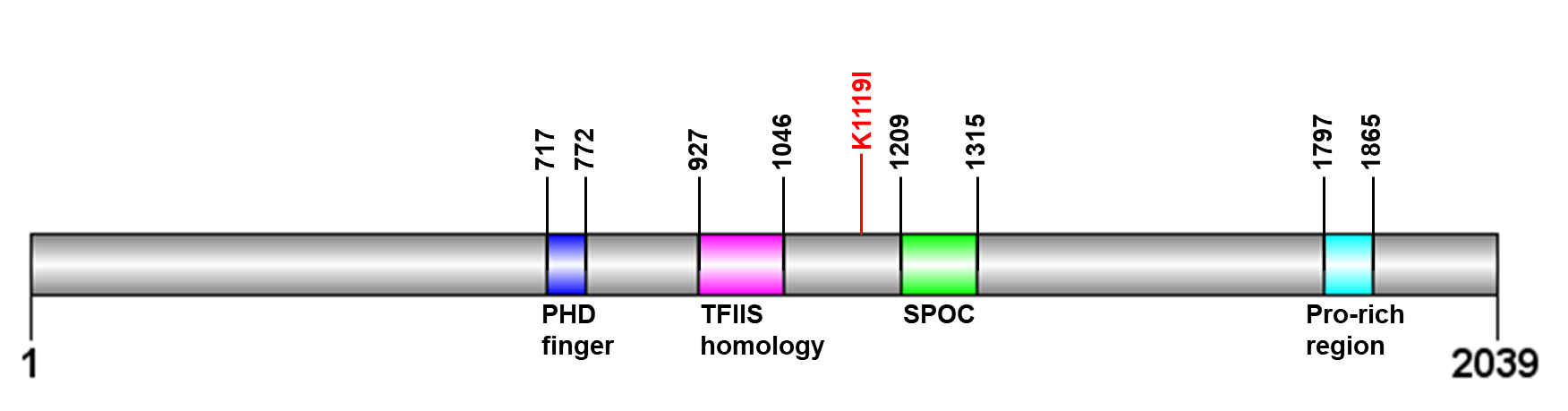

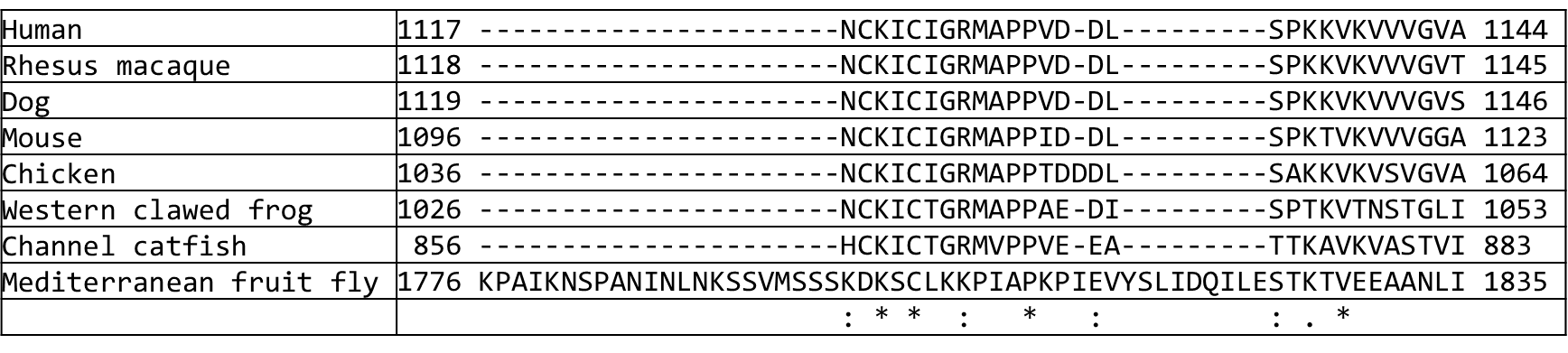

Figure S8

C

A

G
